# Aging activates latent enhancers that can be reversed with exercise

**DOI:** 10.64898/2026.11.02.763799

**Authors:** Jack Clarke, Fujian Wu, Vaibhao Janbandhu, Samuel Bagot, David Zheng, Pablo Mantilla Puccetti, Delfine Cheng, Xueqian Zhuang, Hoor E Maksura, Xuan Zhang, Alvaro Gonzalez Rajal, Julie Feng, Yen Tran, Alexander R Pinto, Peter K Koo, Charles D Cox, Richard P Harvey, Lee W Jones, Tuomas Tammela, Emily S Wong

**Affiliations:** Victor Chang Cardiac Research Institute, Sydney, NSW, Australia; School of Clinical Medicine, UNSW Sydney, NSW, Australia; Simons Center for Quantitative Biology, Cold Spring Harbor Laboratory, Cold Spring Harbor, NY, USA; Cancer Biology and Genetics Program, Sloan Kettering Institute, Memorial Sloan Kettering Cancer Center, New York, NY, USA; School of Biotechnology and Biomolecular Science, UNSW Sydney, NSW, Australia; Baker Heart and Diabetes Research Institute, Melbourne, VIC, Australia; School of Agriculture, Biomedicine and Environment, Baker Department of Cardiovascular Research, Translation and Implementation, La Trobe University, Melbourne, Victoria, Australia; Baker Department of Cardiometabolic Health, University of Melbourne, Parkville VIC, Australia; School of Biomedical Sciences, UNSW Sydney, NSW, Australia; Department of Population Sciences, Beckman Research Institute of City of Hope, Los Angeles, CA, USA

## Abstract

Aging profoundly alters the genome’s regulatory landscape, but the properties of age-acquired enhancers, including how they are selected and sustained in aged animals are unclear. We used single-cell multiomics in the mouse heart to identify age-responsive cis-regulatory elements and endurance exercise to understand which of these elements can be attenuated. Aging increased AP-1 engagement at more than 100,000 regulatory loci across cardiac cell types. Exercise opposed most of these gains. Despite low endogenous accessibility, age-gained enhancer sequences drove reporter activity comparable to stable elements, revealing substantial latent transcriptional potential. Local sequence features distinguished responsiveness to aging and exercise. Reporter perturbations implicated sequence outside the AP-1 core, and mutagenesis localized informative positions to dispersed segments of dynamic enhancers. These enhancers were IL-1β-sensitive and were preferentially responsive to neutralization of the cytokine in vivo. AP-1 inhibition further opposed age-associated transcription. Together, these findings support a model in which aging engages latent cell-type specific enhancers that are sustained in an active state by ongoing trans signaling. Aging therefore changes the deployment of existing regulatory information, while the acquired activity remains responsive to changes in signaling even in old age.

## INTRODUCTION

Aging alters cellular regulatory states, reshaping chromatin accessibility and gene expression across tissues^1,2^. Detailed molecular atlases have catalogued regulatory programs associated with aging, but do not establish whether those states are fixed or remain responsive to intervention. How enhancers are recruited to aged tissues and sustained in vivo is unknown. This distinction is particularly relevant in the heart, where cardiomyocytes are predominantly post-mitotic and long-lived, undergoing limited renewal^3^. Cardiac aging is accompanied by hypertrophy, fibrosis and impaired ventricular relaxation^1,4,5^. Determining which enhancers are engaged, and how they are maintained is central to understanding the mechanisms that enable, and constrain, an aged organ’s response to intervention.

Extracellular stimuli can activate previously unmarked regulatory elements, namely enhancers, expanding the repertoire of engaged elements^6^. Chronic, low-grade inflammation is a hallmark of ageing^7,8^. In the myocardium, IL-1 signalling engages inflammation pathways that promote fibroblast activation and extracellular matrix (ECM) remodelling^8,9^. JUN–FOS AP-1 complexes connect MAPK signalling to distal regulatory elements, where they cooperate with lineage-associated transcription factors (TFs)^9,10^. AP-1-associated chromatin opening recurs across multiple aging cell types^11^, and depletion of c-JUN or ectopic expression of FOXM1 can modify senescence-associated transcription and accessibility in aged cultured fibroblasts^12,13^, demonstrating that age-associated regulatory states remain malleable in vitro. In human cardiac fibroblasts, macrophage-derived IL-1β can activate profibrotic enhancer programs during heart failure^14^, suggesting that a similar mechanism is at play during aging.

Yet, whether inflammatory signalling selects and sustains age-recruited enhancers in aged animals is unknown. The role of the DNA sequence underlying age-acquired chromatin accessibility in driving enhancer activity has not been deciphered, and whether age-acquired enhancers can respond to physiological stimuli is also poorly understood.

Here, we combined single-nucleus multiomic profiling with enhancer reporter assays, sequence perturbations and targeted interventions in aged mice, including short and long-term endurance exercise. Endurance exercise modifies cardiovascular aging and reshapes systemic and tissue signalling^15–17^, providing a powerful, natural perturbation to identify responsive age-acquired regulatory elements. We identify an age-gained AP-1 enhancer program with substantial sequence-encoded transcriptional potential despite relatively weak endogenous chromatin features. We then investigated how local sequence features distinguish these elements and ask whether inflammatory signaling contributes to their engagement and maintenance in old hearts.

## RESULTS

### Exercise shifts aged hearts toward a younger regulatory state

We used endurance exercise to ask whether the regulatory state of an aged heart remains responsive to a physiological intervention. We studied four sex-balanced groups of C57BL/6J mice: young sedentary (YSed), young exercised (YEx), aged sedentary (ASed) and aged exercised (AEx) (**Figure 1A**). Exercised mice completed six months of treadmill training, beginning at 5 or 13 months and ending at 11 or 19 months of age, respectively. This long-term cohort tested how sustained exercise altered the development of age-associated states. Separate short-term training cohorts and pathway perturbations in aged mice tested responsiveness to interventions initiated later in life.

**Figure 1.**
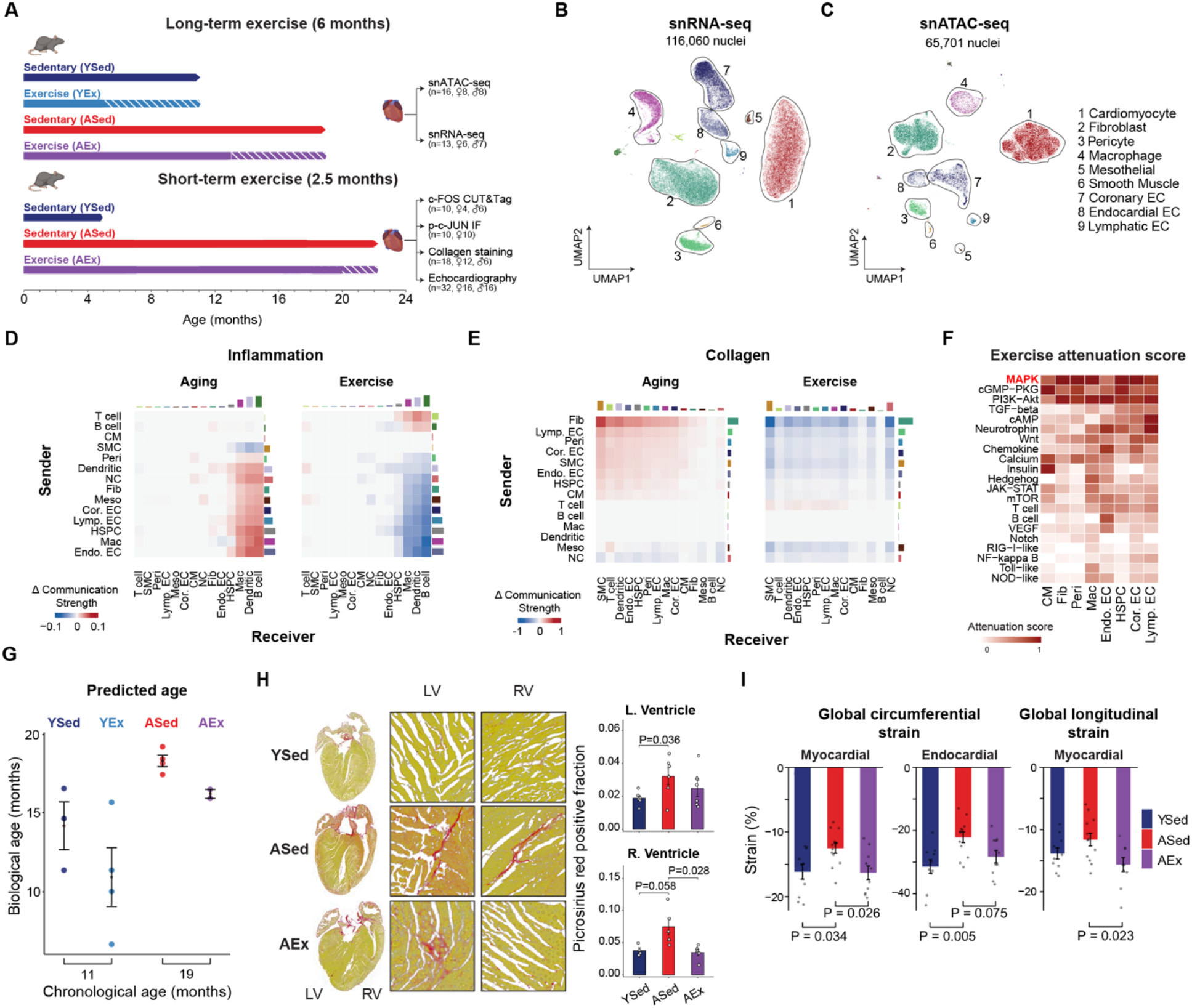
Endurance exercise opposes molecular and functional features of cardiac aging. **(A)** Experimental design for the long-term endurance-exercise and short-term exercise cohorts used in the study (**Methods**). **(B)** UMAP embeddings of snRNA-seq nuclei (n = 116,060). **(C)** UMAP embedding of snATAC-seq nuclei (n = 65,701). **(D–E)** Ligand–receptor analyses show changes in CellChat cell–cell communication strength for **(D)** inflammatory and **(E)** collagen GO pathway genes calculated from summed interaction probabilities. Bar graphs show total magnitude of communication change for each cell type. **(F)** Exercise attenuation scores calculated across KEGG pathways. **(G)** Predicted vs chronological age from a transcriptomic aging clock trained on *Tabula Muris Senis*^21,22^ and applied to pseudobulk profiles grouped by condition. Points are individual animals; error bars show SEM. **(H)** Picrosirius red staining across cardiac regions in a short-term exercise cohort. Points show individual animals, bars show animal-level means. P values were calculated using Wilcoxon rank-sum test on per-animal means. **(I)** Echocardiography-derived myocardial global circumferential strain (GCS), endocardial GCS, and myocardial global longitudinal strain (GLS) in the short-term exercise cohort (n = 10–12 animals per condition). Bars represent mean ± standard error of the mean; sex-adjusted Tukey HSD.

Single-nucleus RNA-seq (snRNA-seq) and ATAC-seq (snATAC-seq) resolved transcriptional and chromatin accessibility differences across the major cardiac cell types (**Figure 1B–C**). After quality control, we retained more than 65,000 nuclei for ATAC-seq and 116,000 for RNA-seq (**Figure S1–S3; Table S1– S4**). We assessed age-associated changes by comparing ASed with YSed hearts, and exercise responses in aged animals by comparing AEx with ASed hearts.

Exercise predominantly opposed age-associated transcriptional changes. Of 176 stringently defined age-associated genes identified using mixed-effects age-by-exercise models, 167 (94.9%) showed opposing expression changes with exercise. A similar directional trend was evident across all seven major cardiac cell types (**Figure S3K, S4–S5; Supplementary Results**). The most prominent transcriptional responses included inflammatory and ECM-remodelling programs, which were increased in aged sedentary hearts, consistent with established features of cardiac aging^1,2,18,8^. Exercise opposed these changes at both the transcriptional and chromatin-accessibility levels (**Figure S3J; Table S5**). Predicted ligand–receptor interactions further implicated altered intercellular signaling in inflammation and collagen programs (**Figure 1D–E; Table S6**). MAPK pathway signatures likewise increased with age and were opposed by exercise (**Figure 1F**), consistent with the role of MAPK signaling in inflammation^19^ and ECM remodeling^20^.

Exercise also moved cardiac gene expression toward a younger transcriptomic state. A cardiac age clock trained on independent *Tabula Muris Senis* RNA-seq profiles^21,22^ achieved an R² of 0.61 and a mean absolute error of 2.8 months in held-out samples (Figure **1G****, S6A–C; Table S7**). Applied independently to our dataset, the clock assigned exercised hearts a significantly lower transcriptomic age than sedentary hearts at both time points: by 2.1 months at the 19-month endpoint and 3.2 months at the 11-month endpoint.

In independent aged cohorts, short-term training attenuated collagen accumulation (**Figure 1H; Table S8**) and improved cardiac function by echocardiographic strain analysis (**Figure 1I, S6D; Table S9**), consistent with established benefits of exercise^23,24^. Exercise therefore shifted cardiac transcription toward a younger state and improved structural and functional features of aged hearts. We next sought to identity the regulatory elements whose activity remained responsive within those hearts.

### Aging engages an exercise-responsive AP-1 enhancer program

We identified aged-associated regulatory elements most responsive to exercise in aged hearts and annotated these elements based on inferred TF occupancy using TF footprinting (**Methods**). Across TFs, exercise opposed 55.3% of significant age-associated occupancy changes, and AP-1 stood out as the family whose age-associated gains in inferred occupancy were the most consistently attenuated (**Figure 2A; Table S10**). Across AP-1 motifs, footprint scores increased with age and decreased with exercise in every cell type, with concordant responses among individual animals (**Figure 2B–C, S7A–B**). Across cell types, aging increased inferred AP-1 occupancy at more than 100,000 loci. NF-κB and STAT motifs showed the same age-up, exercise-down pattern, placing AP-1 within a broader exercise-responsive inflammatory program (**Figure S7C**).

**Figure 2.**
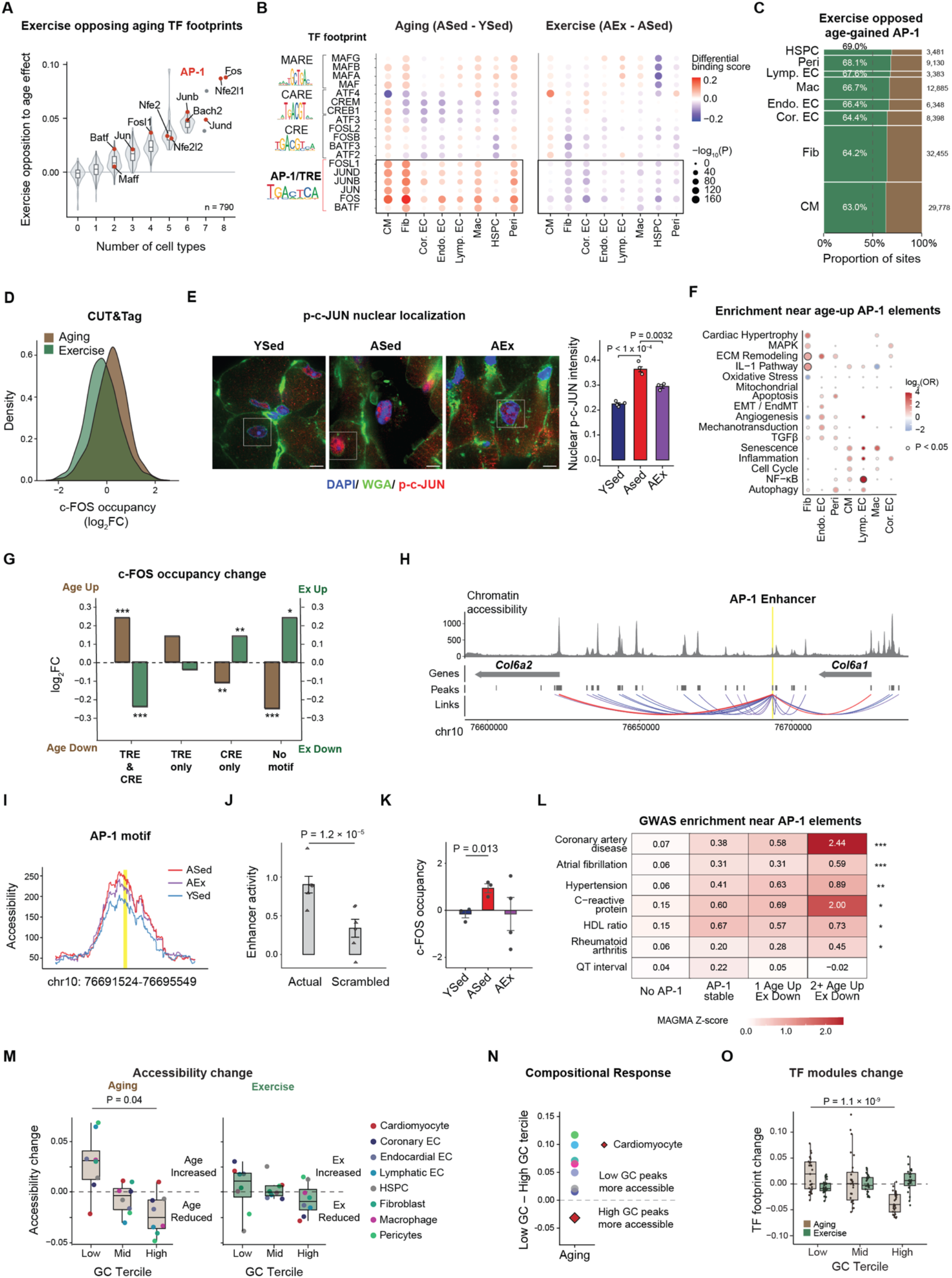
Aging engages an exercise-responsive AP-1 enhancer program. **(A)** Mean exercise-mediated opposition to aging for each TF footprint, grouped by the number of cell types exhibiting opposing age and exercise effects where both |Δ| > 0.02. Exercise opposition to aging calculated as the harmonic mean of age and exercise and averaged. **(B)** Dot plot showing AP-1 subfamily-specific differential TOBIAS scores associated with aging and exercise. **(C)** Proportion of inferred age-up AP-1 sites attenuated (green) or not attenuated (brown) by exercise across cell types. Bar height indicates the relative abundance of age-up AP-1 sites in each cell type with exact n annotated. Dashed line, 50%. **(D)** Density distributions of ageing-associated (ASed − YSed) and exercise-associated (AEx − ASed) log_2_FC values for c-FOS CUT&Tag peaks. Fisher’s exact test; OR = 37, *P* = 1.5 × 10^−10^. **(E)** Representative cardiac immunofluorescence images: DAPI (blue), WGA (green), p-c-JUN (red); Scale bar, 5 μm (left). Mean nuclear p-c-JUN intensity, quantified within DAPI-defined nuclei (right). Bars show mean ± SEM of mouse-level values; dots represent mouse means (n = 3–4 animals per condition). *P* values from Welch’s t-test on mouse-level means. **(F)** Enrichment of curated gene sets among genes linked to age-up AP-1 sites relative to stable. Points show log₂(OR); size indicates −log_10_(P). Outlined dots denote Fisher’s exact tests *P* < 0.05. **(G)** Median log_2_FC in c-FOS occupancy across peak classes defined by TRE and CRE motif presence. \**P* < 0.05; \*\**P* < 0.01; \*\*\**P* < 0.001. **(H)** A representative AP-1 site at chr10:76691524-76695549 linked to the Col6 locus by co-accessibility. Aged sedentary fibroblast snATAC-seq accessibility shown. **(I)** Accessibility at AP-1-containing enhancer. **(J)** Reporter activity of the endogenous enhancer and full scrambled control. Points show single-library log₂(α) estimates; shapes indicate biological replicates. Error bars show ± SEM. Likelihood-ratio test. **(K)** c-FOS CUT&Tag signal log2 CPM at the Col6-linked enhancer in the short-term exercise cohort. Bars show mean ± SEM of biological replicates; YSed n = 3, ASed n = 3, AEx n = 4. P values are quasi-likelihood F-tests on per-animal counts. **(L)** Mean MAGMA gene-level Z-score near cardiomyocyte and fibroblast AP-1-containing elements for GWAS traits. Stars indicate significance based on competitive gene-set analysis. P < 0.05; P < 0.01; P < 0.001. **(M)** Boxplots show centred mean snATAC-seq accessibility changes across cell-type-specific GC-content terciles for age and exercise. Points represent each cell type. Wilcoxon signed-rank tests (low GC vs high GC). **(N)** GC-associated accessibility responsive calculated by low-GC tercile minus high-GC tercile across cell types for aging contrasts. **(O)** Mean age-and exercise-associated changes in TF footprinted occupancy across NMF modules (n = 80; 10 per cell type), stratified by GC-content terciles of assigned peaks. Boxplots show the median, interquartile range (IQR) and whiskers extending to 1.5 × IQR; points represent individual modules. Wilcoxon rank-sum test, low versus high GC.

Direct binding measurements supported age-associated recruitment of AP-1 to these elements and their attenuation with endurance exercise. We mapped the AP-1 subunit c-FOS using Cleavage Under Targets and Tagmentation (CUT&Tag) in an independent cohort of young and aged sedentary mice and aged mice exposed to short-term exercise (**Figure S7D–E; Table S11**). Among H3K27ac-marked loci, indicative of active promoters and enhancers, sites that gained c-FOS binding with age were strongly enriched among footprint-defined age-up AP-1 sites opposed by exercise (OR = 37, P = 1.5 × 10^−10^; **Figure 2D**). AP-1 occupancy increased with the number of AP-1 motifs within an element (ρ = 0.32, P = 1.7 × 10^−8^^5^; **Figure S7F**).

Changes in AP-1 engagement were accompanied by changes in its activation. Nuclear phospho-c-JUN increased with age and declined after exercise (**Figure 2E, S7G; Table S12**), consistent with the age-opposing exercise effect on MAPK-associated transcriptional programs (**Figure 1F**).

The shared AP-1 response was implemented through predominantly lineage-restricted regulatory elements. More than 70% of stable sites, defined as unchanged with age or exercise, and dynamic AP-1 sites were restricted to a single cell type (**Figure S8A–B; Table S13**). In fibroblasts, age-gained AP-1 sites were preferentially located near genes involved in IL-1 signaling and ECM remodeling (IL-1: OR = 3.4, P = 0.015; ECM: OR = 2.7, P = 0.013; **Figure 2F; Table S14**). Exercise reduced AP-1 footprint scores at these inflammatory and matrix-associated loci, including fibrillar collagen genes (*Col1a1*, *Col3a1*) and inflammatory mediators (*Ccl2*, *Cxcl1*, *Il1r1*) (**Figure S8C–E**; **Table S15**).

AP-1 redistribution also varied with local bZIP motif architecture. Canonical FOS–JUN complexes predominantly bind the 7-bp TRE (TPA-responsive element) motif, whereas related CREB/ATF proteins favor the 8-bp CRE (cyclic AMP-responsive element) motif (**Figure S7A**). TRE motifs occupied low-GC regions, whereas CRE motifs occupied high-GC elements. With aging, c-FOS binding increased at TRE sites and decreased at CRE-only sites; exercise counteracted both trends, reducing occupancy at TREs and increasing it at CREs (aging P = 1.7 × 10^−10^; exercise P = 3.5 × 10^−6^; **Figure 2G, S8F**). Thus, aging and exercise redistributed c-FOS selectively between closely related sequence contexts.

Genes linked to distal AP-1 elements were more likely to be differentially expressed, particularly with age (**Figure S9A**). At a representative fibroblast locus associated with cardiac fibrosis, a distal AP-1 enhancer linked to *Col6a1* and *Col6a2* gained accessibility with age and lost accessibility with exercise, mirroring the expression of the linked genes (**Figure 2H–K**).

Genes near age-gained, exercise-opposed AP-1 elements were also enriched for human genetic associations with coronary artery disease, atrial fibrillation and hypertension (**Figure 2L**). Enrichment was strongest for genes near elements containing multiple AP-1 motifs. These associations connect the exercise-responsive enhancer program to genes implicated in human cardiovascular disease and suggest that exercise could mitigate disease-relevant regulatory activity via enhancers modulating expression changes at these specific genes.

This shared AP-1 response occurred within distinct genome-wide changes in chromatin accessibility. In non-cardiomyocyte lineages, AT-rich (low-GC) elements gained accessibility with age, whereas GC-rich elements lost it (**Figure 2M, S9B**), a relationship reproduced in an independent aging cohort^25^ (**Figure S9C**). Cardiomyocytes showed the opposite trend, with age-associated gains concentrated at GC-rich elements (**Figure 2N**), yet shared the age-associated increase in AP-1 engagement and its opposition by exercise. The GC-associated accessibility pattern remained evident in trained aged hearts (**Figure 2M**).

This GC-compositional pattern remained after grouping elements into functional modules with shared TF-footprint profiles: low-GC modules gained inferred occupancy with age and high-GC modules lost it, even after adjusting for promoter distance (**Figure 2O, S9D–E; Tables S16–18**). This relationship extended across diverse TF families (**Figure S9E**). Exercise opposed 23.2% of the age-associated occupancy gain at low-GC relative to high-GC modules, while a per-animal accessibility score contrasting these modules separated aged from young hearts and was unchanged by training (**Figure S9F**). Likewise, the exercise-responsive AP-1 footprints did not identify peaks with preferential restoration of age-associated accessibility. Regulatory engagement therefore remained responsive even where the broader accessibility profile persisted.

Together, these findings identify an age-gained AP-1 enhancer program that can be significantly reversed by exercise across cardiac lineages occurring within a broader compositional shift in chromatin accessibility. We next asked what distinguishes the elements recruited with age from those occupied constitutively, and whether their native chromatin state reflects their intrinsic regulatory potential.

### Age-gained enhancers harbor latent transcriptional potential

Elements that gained AP-1 engagement with age showed lower endogenous accessibility than stably occupied elements (**Figure 3A, S10A, Table S19**). We asked whether this profile reflected limited transcriptional capacity encoded by their sequences. Using lenti-STARR-seq, a massively parallel reporter assay (MPRA), we tested 1,861 endogenous 170-bp sequences outside their original genomic loci in primary cardiac fibroblasts isolated from 5-month-old mice. The reporter library included matched controls in which either the core AP-1 motif or the full sequence was shuffled (**Figure 3B, S10B; Table S20**).

**Figure 3.**
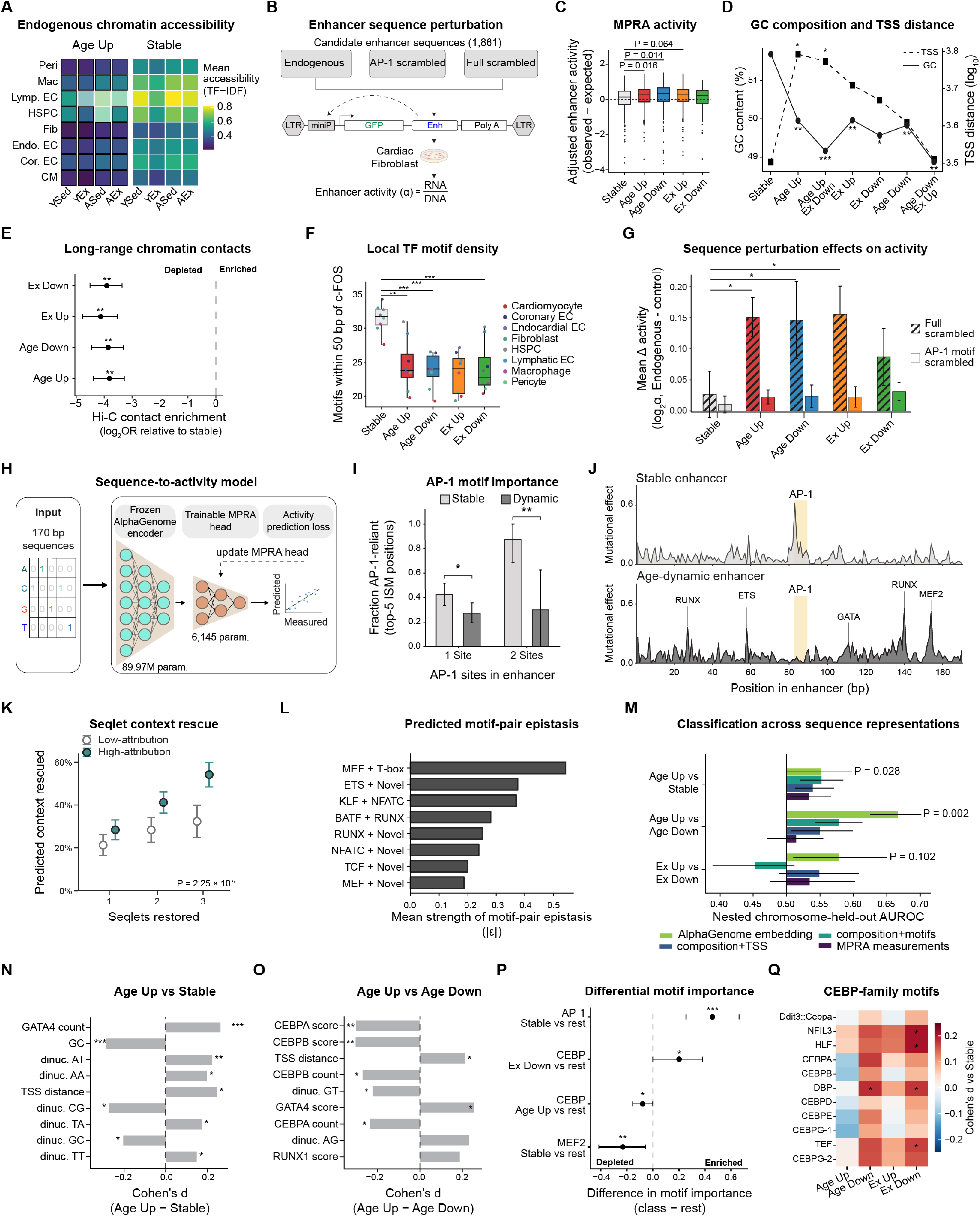
Age-dynamic AP-1 enhancers depend on distributed native sequence context. **(A)** Mean condition-specific TF–IDF-normalized snATAC-seq accessibility at c-FOS-containing peaks, stratified as age-up or stable. Significant differences between age-up and stable peaks are outlined black; Wilcoxon rank-sum test. **(B)** Lentiviral STARR-seq design testing 1,861 endogenous 170-bp AP-1-bound sequences, matched AP-1 motif-scrambled controls and fully scrambled controls downstream of a minimal promoter. **(C)** GC-and TSS-distance-adjusted MPRA activity comparison between dynamic and stable sequences. Wilcoxon rank-sum tests. **(D)** GC content (solid) and TSS distance (dashed) of MPRA sequences across sequence classes. Error bars represent 95% CI. Wilcoxon rank-sum tests versus stable: *FDR < 0.05, **FDR < 0.01, ***FDR < 0.001. **(E)** Enrichment of dynamic c-FOS-bound regions at cardiomyocyte Hi-C anchors, relative to stable c-FOS sites. Points show log₂(OR); Fisher’s exact tests. **(F)** Number of TF motifs bound within 50 bp of a bound c-FOS motif, stratified by binding stability and cell type. Wilcoxon rank-sum tests; \*\**P* < 0.01; \*\*\**P* < 0.001. **(G)** Mean MPRA activity change between endogenous and scrambled sequences. Error bars show mean ± SEM. **(H)** Schematic of MPRA model training^27^. **(I)** AP-1 reliance defined by AP-1 overlap among the five highest-importance positions following in silico saturation mutagenesis (ISM) of enhancers (150 stable and 160 dynamic) stratified by AP-1-site count. Bars show proportions and bootstrapped 95% CI (5,000 resamples); Fisher’s exact test *P < 0.05, **P < 0.01. **(J)** Endogenous sequence attribution maps for a stable and an age-down/exercise-up enhancer. Track height shows signed contribution to the chromosome-held-out non-AP-1 context model. Motifs with high attributions are annotated. **(K)** Composition-preserving in silico rescue across 91 enhancers. Fraction of the model-predicted non-AP-1 context effect recovered by restoring one, two, or three high-attribution 12-bp seqlets or matched low-attribution regions within fully scrambled sequences. Points show means and error bars show bootstrapped 95% CI. **(L)** Strongest predicted epistatic interactions identified within high attribution seqlets and matched to TF motifs. **(M)** Nested chromosome-held-out AUROC for four feature representations across pairwise contrasts. Error bars show 95% chromosome-bootstrap intervals. Empirical P values, based on 499 within-chromosome permutations. **(N-O)** The nine most strongly associated sequence features in the age-up vs stable **(N)** and age-up vs age-down **(O)** contrast. Two-sided feature-association tests: *FDR < 0.05, **FDR < 0.01, ***FDR < 0.001. **(P)** Forest plot of class-specific motif reliance inferred by ISM. Points and bootstrapped-derived 95% CIs show motif-count-matched differences in top 20 ISM positions overlapping each motif footprint; Wilcoxon rank-sum test; *FDR < 0.05, **FDR < 0.01. **(Q)** Cohen’s d for individual C/EBP-related motif scores relative to stable across sequence classes. Wilcoxon rank-sum *FDR < 0.05. Motifs annotated with JASPAR 2024 CORE.

Despite their weaker endogenous chromatin signatures, age-gained sequences drove reporter activity comparable to stable sequences (**Figure 3C, S10C–E**). Age-gained elements had lower GC content and greater distance from promoters than stable elements (**Figure 3D, S10F–I**). Dynamic elements also showed lower enrichment at cardiomyocyte Hi-C anchors and a sparser local landscape of other TF motifs than stable elements (**Figure 3E–F, S10J–K**).

The contrast between weak native accessibility and substantial reporter activity reveals latent transcriptional potential in these age-gained enhancer sequences. Their DNA can support strong activity when tested outside its original genomic setting, consistent with endogenous context regulating when that potential is deployed. We next asked where this competence resides within the enhancer sequence.

### Sequence outside the AP-1 core supports dynamic enhancer activity

The strong reporter activity of age-gained enhancers prompted us to ask whether their potential resided primarily in the AP-1 motif or in the surrounding sequence. Scrambling the AP-1 core had relatively minor effects across regulatory classes. Comparing the AP-1-core and full-sequence scrambling controls indicated a greater contribution from non-core sequence in dynamic than stable elements (**Figure 3G**; 0.128 versus 0.016; Cohen’s d = 0.144, FDR = 0.034). Thus, the contribution of sequence outside the AP-1 core distinguished these classes more clearly than dependence on the core motif.

We next used sequence modeling to locate the bases contributing to enhancer activity. A regression head trained on frozen AlphaGenome representations of the assayed sequences (**Figure 3H**)^26,27^ achieved a Pearson correlation of 0.73 on held-out sequences. In silico mutagenesis (ISM) placed the most consequential substitutions close to the AP-1 site in stable elements but across dispersed positions in dynamic enhancers (**Figure 3I-J**).

Individual dynamic enhancers contained multiple informative 12-bp segments, termed ‘seqlets’, outside the AP-1 core that contributed to predicted activity. Restoring these high-impact segments to shuffled constructs in silico progressively recovered predicted activity; a subset of these segments corresponded to known TF-binding motifs (**Figure 3K**).

Motif analyses identified candidate TF inputs into this distributed architecture. Dynamic elements were enriched for motifs annotated to MEF2C, TCF7L2 and GATA4, whereas stable elements were enriched for NFATC3 and ETV4 motifs (FDR ≤ 0.04). ISM-based epistasis analysis further predicted locus-specific, non-additive contributions from motif-containing regions (**Figure 3L; Supplementary Results**).

Having identified sequence contributions to reporter activity, we asked whether local sequence also distinguished the direction of enhancer remodeling. Sequence-based models separated age-gained from age-decreased elements more readily than age-gained from stable elements. Models using AlphaGenome embeddings outperformed those using annotated sequence features or MPRA measurements (**Figure 3M**).

GC content and motif diversity distinguished age-gained from stable elements, whereas C/EBP motif enrichment was the strongest annotated feature separating age-gained from age-decreased elements (**Figure 3N–O**). C/EBP motif scores were highest in elements that lost AP-1 with age, and this association persisted after accounting for AP-1 motif count, GC content and promoter proximity. A related C/EBP– PAR-bZIP signature, including DBP-related features, was also enriched among elements that lost AP-1 engagement with exercise (**Figure 3P–Q, S10L**).

This sequence signature also distinguished exercise responsiveness within the age-gained class. A composite motif score integrating C/EBP, PAR-bZIP and ATF4 features was elevated in age-gained elements whose AP-1 engagement decreased with exercise (*P* = 0.003). Local sequence was therefore associated with both the direction of age-related AP-1 remodeling and subsequent responsiveness to exercise.

Together, reporter perturbations established a contribution from sequence outside the AP-1 core, and modeling localized predictive information to dispersed segments of dynamic enhancers. These results identify a distributed cis-regulatory basis for enhancer potential and associate local sequence with age-and exercise-related responsiveness. We next asked which signals engage this potential in the aging heart.

### IL-1β preferentially recruits AP-1 to age-gained enhancers

The enrichment of IL-1 signaling genes near age-gained AP-1 elements suggested an inflammatory input into this enhancer program. We tested this directly by stimulating primary cardiac fibroblasts with IL-1β and measuring c-FOS recruitment across enhancer classes defined in vivo (**Figure 4A, S11A-C**). Age-up elements responded most strongly, stable elements showed an intermediate response and age-down elements responded most weakly (P < 9 × 10^−8^; **Figure 4B, S11D; Table S21**). IL-1-responsive elements also showed greater age-associated c-FOS gain in vivo than unresponsive elements with comparable baseline chromatin accessibility (**Figure S11E**). Thus, IL-1β preferentially engaged the enhancer class that acquired AP-1 occupancy during cardiac aging.

**Figure 4.**
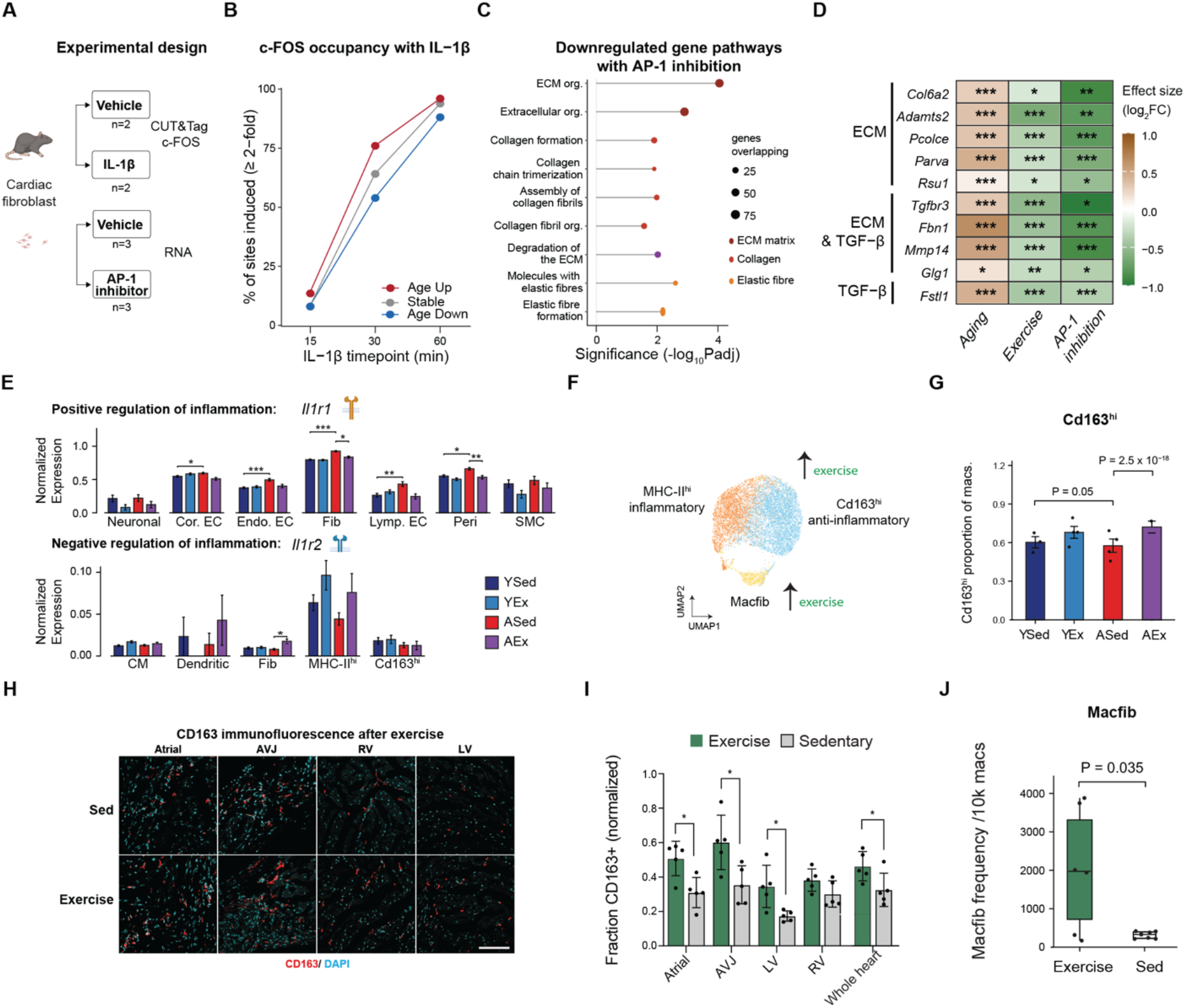
IL-1β selectively engages AP-1 to age-gained fibroblast enhancers. **(A)** Experimental design for IL-1β stimulation and AP-1 inhibition in mouse-derived cardiac fibroblasts. **(B)** Proportion of TOBIAS-defined fibroblast AP-1 sites with at least a two-fold increase in c-FOS CUT&Tag occupancy after IL-1β stimulation, stratified by AP-1 site class and time point. **(C)** Enrichment of terms in downregulated genes after 6 h AP-1 inhibitor treatment in cardiac fibroblasts (log_2_FC < −0.585 and P < 0.05). Point size denotes gene number. **(D)** Heatmap showing log2FC for genes associated with ECM and TGF-β-associated gene sets. Genes are grouped according to pathway annotation as ECM-only, ECM and TGF-β-associated, or TGF-β-associated. *FDR ≤ 0.05, **FDR ≤ 0.01, and ***FDR ≤ 0.001. **(E)** Normalized expression of *Il1r1* and *Il1r2* across cell types and conditions. Error bars show mean ± SEM across nuclei. MAST differential expression tests *FDR < 0.05; **FDR < 0.01; ***FDR < 0.001. **(F)** UMAP embedding of cardiac macrophages colored by annotated subpopulation. **(G)** Cd163^hi^ macrophage frequency calculated separately per animal; points represent animals; χ² test. **(H)** Representative immunofluorescence images showing CD163 (red) and DAPI (blue) staining across separate cardiac regions in sedentary (top) and 1-year exercised (bottom) mice. Scale bar, 100 μm. **(I)** Regional quantification of the proportion of CD163^+^ cells in exercised and sedentary animals across cardiac regions. Error bars show mean ± SEM; Welch’s t-test *P < 0.05 (Atria P = 0.0105, AVJ P = 0.0210, LV P = 0.0161, RV P = 0.1087). **(J)** Macfib frequency calculated per animal as the number of nuclei per 10,000 macrophage nuclei. Each point represents an animal. Boxplots show the median, IQR and 1.5 × IQR whiskers; Welch’s t-test.

In a complementary experiment, pharmacological AP-1 inhibition in the same cells reduced transcriptional programs associated with ECM organization, collagen biosynthesis and TGF-β signaling (**Figure 4C–D, S12A-E; Table S22**). These responses connect the signal-responsive AP-1 program to fibroblast matrix regulation.

We then examined whether exercise altered the cardiac signaling environment associated with this response. Exercise changed receptor expression associated with IL-1 responsiveness. Il1r1 increased with age in fibroblasts, pericytes and vascular lineages, and decreased with exercise in fibroblasts and pericytes (**Figure 4E**). Conversely, exercise increased the IL-1 decoy receptor Il1r2, predominantly in inflammatory macrophages. These changes were consistent with reduced IL-1 responsiveness with exercise. Sparse detection of IL-1 ligands limited assessment of their expression changes (**Figure S12F–G**).

Exercise also remodeled the cardiac macrophage landscape. Cardiac macrophages partitioned into an inflammatory MHC-II^hi^ population, a Cd163^hi^ population with resident and remodeling signatures, and a rare ECM-expressing macfib state^28^ (**Figure 4F, S13; Tables S23–26**). Exercise reduced the inflammatory macrophage fraction and increased the Cd163^hi^ population in both young and aged hearts, a shift supported by tissue imaging in an independent cohort (**Figure 4G–I**). The macfib state expanded approximately sevenfold, from 2.9% of macrophages in sedentary hearts to 20.7% after exercise (**Figure 4J**). Human counterparts of the Cd163^hi^ and macfib states, but not the inflammatory population, were positively associated with left ventricular ejection fraction (**Figure S14; Table S27**), linking these exercise-associated states to cardiac function in humans.

IL-1β can therefore preferentially recruit AP-1 to age-gained enhancers, while exercise alters cellular states and receptor expression associated with inflammatory responsiveness. We next tested whether continued AP-1 activity and inflammatory input were required to sustain age-associated regulatory activity in vivo.

### Ongoing AP-1 activity sustains age-associated transcription

The reduction of AP-1 engagement after exercise suggested that ongoing AP-1 activity helps maintain the aged transcriptional state. To test this in aged tissue, we delivered AAV9 encoding dominant-negative AP-1 (dnAP-1) under the cardiomyocyte-restricted cTnT promoter, or a GFP control, to ∼23 month old mice and profiled their hearts five weeks later (**Figure 5A, S15; Table S28**). Co-expressed tdTomato confirmed myocardial transduction, with reporter signal covering 54.6 ± 3.3% of ventricular area (*n* = 4; **Figure S16A**).

**Figure 5.**
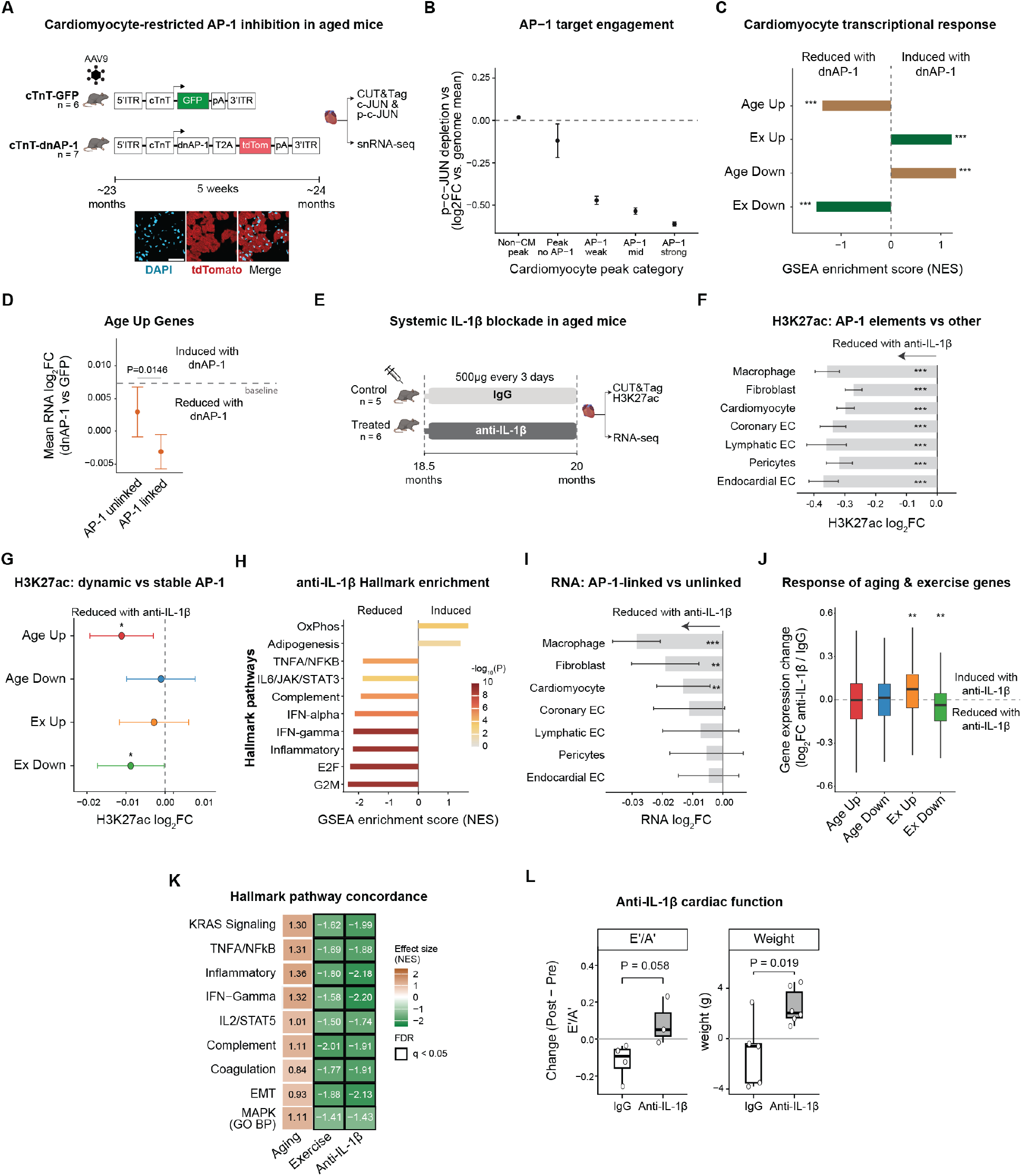
AP-1 inhibition and IL-1β neutralization interventions remodel age-associated enhancer activity in vivo. **(A)** Cardiomyocyte-restricted AP-1 inhibition: aged C57BL/6J mice received AAV9–cTnT–GFP (n = 6) or AAV9– cTnT–dnAP-1–T2A–tdTomato (n = 7). 5 weeks later, hearts were collected for c-JUN/p-c-JUN CUT&Tag (n = 4 dnAP-1, n = 3 GFP) and snRNA-seq (n = 3 dnAP-1, n = 2 GFP) analyses. **(B)** Spike-in-normalized peak-level dnAP-1-versus-GFP log_2_FC centered on the genome-wide mean. Zero denotes the mean treatment response. Low-high activity defined by tercile of inferred AP-1 binding strength. Error bars represent 95% CI. **(C)** Gene set enrichment analysis (GSEA) normalized enrichment scores (NES) for cardiomyocyte age- and exercise-associated gene sets in the dnAP-1 transcriptional response; positive and negative NES indicate enrichment among dnAP-1-up- and down-regulated genes, respectively. ***P < 0.001. **(D)** Mean dnAP-1 snRNA-seq log_2_FC for ageing-regulated cardiomyocyte genes grouped by proximity to a footprint-defined cardiomyocyte AP-1 peak. Points show means ± 95% CI; Wilcoxon rank-sum test. **(E)** Experimental design for IL-1β neutralization. Aged C57BL/6J mice received control IgG (n = 5) or anti-IL-1β antibody (n = 6) for 5 weeks, with echocardiography performed at baseline and endpoint; subsequent H3K27ac CUT&Tag and bulk RNA-seq. **(F)** Association between anti-IL-1β induced changes in H3K27ac signal at peaks with and without an overlapping AP-1 footprint; adjusted for baseline signal and log peak width. ***P < 0.001. **(G)** H3K27ac response to anti-IL-1β neutralization at dynamic AP-1 sites versus stable sites. Points show the difference in anti-IL-1β-associated H3K27ac change between dynamic and stable sites; negative values indicate a greater loss at dynamic sites with 95% CI shown. **(H)** Hallmark GSEA of genes ranked by anti-IL-1β associated RNA log_2_FC. **(I)** Association between anti-IL-1β induced changes in gene expression in AP-1-proximal gene (within 100 kb of the TSS) vs non proximal genes. Linear-model coefficients adjusted for baseline expression and 95% CIs are shown. **P < 0.01, ***P < 0.001. **(J)** Anti-IL-1β induced RNA log_2_FC among the top 250 age-and exercise-responsive gene sets identified from all-cell pseudobulk analyses and ranked by significance followed by fold change. Boxes show median and IQR; whiskers extend to 1.5 × IQR. Directional gene-set enrichment assessed using CAMERA. **P < 0.01. **(K)** GSEA of MSigDB Hallmark gene sets across anti-IL-1β, Aging and exercise contrasts. Selected terms are shown. **(L)** Changes in echocardiographic and physiological measurements, calculated as end-of-study minus baseline values, comparing IgG and anti-IL-1β groups. Points represent individual animals, boxplots show median, IQR and 1.5 × IQR whiskers; Welch’s t-test nominal P values are shown. E′/A′, early-to-late diastolic tissue Doppler velocity ratio.

dnAP-1 reduced phospho-c-JUN occupancy at cardiomyocyte-associated AP-1 elements, with the largest reductions at sites with the strongest AP-1 footprints (**Figure 5B**). H3K27ac also decreased at overlapping AP-1 sites (**Figure S16B**). These responses did not differ detectably between age-gained and age-lost elements (*P* = 0.69). Non-cardiomyocyte-associated regulatory elements showed little change, consistent with cardiomyocyte-restricted dnAP-1 expression.

Single-nucleus RNA-seq showed that cardiomyocyte genes downregulated by dnAP-1 were enriched for age-up and exercise-down gene sets (*n* = 3 dnAP-1, *n* = 2 GFP; **Figure 5C, S16C–D; Table S29**). The age-opposing response was strongest for genes near AP-1 sites (*P* = 0.01; **Figure 5D**). The dominant transcriptional responses involved fatty acid metabolism (FDR = 0.005) and oxidative phosphorylation (FDR = 4 × 10^−4^; **Figure S16E**). For oxidative phosphorylation, the response matched the direction of exercise and opposed the age-associated change. Fibroblasts also reduced expression of ECM-associated genes (**Figure S16F**), suggesting a non-cell-autonomous response to cardiomyocyte AP-1 inhibition.

Continued AP-1 activity therefore contributes to the maintenance of age-associated cardiac transcription. Because direct AP-1 inhibition affected both age-gained and age-lost elements, we next asked whether removing an upstream signal could preferentially attenuate the enhancer program acquired with age. IL-1β provided such a candidate because it preferentially recruited c-FOS to age-gained elements in cardiac fibroblasts.

### IL-1β sustains age-gained enhancer activity in old hearts

IL-1β stimulation established that age-gained enhancers were preferentially responsive to an inflammatory input. We next asked whether continued IL-1β signaling was required to sustain their activity in an already aged heart. Aged mice received an IL-1β-neutralizing antibody or control IgG for five weeks, followed by bulk-heart RNA-seq and H3K27ac CUT&Tag (**Figure 5E, S17A–D**).

IL-1β neutralization reduced H3K27ac more strongly at distal AP-1 elements than at non-AP-1 regions across cell-type-annotated regulatory sets (**Figure 5F, S17E**). Within AP-1 elements defined by their age- and exercise-associated responses, H3K27ac loss was most pronounced in the age-gained and exercise-down classes, whereas stable elements were less affected (**Figure 5G**). Thus, withdrawing an inflammatory input preferentially attenuated acetylation at the enhancer classes engaged by aging and opposed by exercise.

Neutralization also reduced known downstream targets of IL-1β signaling^29^, including IL-1β/NF-κB (P = 0.0087) and IL-6 pathway genes (P = 0.037; **Figure 5H**). Genes near AP-1 elements showed stronger transcriptional responses than genes without these associations (**Figure 5I**). The response was directionally concordant with exercise: exercise-induced genes generally increased after neutralization, whereas exercise-downregulated genes decreased (**Figure 5J**). Both interventions attenuated age-elevated signatures of TNFα–NF-κB signaling, interferon-γ response, complement, coagulation and epithelial–mesenchymal transition (**Figure 5K, S17F**). Metabolic transcriptional programs increased after neutralization (**Figure S17F**).

Over five weeks, these molecular responses were not accompanied by a statistically supported improvement in cardiac function. Among diastolic indices, the E′/A′ ratio showed a trend toward preservation (P = 0.058; **Figure 5L, S17G**), in the direction of previously reported improvements in diastolic function^30^. Treated animals gained body weight relative to controls (P = 0.019), consistent with the established role of IL-1 signaling in food intake and energy homeostasis in mice^31,32^.

An inflammatory signal that preferentially recruits AP-1 to age-gained enhancers therefore also contributes to maintaining their active state in old hearts. Together with their intrinsic transcriptional competence and exercise responsiveness, these findings support a model in which aging engages latent cardiac regulatory potential whose continued activity is sensitive to the signaling environment. Cis-regulatory architecture shapes that potential and its responsiveness, while ongoing trans signaling helps sustain the age-associated enhancer program.

## DISCUSSION

Aging is an interplay between cis-regulatory architecture and trans-acting signalling. Our findings support a model in which aging reshapes chromatin accessibility thereby exposing latent enhancer potential. These age-acquired enhancers are selectively acted upon and their activity maintained through ongoing inflammatory signaling. Age-gained AP-1 elements possess a distinct sequence structure and combine relatively weak endogenous chromatin features with substantial reporter activity, demonstrating that low accessibility need not imply weak regulatory capacity. Here, latent potential describes the transcriptional capacity encoded by sequences that are weakly engaged in their native context. Exercise and interventions initiated in aged animals attenuated components of the AP-1 enhancer and transcriptional programs. Aging therefore changes the deployment of existing regulatory information, while the acquired activity remains responsive to changes in signaling even within an old organ.

There is a central distinction between the transcriptional capacity of an enhancer sequence and the extent to which that capacity is engaged in its native setting. Despite lower accessibility, lower GC content and sparser motif neighborhoods, age-gained sequences drove reporter activity comparable to stable elements in cardiac fibroblasts from young mice. Their regulatory capacity is therefore greater than their endogenous chromatin features alone would suggest, consistent with context-dependent deployment. Experimental perturbations identify a distinguishing contribution from sequence outside the canonical AP-1 site. Sequence modeling then localized influential positions to dispersed segments in dynamic enhancers, and restoring those segments into shuffled constructs progressively recovered predicted activity. The results support a distributed regulatory architecture in which multiple sequence features contribute to enhancer output^33,34^; the model-based epistasis results further indicate that these contributions need not be additive. Aging acts at both the underlying chromatin context in which these sequences are embedded and on their upstream signals.

Enhancer architecture specifies where the changing trans environment is read. Sequence outside the AP-1 core carries information about both age-associated remodeling and responsiveness to intervention. C/EBP-associated features were enriched at elements that lost AP-1 with age, and related C/EBP–PAR-bZIP features marked age-gained elements whose engagement declined with exercise. Previous work has linked C/EBPβ to exercise-induced cardiac growth^35^.

Sequence-selective responses extend to closely related bZIP recognition sites. Aging shifted c-FOS occupancy toward TRE-containing elements and away from CRE-only elements, and exercise opposed both changes. Thus, increased AP-1 activation with age was accompanied by redistribution across regulatory contexts, while exercise altered both the amount and location of factor engagement. Together with the C/EBP-associated signature, this reciprocal response shows that a shared change in physiological state can have different consequences at closely related regulatory sequences.

The persistence of the age-gained program depends in part on continuing inflammatory input. IL-1β preferentially recruited c-FOS to fibroblast elements that gained AP-1 with age, and neutralization in aged mice preferentially reduced H3K27ac at age-gained and exercise-down AP-1 elements. These complementary interventions connect signal responsiveness to maintenance of the acquired regulatory state in vivo. The result extends the established role of IL-1β in profibrotic fibroblast regulation^14^ to an enhancer program engaged during aging. Exercise-associated changes in macrophage states and IL-1 receptor expression provide a plausible tissue context for this responsiveness.

Continued AP-1 activity also contributes to the transcriptional state in aged cardiomyocytes. Cardiomyocyte-restricted AP-1 inhibition reduced phospho-c-JUN occupancy and acetylation at AP-1 sites and opposed age-associated expression changes, particularly among genes near those elements. Direct AP-1 inhibition did not preferentially affect age-gained over age-lost elements, whereas IL-1β blockade selectively reduced acetylation at the age-gained and exercise-down classes. This difference highlights the role of cis-regulatory architecture beyond the AP-1 site for age-associated enhancer selection through upstream signaling.

The responsiveness of the AP-1 enhancer program did not require reversion of the broader age-acquired accessibility landscape. Exercise attenuated AP-1 engagement while the GC-associated redistribution of accessibility remained evident. Age-associated changes in heterochromatin and higher-order genome organization^36,37^ offer possible sources of persistent regulatory context, and models in which signaling and chromatin organization operate across different timescales^38^ provide a framework for interpreting this behavior. Our observations establish that a substantial regulatory attenuation can occur while the tissue retains broad chromatin features associated with age.

These findings connect the potential of regulatory DNA to the plasticity of age-acquired enhancer states. Cis-regulatory architecture shapes which elements are responsive, while ongoing trans signaling helps determine whether their potential is engaged. This dependence provides a basis for attenuating age-associated expression without erasing the broader GC-associated chromatin features acquired with age. Aging exposes regulatory potential at stimuli responsive enhancers whose activity remains responsive to intervention, leaving opportunities to modify gene regulation late in life.

## RESOURCE AVAILABILITY

### Lead contact

Requests for further information and resources should be directed to and will be fulfilled by the lead contact, Emily S Wong.

### Data and Code Availability

Raw sequencing data have been deposited in the NCBI Expression Omnibus (GEO) under accession numbers GSE342513 (snRNA-seq), GSE342511 (snATAC-seq), GSE342503 (c-FOS CUT&Tag), GSE342504 (IL-1β fibroblast CUT&Tag), GSE342508 (fibroblast bulk RNA-seq), GSE339085 (MPRA), GSE342506 (AAV9 CUT&Tag), GSE342509 (AAV9 snRNA-seq), GSE343166 (anti-IL-1β CUT&Tag), and GSE343167 (anti-IL-1β bulk RNA-seq). All code is available at GitHub https://github.com/ewonglab/Cardiac_AgeExercise. Publicly available datasets used in this study were obtained from the Gene Expression Omnibus under accession numbers GSE132040 for Tabula Muris Senis bulk RNA-seq, GSE281772 for mouse cardiac ageing, GSE213128 for human cardiac valve, and GSE226314 for human heart failure snRNA-seq.

## Acknowledgements

We thank Karthik Hemanthekumar, Disha Anand, Robert Shearer, Osvaldo Contreras, Qing Wang, Paola Cornejo, Veronika Petrova, and Wunna Kyaw for their support. We thank Mathias Francois and Robert Graham for manuscript feedback. We thank the Garvan Genomics Platform at the Garvan Institute of Medical Research for nuclei sorting, library preparation, and single-cell sequencing, and the Australian Genome Research Facility for additional sequencing services. We thank Yitong Zhao from the University of Technology Sydney for access to the ARC LIEF funded National Live Cell Analytics Facility for Organelles’ Interactome Discovery (LE250100063). We also thank Nana Sunn, Hardik Doshi, Paul Young and David Humphreys of the Victor Chang Cardiac Research Institute Innovation Centre for their technical assistance. This work was supported by a Snow Medical Fellowship and NHMRC Investigator Grant (RG211011) to E.W., and an Australian Research Training Program scholarship to J.C. Experiments were performed on the computational facility of the National Computational Infrastructure (NCI) through the UNSW HPC Scheme (DOI: 10.26190/PMN5-7J50).

## Author Contributions

F.W., D.C., X.Z., V.J., D.Z., Y.T., H.M., J.F. performed experiments. J.C., F.W., S.B., X.Z., P.P., D.C., V.J., and E.W. analysed data and interpreted results. R.H., C.C., P.K., and A.P. contributed specialist methods and resources. F.W., V.J., A.G., L.J., T.T., and E.W. designed experiments. E.W., J.C., V.J. and F.W. prepared the figures. E.W., J.C. and F.W. manuscript writing. R.H., P.K., C.C., L.J., T.T. and E.W. supervised.

## Declaration of Interests

The authors declare no competing interests.

## AI Disclosure Statement

Language editing, proof editing and coding were supported by AI tools. The authors take full responsibility for the contents of this manuscript.

## METHODS

All genomics analyses were performed using the mouse GRCm38 (mm10) reference genome assembly. Statistical tests were two-sided unless stated otherwise.

### Animals and exercise protocols

C57BL/6J mice were maintained in a certified specific-pathogen–free (SPF) facility under controlled temperature and a 12-h light–dark cycle with ad libitum access to standard chow and water. Following one week of acclimation, mice were randomly assigned to exercise-training or sedentary control groups. Three independent cohorts were used in this study. For the long-term cohort, used for snRNA-seq and snATAC-seq, mice were allocated at 5 months (young) or 13 months (aged) of age and underwent treadmill exercise for 6 months. Exercise-trained mice ran on a motorized treadmill at 28 m/min (young) or 25 m/min (aged) for 45 min per day, 5 days per week^39,40^. This moderate-intensity endurance protocol corresponds to approximately 70% of maximal aerobic capacity in this strain. Sedentary mice underwent sham treadmill exposure. At study completion, mice were 11 months (young) or 19 months (aged) old. Aging and exercise effects were estimated as separate pairwise contrasts against a common aged sedentary reference (ageing, ASed − YSed; exercise, AEx − ASed). Because the two share a reference group with opposite sign, statistics combining them are not independent, and are treated accordingly throughout.

We further examined independent cohorts spanning both age classes and exercise training duration, to assess whether the regulatory effects observed in the primary cohort generalized across age and training contexts. For the short-term cohorts, mice were studied at 5 months (young) or 22 months (aged). Aged exercise-trained mice underwent treadmill exercise at 18 m/min for 45 min per day, 5 days per week for 10–11 weeks. An independent long-term imaging cohort was initiated at 3 months of age and underwent 12 months of treadmill exercise using the same protocol. Mice were assigned to sedentary or exercise-trained groups only (*n* = 5 per group). All long-term experiments were performed at the Memorial Sloan Kettering Cancer Center, whereas short-term studies were performed at the Victor Chang Cardiac Research Institute.

### Cardiac tissue collection and processing

Mice were euthanized by CO₂ asphyxiation and perfused with ice-cold phosphate-buffered saline (PBS) to remove circulating blood prior to tissue collection. Hearts were rapidly excised, trimmed of surrounding connective tissue and major vessels, and rinsed in ice-cold PBS to remove residual blood. Cardiac tissues were snap-frozen in liquid nitrogen and stored at −80 °C until further processing.

### Nuclei isolation and single-nucleus sequencing

Frozen hearts were minced and homogenized in a hypotonic lysis buffer (91 mM NaCl, 5.3 mM KCl, 0.5 mM MgCl₂, 0.293 mM CaCl₂, 10 mM glucose, and 10 mM HEPES [pH 7.4]), supplemented with 0.05% Triton X-100, 0.5 mM spermidine, and 0.2 U/μl RNase inhibitor. Homogenates were centrifuged at 500 × g for 5 min at 4 °C. Pellets were washed in 1× DPBS containing 2% BSA and filtered through a 40 μm cell strainer and resuspended in 1× DPBS supplemented with 2% BSA and 0.2 U/μl RNase inhibitor. Following purification, nuclei quality and integrity were assessed by DAPI and Lamin A/C immunostaining, and DNA-content profiling confirmed a consistently >90% 2N fraction, with no difference in ploidy between conditions (**Figure S1A–S1C).**

For snRNA-seq, isolated nuclei were stained with Hoechst 33258 (2 µg/ml) and sorted by FACS to exclude debris and aggregates, using an initial scatter gate, four sequential singlet-discrimination gates, and a final DAPI area gate to retain nuclei within the expected ploidy range. Libraries were generated using the 10x Genomics Chromium platform^41^. For each library, nuclei from one male and one female mouse within the same experimental condition were combined at equal proportions and later demultiplexed based on sex-specific gene expression. For snATAC-seq, sorted nuclei were permeabilized with 0.05% digitonin for 1 min, and processed using the 10x Genomics Chromium Single Cell ATAC v1 platform^42^. snATAC-seq libraries were generated across eight Chromium runs using an experimental design matched to that of the snRNA-seq libraries. All snRNA-seq and snATAC-seq libraries were sequenced on an Illumina NovaSeq 6000 using paired-end sequencing, according to the manufacturer’s instructions.

### Processing and analysis of snRNA-seq and snATAC-seq data

snRNA-seq reads were aligned to the mouse mm10 reference transcriptome (refdata-gex-mm10-2020-A) with Cell Ranger (v7.0.0)^41,43^, with intronic reads included to account for pre-mRNA signal in nuclei. One low-yield library, containing two AEx samples (one male and one female), was excluded. Ambient RNA and empty droplets were removed with CellBender remove-background (v0.3.0; expected-cells = 20000, total-droplets-included = 30000, fpr = 0.01, epochs = 150, low-count-threshold = 150)^44^. Seurat (v4.4.0)^45,46^ in R (v4.2.1)^47^ was used for downstream analysis, and nuclei passing QC thresholds were retained (≤ 5% mitochondrial reads and 500-5,500 detected genes)^48,49^.

snATAC-seq reads were aligned to the mouse mm10 reference genome (refdata-cellranger-atac-mm10-1.2.0) using Cell Ranger ATAC count (v1.2.0)^50^, and peaks between 20 bp and 10 kb were retained. Downstream analysis was performed using Signac (v1.15.0)^51–53^. Nuclei passing QC thresholds (FRIP > 30%, peak-region-fragments > 5,000, blacklist ratio < 0.05, TSS enrichment > 2 and a nucleosome signal < 4) were doublet filtered using scDblFinder (v1.13.8, aggregateFeatures = TRUE)^54^ utilizing precomputed Seurat::FindClusters as input. After quality control and doublet removal, the final snATAC-seq dataset contained 65,701 nuclei.

Pooled male and female samples were demultiplexed using sex-based GLM classifiers trained on matched lung scRNA-seq data from the same animals (**Supplementary Methods**). After removal of two low-yield AEx samples and one low-assignment YSed female, the final snRNA-seq dataset comprised 13 animals: 3 YSed, 4 YEx, 4 ASed and 2 AEx. For snATAC-seq, peaks were recalled with MACS3 (v3.0.0b1)^55,56^ for each individual and merged using GenomicRanges::reduce (v1.50.2)^57^. Sequencing-run batch effects were assessed using Spearman correlations of normalized pseudobulk profiles (**Supplementary Results**). snRNA-seq cell types were annotated using canonical marker-gene expression (**Table S30**), and snATAC-seq cell identities were assigned by LSI-based label transfer from the snRNA-seq as reference (**Supplementary Methods**)^58–60^. Enhancer–promoter ATAC co-accessibility was inferred with Cicero (v1.3.9)^61^ using a maximum distance of 500 kb and a co-accessibility cutoff of 0.2.

### Differential gene expression and gene ontology analysis of snRNA-seq data

Cell-type-specific differential expression was performed to identify transcriptional responses to aging and exercise using MAST (v1.24.1)^62^ on log_2_(CPM + 1) values for genes detected in > 20% of nuclei. Models included fixed effects for sex and cellular detection rate (CDR)^63,64^. Differential-expression contrasts tested aging (ASed versus YSed) and exercise (AEx versus ASed); significance required FDR < 0.05 and |log_2_FC| > 0.5. These fixed-effects models served as the primary discovery analysis. However, these models do not explicitly account for animal-level replication, hence we also performed animal-aware analyses using mixed-effects models and per-animal pseudobulk which accounted for biological replicates (**Supplementary Results**). Age-by-exercise interaction terms were used to determine whether the transcriptional effect of exercise differed between young and aged animals.

GO enrichment was performed using clusterProfiler (v4.6.2)^65^, org.Mm.eg.db (v3.16.0)^66^ and gseGO on genes ranked by log_2_FC. Pathway-level attenuation scores were calculated from differential-expression effect sizes. KEGG pathway gene sets were retrieved using KEGGREST (v1.50)^67–69^ and pathway-level attenuation scores were calculated for each cell type (*c*) as the negative summed product of aging and exercise log_2_FC across expressed pathway genes (*g*), weighted by the proportion of pathway (*p*) genes expressed in that cell type (*PropExpressed_c,p_*).

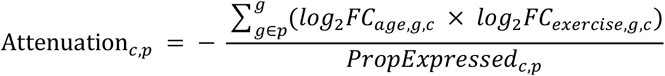

### Differential chromatin accessibility analysis of snATAC-seq data

Cell-type-resolved differential accessibility was performed using per-animal pseudobulk profiles, with the animal as the experimental unit. For each cell type, peaks were recalled independently per replicate using MACS3 (v3.0.0b1, effective genome size = 1.87 × 10⁹, extension size = 200 bp)^55,56^ and merged into cell-type-specific peak sets using GenomicRanges::reduce (v1.50.2, 1 bp overlap)^57^, excluding peaks overlapping the mm10 ENCODE blacklist^70^.

Pseudobulk counts were size-factor normalized and regions tested for differential accessibility using DESeq2 (v1.38.3)^71^, with sex as a covariate and experimental condition as the primary factor. Low-abundance regions were excluded by retaining peaks with ≥4 normalized counts in at least four replicates. Candidate differentially accessible regions (P < 0.1) were evaluated for gene set enrichment using rGREAT (v2.0.2, tss_source = “mm10”)^72,73^.

Within each cardiac cell type, peaks were ranked by GC content and divided into three terciles. Mean age-associated accessibility changes were calculated for each tercile across cell types. Peaks with nominal age-associated accessibility (P < 0.1) were classified as age-gained or age-lost and GC-content distributions were compared within each cell type using Wilcoxon rank-sum tests.

### Cell-cell communication analysis of snRNA-seq data

Cell–cell communication was inferred using CellChat (v1.6.1)^74^ after excluding low-abundance cell types. For each condition, overexpressed genes and interactions were determined using identifyOverExpressedGenes and identifyOverExpressedInteractions, and communication probabilities estimated with computeCommunProb (min.cells = 10). Differential communication strength was calculated for aging (ASed versus YSed) and exercise (AEx versus ASed). ECM signalling was assessed using the CellChat COLLAGEN pathway and inflammatory signalling assessed using a custom pathway containing genes in the ‘positive regulation of inflammatory response’ (GO:0050729) gene set that intersected with CellChat ligands, receptors or cofactors.

### Transcriptomic age prediction

Cardiac bulk RNA-seq data from *Tabula Muris Senis*^22^ (n = 54; **Table S31**) were used to construct a transcriptomic age predictor. We combined snRNA-seq pseudobulk profiles from this study and the *Tabula Muris Senis* dataset based on genes with > 10 counts across all samples. Counts were normalized by the trimmed mean of M values (TMM) method with edgeR (v3.40.2)^75^, log₂(CPM + 1)-transformed and batch-adjusted using limma::removeBatchEffect (v3.54.2)^76^, adjusting for sex and experimental effects. A glmnet LASSO model (v4.1.4; α = 1, standardize = TRUE)^77,78^ was trained on normalized *Tabula Muris Senis* profiles with a 70/30 training/test split. λ.min was determined by five-fold cross-validation and held-out performance was assessed by Pearson correlation and mean absolute error. The final age predictor comprised 33 genes and was applied to replicate-level pseudobulk snRNA-seq profiles to predict transcriptomic age.

### Echocardiography and speckle-tracking strain analysis

Echocardiography was performed using a Vevo 3100 high-resolution ultrasound imaging system. Mice were anesthetized with 1–1.5% isoflurane and maintained on a heating platform to preserve body temperature. Heart rate and respiratory rate were continuously monitored throughout imageing. Two-dimensional (B-mode) and M-mode images were acquired from parasternal long-and short-axis views. Left ventricular internal diameters at end-diastole (LVIDd) and end-systole (LVIDs), together with interventricular septum and posterior wall thickness, were measured from M-mode images at the level of the papillary muscles. Fractional shortening (FS) was calculated and averaged across three consecutive cardiac cycles. Myocardial and endocardial deformation was assessed by speckle-tracking–based strain analysis. Global longitudinal strain (GLS) was calculated using Vevo LAB (v5.11.0, FUJIFILM VisualSonics, Toronto, Canada), following manual delineation of endocardial borders with visual verification of tracking quality. Group differences were assessed using Type II two-way analysis of variance (ANOVA) with treatment group and sex as fixed factors. We obtained sex-adjusted values by regressing the measured values by sex and adding model residuals to the grand mean. Pairwise comparisons were performed using Tukey’s honestly significant difference (HSD) test with family-wise error rate correction (two-sided α = 0.05). Analyses were performed in Python 3.12 using SciPy (v1.14)^79^ and statsmodels (v0.14)^80^.

### Chromatin TF footprinting analyses

TF footprinting was performed using TOBIAS (v0.17.0)^81^ in cell types with > 5,000 accessible peaks to identify condition-associated changes in motif occupancy across cell types. For motif-site discovery, snATAC-seq fragments were pooled across biological replicates per cell type and condition, corrected for Tn5 insertion with ATACCorrect excluding mm10 blacklist regions, and footprint scores computed across peaks identified in ≥ three replicates per condition. Biological consistency was then evaluated from animal-level pairwise binding-score contrasts.

To test for differences in inferred TF occupancy, motif sites were identified by scanning accessible peaks with 790 mouse CIS-BP motifs (v2.0) using TOBIAS BINDetect^82^. For each cell type, BINDetect compared the mean log*2*FC in footprint scores for aging (ASed versus YSed) and exercise (AEx versus ASed) with a background distribution estimated from 100 permutations of randomly sampled positions using a one-sample t-test. A profile was classified as exercise-opposed when both contrasts had FDR < 0.05 and opposite effect directions. Biological consistency was then evaluated from animal-level pairwise binding-score contrasts. A motif was thus considered as exercise-opposed when its age and exercise contrasts were both significant (FDR < 0.05) and had opposite effect directions in a cell type. Age-response classes were defined using the aging contrast alone: age-up (ASed footprinted, YSed not), age-down (YSed footprinted, ASed not) or age-stable (no age-associated change). Exercise-response classes were assigned separately using AEx versus ASed as exercise-up, exercise-down or exercise-stable, without changing the locked age-response labels. AP-1 sites were linked to genes if located within 200 kb of a TSS. TRE motifs were assigned to FOS/JUN family members, whereas CRE motifs were assigned to CREB3, CREM and ATF1/2/4/5/6/7.

Using non-negative matrix factorization (NMF), we summarized snATAC-seq peaks using TF footprints into 10 modules per cell type. These were then sorted into GC-content terciles by mean peak GC content. Next, we calculated module-level aging and exercise binding shifts as the mean TOBIAS binding-score difference (*B*) across all sites (*Np*) linked to genes in each module:

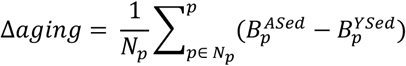

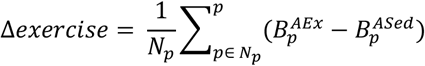

Differences between GC terciles were tested using two-sided Wilcoxon rank-sum tests. Opposition of ageing-associated occupancy by exercise was quantified via Spearman correlation between aging and exercise effects.

To test GC-dependent aging effects accounting for repeated measurements, standardized module scores from young sedentary and aged sedentary animals were analysed using a crossed linear mixed-effects model implemented in ‘lme4’ (v1.1-35.1): standardized module score ∼ age × GC-tercile + sex + (1 | animal) + (1 | module). Models were fitted by restricted maximum likelihood (anova() on ML fits), and the age-by-GC-tercile interaction was tested by likelihood-ratio test with a nested model lacking the interaction. Exercise effects in aged animals were evaluated using the analogous model: standardized module score ∼ exercise × GC-tercile + sex + (1 | animal) + (1 | module). Model-estimated exercise effects and 95% confidence intervals were calculated for each GC tercile. The exercise-by-GC-tercile interaction tested whether exercise differentially affected the GC gradient.

For an experimental-unit-level summary, a GC-redistribution index was calculated for each animal as the mean standardized activity of low-GC modules minus the mean standardized activity of high-GC modules. Aging was assessed by comparing this index between young sedentary and aged sedentary animals, and the exercise effect by comparing aged exercised with aged sedentary animals using two-sided Welch’s t-tests. Additionally, for each condition, the GC-binding contrast was calculated as mean inferred binding in low-GC modules minus that in high-GC modules. The fraction of the ageing-associated contrast opposed by exercise was calculated as −(mean AEx − mean ASed)/(mean ASed − mean YSed) × 100.

### CUT&Tag

CUT&Tag of the key AP-1 subunit c-FOS was performed on whole hearts from an independent short-term exercise cohort^83^ (**Supplementary Methods**; **Table S32**), with libraries sequenced on the Illumina NextSeq 2000 platform. Sequences were aligned to mm10 and peaks called with SEACR using nf-core/cutandrun (v3.2.2)^84,85^. *E. coli* spike-in reads were quantified by the pipeline and used to derive sample-specific normalization factors. CUT&Tag peaks were analysed with HOMER (v5.1, -size given)^86^. Consensus-peak counts were retained per animal, filtered with filterByExpr and analysed with edgeR (v3.40.2)^75^ using spike-in-derived normalization factors and quasi-likelihood F-tests for aging (ASed versus YSed) and exercise (AEx versus ASed), with animals as the experimental unit. Age-up c-FOS peaks were defined solely from the aging contrast (ASed versus YSed; log_2_FC > 0, P < 0.05). This set was then locked before the exercise contrast was examined; age-up peaks with reduced c-FOS occupancy in AEx relative to ASed mice (log2FC < 0, P < 0.05) were annotated downstream as age-up, exercise-opposed. c-FOS CUT&Tag peaks were assigned to TRE+CRE, TRE-only, CRE-only, and motif-negative categories based on FIMO (v5.5.5)^87^ motif detection. Enrichment of motif classes among peaks exhibiting ageing-associated gain and exercise-associated attenuation was assessed using competitive gene-set testing with the CAMERA test (‘limma’ v3.58.1).

### AP-1 regulatory element analysis

To determine whether AP-1 occupancy corresponded to measurable snATAC-seq accessibility differences, TOBIAS-defined AP-1 peak accessibility was averaged across nuclei by condition and cell type. Accessibility at age-up versus stable AP-1 peaks was compared by two-sided Wilcoxon rank-sum tests. Cell-type specificity of AP-1 binding was quantified by overlapping AP-1 peaks (≥1-bp overlap). To test whether dynamic AP-1 elements were depleted from high-contact cardiomyocyte anchors, published mouse cardiomyocyte Hi-C loops (PMID: 28802249)^88^ identified using Juicer HiCCUPS (v2.0, KR normalization, 10 kb resolution, observed/expected enrichment > 1)^89^ were overlapped with c-FOS TOBIAS peaks. Co-binding motif analyses used sites within 50 bp of bound AP-1 motifs with motifs detected < 10 times excluded to reduce unstable estimates.

For each cell type, XGBoost models (v2.1.4; 350 estimators, maximum depth 5, learning rate 0.05)^90^ were trained to distinguish dynamic from stable c-FOS sites based on TF-footprint density in the 50 bp window surrounding c-FOS motifs. HSPCs and lymphatic ECs were excluded because of low peak counts. Performance was assessed by five-fold stratified cross-validation and summarized by mean ROC AUC after interpolation to 300 false-positive-rate points.

### Tissue processing

Hearts were fixed in 4% paraformaldehyde overnight, cryoprotected in 30% sucrose, embedded in optimal cutting temperature (OCT) compound, and cryosectioned. All samples used for comparative analyses were processed in parallel under identical conditions.

### Histology and image analysis

Picrosirius red staining was performed on fixed-frozen heart sections from the short-term cohort using a commercial kit (Picrosirius Red, ab245887, Abcam) for 1.5 h following the manufacturer’s protocol, and sections were mounted using DPX. Brightfield images of whole heart sections were acquired using a slide scanner (VS200, Olympus; SLIDEVIEW) equipped with a 40× objective, under identical acquisition settings. Collagen and myocardial regions were segmented using stain vector–based pixel classification in QuPath (v0.7.0)^91^ across manually annotated anatomical regions of interest (LV, RV, LA, RA, LV perivascular, LV interstitial, RV perivascular, RV interstitial), and collagen-to-myocyte area ratios were calculated for each region. Analysed regions of interest included ventricular and atrial myocardium: the epicardium, the valves and the atrial-ventricular regions were excluded from the analysis.

### Immunofluorescence staining

Sections were blocked and permeabilized, then incubated with primary antibodies overnight at 4 °C, followed by Alexa Fluor–conjugated secondary antibodies and DAPI nuclear counterstaining (Merck Life Science, cat. no. D9542). CD163 (BioLegend, cat. no. 155307), CD34 (Abcam, cat. no. 198395), CD68 (Bio-Rad, cat. no. MCA1957), and Lumican (Abcam, cat. no. 168348) staining were performed using identical conditions across experiments. Images were acquired using a Zeiss LSM900 confocal microscope or an Olympus VS-series slide-scanning system, with acquisition settings kept constant within each experiment.

CD163/DAPI quantification images were acquired across distinct anatomical regions. CD163 immunofluorescence was quantified in ImageJ/Fiji (v1.54s) from 16-bit images using background subtraction and fixed-threshold segmentation, followed by particle-based quantification of CD163⁺ cells. For each region, the number of CD163⁺ cells was expressed as a proportion of total DAPI⁺ cell count, and per-animal mean CD163⁺/DAPI⁺ ratios were used for statistical comparison.

### CD34/CD68 and Lumican/CD68 co-localization

Double-positive cells were identified by high-resolution z-stack confocal microscopy with orthogonal sectioning to confirm co-localization; these data were not quantified. Z-stack acquisition of all stained sections was performed using an LSM900 inverted confocal laser scanning microscope; confocal scanner detection and laser excitation settings were kept identical between samples wherever comparisons were made. Z-stacks were processed using Fiji (v1.54s), and distinct LUTs were applied to each channel to improve visibility.

### Phospho-c-JUN quantification

The subcellular distribution and expression pattern of p-c-JUN observed by immunofluorescence were in accordance with those previously published^92^. Nuclear p-c-JUN signal was quantified in CellProfiler (v4.2.8) by segmenting DAPI-positive nuclei and measuring mean fluorescence intensity in the Alexa Fluor 647 channel. Image-level measurements were generated by averaging all nuclei within each image, and per-animal means were used for statistical comparison.

### Macfib analysis by flow cytometry

Pdgfra-MerCreMer mice (MGI Cat# 5475226) and R26-tdTomato reporter mice (MGI Cat# 3809523) were crossed to generate Pdgfra-reporter (PdgfraMCM/+; R26Tom/+) mice. Healthy adult Pdgfra-reporter mice were administered tamoxifen (Sigma-Aldrich, Cat# T5648) once daily (100 mg kg⁻¹ body weight) for three consecutive days to induce Cre-mediated recombination and permanent labelling of Pdgfra-expressing cells with tdTomato. Following tamoxifen administration, mice were allowed a 2-week washout period before tissue collection. Cell isolation, staining and flow cytometry procedures were performed as previously described^93^. Briefly, single-cell suspensions were stained with fluorophore-conjugated anti-CD45 antibody (BD Biosciences, Cat# 557659) and analysed by flow cytometry using a BD FACSFortessa analyser (BD Biosciences). CD45⁺ tdTomato⁺ cells within the Pdgfra lineage-traced population were identified as macfibs. Flow cytometry data were acquired using FACSDiva software and analysed using FlowJo software (version 10.8.2; BD Biosciences). Macfib frequency was quantified from three biological replicates.

### Macrophage annotation and analyses

snRNA-seq macrophages (n = 11,143) were re-embedded using the second to ninth principal components (PCs) and hierarchically clustered with too-many-cells (v3, min_size = 100 and smart_cutoff = 0.5)^94^ to reduce resolution-dependent border assignment. States were annotated as MHC-II^hi^, Cd163^hi^ and macfib populations based on *H2-Ab1*, *Cd163* and *Col3a1* expression, respectively. Additional analyses were performed to exclude the possibility of doublets in the macfib population (**Supplementary Results**).

Mouse macrophage states were compared with human cardiac scRNA datasets from Shu et al. 2022 (GSE213128, PMID: 36200446)^95^ and Amrute et al. 2023 (GSE226314, PMID: 37583573)^96^. Heart-valve-derived cells from Shu et al. (2022) were QC filtered (200–4,000 detected genes, < 10% mitochondrial reads, zero *HBB* counts) and myeloid cells reclustered with PCs 1–10 at a resolution of 0.5. Human genes were mapped to mouse orthologs using seuratTools::convert_human_seu_to_mouse (v0.4.12); one ortholog was selected at random when the mapping was ambiguous. Integration with mouse macrophages used Seurat::FindIntegrationAnchors and Seurat::IntegrateData (dims = 1–30)^45,46^. Macfib clusters were identified by overlap with annotated mouse macrophage states. Discriminating markers for the human macfib population were identified using Seurat::FindAllMarkers (Wilcoxon test, P < 0.001 and log_2_FC > 0.5). Macrophage-state signatures comprised the top 50 significant positive markers identified by Seurat::FindAllMarkers (Wilcoxon test) and ranked by log_2_FC. Per-patient pseudobulk counts were variance-stabilized using DESeq2 (v1.38.3)^71^, converted to gene-wise z-scores, averaged across signature genes, and correlated with ejection fraction by Pearson correlation.

### Lenti-STARR-seq reporter assay

Primary adult cardiac fibroblasts were isolated from 5-month-old female C57BL/6J mice (Jackson Laboratory) using collagenase digestion followed by 3 h preplating on gelatin-coated dishes (**Supplementary Methods**)^97^. Enhancer activity was assessed in cultured primary cardiac fibroblasts using lenti-STARR-seq MPRA^98^. The oligonucleotide library comprised 1,861 endogenous 170-bp sequences centered on c-FOS/AP-1 motif sites selected from the complete fibroblast peak set, including all dynamic sites and 452 stable control sites, together with matched full-sequence and motif-specific scrambled controls. The final library contained 5,583 oligonucleotides synthesized by Twist Bioscience. Vector-compatible overhangs were added by PCR, and the library was Gibson-cloned into EcoRI-digested pLS-mP (Addgene #81225) downstream of EGFP (**Table S33**). The lentiviral reporter library was packaged in HEK293T cells and concentrated as described in the **Supplementary Methods**. Primary cardiac fibroblasts were transduced with the concentrated lentiviral STARR-seq library and paired RNA and genomic DNA were collected for reporter quantification. Reporter-derived cDNA and matched genomic DNA input libraries were amplified, prepared for sequencing using NEBNext Ultra II DNA Library Prep Kit and sequenced on an Illumina NextSeq 2000 platform.

MPRA libraries were assessed using FASTQC (v0.12.1)^99^ and trimmed using fastp (v0.20.0)^100^. Reads were aligned to the MPRA reference with BWA (v0.7.17, MAPQ > 10)^101^. Element-level counts were generated using samtools idxstats (v1.22)^102^ and elements with < 10 reads in at least 3 replicates were removed, leaving 1,653 elements. Per-element transcription rates (α) and differential activity relative to scrambled controls were estimated in R with MPRAnalyze (v1.28.0)^103^. Differential activity between AP-1 binding classes was tested via likelihood-ratio tests comparing a full model containing sequence class and batch with a reduced model containing batch only. Additionally, to compare dynamic and stable enhancer classes while controlling for sequence and genomic-context differences, dynamic elements were matched to stable elements based on GC content and distance to the nearest TSS using nearest-neighbor propensity-score matching implemented in MatchIt (version 4.7.2, caliper = 0.2, ratio = 1)^104^.

### Sequence modeling and in silico mutagenesis

We adapted AlphaGenome^26^ for lenti-STARR-seq activity prediction by training a linear probe on frozen convolutional sequence representations. The resulting AlphaGenome-MPRA model^27^ predicts reporter activity from 200-bp constructs containing the assayed 170-bp genomic sequence. Models were trained on 1,653 endogenous AP-1 elements using cross-validation, sequence augmentation and fixed hyperparameters selected by grid search. Final predictions were generated using a nine-model ensemble and evaluated on held-out sequences. Model performance and robustness were assessed across multiple independent data partitions, including chromosome-partitioned splits (**Supplementary Methods**).

*In silico* saturation mutagenesis was performed for all 1,653 enhancer sequences. At each nucleotide, the three alternative bases were introduced independently, and positional importance was defined by the largest predicted reduction in reporter activity across the three substitutions. Further details on seqlet and perturbation analyses are in the **Supplementary Methods**.

We assessed different sequence models in their ability to predict sequence classes. We used four distinct classes of input data: 1) MPRA activity, 2) sequence composition and TSS distance, 3) motif augmented sequence composition features and 4) AlphaGenome sequence embeddings. Sequence-composition features comprised GC content, dinucleotide frequencies, homopolymer counts, trinucleotide entropy, AP-1 count and distance to the nearest TSS. Motif-augmented features additionally included PWM scores, motif counts, motif-family counts, and AP-1 spacing and orientation features. Performance was evaluated from held-out predictions and summarized by the area under the receiver-operating-characteristic curve (AUROC).

### In vitro IL-1β stimulation and AP-1 inhibition in cardiac fibroblasts

Primary cardiac fibroblasts were used in two independent perturbation experiments. For the AP-1 inhibition experiment, cells were treated with T-5224 (80 µM, MedChemExpress, cat. no. HY-12270) or vehicle for 6 h (three libraries per group). In a separate experiment, serum-starved cells were stimulated with IL-1β (Thermo Fisher Scientific; cat. no. 200-01B-10UG) for 3 or 6 h (**Supplementary Methods**). RNA was extracted, and libraries were prepared and sequenced as described below (‘Bulk RNA-seq’). Reads were processed using the nf-core/rnaseq pipeline (v3.14.0, STAR–Salmon, mm10 reference)^84^, and differential expression was performed using edgeR^75^ with TMM normalization and quasi-likelihood negative binomial models.

Cardiac fibroblasts were stimulated with IL-1β (10 ng/mL) for 15, 30 or 60 min, and CUT&Tag for c-FOS and c-JUN was performed for each condition and time point and processed as above. Due to differences in systematic FC bias, c-FOS CUT&Tag was normalized using *E. coli*-derived carry-over reads originating from the pAG-Tn5 preparation, rather than from deliberately added spike-in DNA, whereas c-JUN CUT&Tag was normalized using TMM normalization.

To compare IL-1β-induced AP-1 occupancy with ageing-associated AP-1 chromatin remodelling in vivo, IL-1β CUT&Tag peaks were intersected with fibroblast AP-1 footprint classes derived from TOBIAS analysis. AP-1 family motifs were pooled across FOS, FOSB, FOSL1, FOSL2, JUN, JUNB and JUND. Pathway enrichment analysis of AP-1-inhibited differentially expressed genes was performed using clusterProfiler (v4.6.2)^65^ against GO BP, KEGG, MSigDB Hallmark and Reactome gene sets^105^, using expressed genes as the background.

### Bulk RNA-seq

Total RNA was isolated from cells or nuclei using Direct-zol RNA Microprep Kits (Zymo Research, cat. no. R2060). RNA integrity was assessed using High Sensitivity RNA ScreenTape assay (Agilent, cat. no. 5067-5576), and total RNA was quantified using Qubit RNA HS assay kit (Invitrogen, cat. no. Q32852). ∼120 ng total RNA was used to prepare each cDNA library using Illumina Stranded Total RNA Prep with Ribo-Zero Plus kit (Illumina, cat. no. 20040529) following the manufacturer’s protocol. cDNA library concentration was calculated using Qubit dsDNA HS assay kit (Invitrogen, cat. no. Q32851). Libraries were pooled and sequenced on an Illumina NovaSeq X plus platform with paired-end 150 bp reads, targeting at least 20 million reads per sample.

### In vivo AP-1 inhibition

An AAV9 transfer plasmid expressing dominant-negative AP-1 (dnAP-1) under the control of a cardiomyocyte-specific cTnT promoter was generated by Gibson assembly. The construct contained dnAP-1-T2A-tdTomato and was assembled from CMV500-A-FOS, pHDR_DDX4-T2A-tdTomato-puroTK and pENN-AAV-cTNT-PI-eGFP-WPRE-rBG backbone vectors (**Supplementary Methods**). Plasmids were sequence verified, purified using a low-endotoxin plasmid preparation protocol and packaged as AAV9 by the CMRI Vectorology Facility (Sydney). Mice aged 22–25 months received a single retro-orbital injection of AAV9-dnAP-1 or AAV9-GFP control (1 × 10¹¹ viral genomes in 100 μl) and were harvested five weeks after administration.

CUT&Tag was performed on cardiac tissue using antibodies against c-FOS, total c-JUN and p-c-JUN. Libraries were sequenced on the Illumina NovaSeq X Plus platform and processed using the whole-heart c-FOS CUT&Tag workflow implemented in nf-core/cutandrun, including alignment to mm10, SEACR peak calling, consensus peak construction, motif annotation, read counting, normalization and differential occupancy testing. Peaks overlapping TOBIAS-defined bound AP-1 motifs in cardiomyocytes were classified as cardiomyocyte AP-1-associated sites and summarized by their mean TOBIAS AP-1 differential binding score. For each target, a relative peak-level log_2_FC was calculated by subtracting the genome-wide mean dnAP-1 log_2_FC from each peak’s log_2_FC, centering every peak on the global dnAP-1 effect.

snRNA-seq libraries from AAV9-treated hearts were sequenced on an Illumina NovaSeq 6000 using on-chip multiplexing (OCM) and processed as described above for the ageing–exercise cohort, with the following modifications. Reads were aligned with 10x Genomics Cell Ranger multi (v10.0.0)^41^. Ambient RNA was removed using CellBender^44^ remove-background (expected-cells = 8,000, total-droplets-included = 25,000, fpr = 0.01, epochs = 150, low-count-threshold = 100). Corrected matrices were imported into Seurat^45,46^ and filtered to retain nuclei with 500–5,000 detected genes and ≤5% mitochondrial reads. Putative doublets were removed using Scrublet^106^ with an expected doublet rate of 0.08. Cell types were assigned using canonical marker genes^58–60^. Differential expression between AAV9-dnAP-1 and AAV9-GFP-treated hearts was assessed using MAST^62^ for genes detected in >20% of nuclei per cell type, with treatment, cellular detection rate and OCM pool as fixed effects. Differentially expressed genes were defined by FDR <0.05 and |log_2_FC| ≥0.585 (a 1.5-fold change).

### In vivo IL-1β inhibition transcriptional and chromatin analyses

Female C57BL/6J mice aged 18.5 months received intraperitoneal injections of either a polyclonal Armenian hamster IgG isotype control antibody (InVivoMab, BE0091, lot no. 765821F1) or an anti-IL-1β neutralizing antibody (InVivoMab, BE0246, lot no. 745620M2), at a dose of 500 µg per injection every 3 days for a total of 10 injections. Three days after the final injection, cardiac function was assessed by transthoracic echocardiography over a 2-day period. Mice were euthanized and heart tissue was harvested the day after echocardiographic evaluation for subsequent analyses. Bulk RNA-seq and H3K27ac CUT&Tag were performed as described above. Sequencing-library alignment, peak calling, and differential occupancy analysis were performed using the nf-core/cutandrun^84^ and edgeR^75^ workflow described above. Two anti-IL-1β libraries with < 10,000 spike-in reads were excluded from further analyses. Consensus peaks were constructed from peaks identified in at least 2 replicates in a single condition.

Mouse Hallmark gene sets were obtained from MSigDB. Preranked gene-set enrichment analysis was performed using clusterProfiler::GSEA. For pseudobulk analysis, raw single-nuclei RNA-seq counts were summed by mouse. Lowly expressed genes were removed using edgeR::filterByExpr, libraries were TMM-normalized and differential expression was tested using edgeR quasi-likelihood negative-binomial models.

Directional age- and exercise-associated gene sets were derived from the pseudobulk differential-expression results and tested for enrichment in the anti–IL-1β RNA-seq data using CAMERA^107^. Cell-type-resolved TOBIAS AP-1 sites were assigned to the nearest expressed gene within 100 kb of a TSS. Genes were deduplicated before gene-level statistical testing. Associations between AP-1 linkage and anti–IL-1β transcriptional responses were tested by linear regression, adjusting for baseline expression.

H3K27ac peaks were classified according to overlap with cell-type-resolved AP-1 sites. Associations between AP-1 overlap and anti–IL-1β H3K27ac responses were tested by linear regression, adjusting for baseline signal and peak width. Responses of dynamic and stable AP-1-associated peaks were compared using Wilcoxon rank-sum tests.

Gene-level promoter H3K27ac responses were calculated by averaging responses across peaks overlapping −2 kb to +0.5 kb from the transcription start site and compared with the corresponding transcriptional responses.

To test whether IL-1β-responsive AP-1 sites were preferentially associated with in vivo H3K27ac, we focused on TOBIAS-defined AP-1 footprint centres within age-up regions and overlapped them with 60-min IL-1β-stimulated c-FOS CUT&Tag peaks from primary cardiac fibroblasts. For footprints overlapping multiple c-FOS peaks, the peak with the highest logCPM was retained. H3K27ac peaks were included if they overlapped at least one mapped AP-1 footprint; when multiple eligible AP-1 sites mapped to the same H3K27ac peak, their in vitro c-FOS log_2_FC were averaged. Within each AP-1 site set, in vitro c-FOS log_2_FC were z-scored across peaks. H3K27ac logCPM values were centred across animals, and an across-peak slope against the standardized in vitro response was calculated for each animal. The treatment effect was defined as mean anti-IL-1β minus mean IgG and tested by exact one-sided label permutation across all 126 assignments of four treated labels among nine animals, with lower slope after IL-1β neutralization prespecified.

### GWAS gene-to-AP-1 association

Human disease enrichment was assessed for genes linked to age-up, exercise-opposed AP-1 regulatory elements. Age-up, exercise-opposed sites identified by c-FOS CUT&Tag were retained only when supported in cardiomyocytes or fibroblasts by concordant, exercise-opposed changes in chromatin accessibility or TOBIAS-derived AP-1 footprinting. AP-1 sites were assigned to genes whose gene bodies lay within 20 kb. Mouse genes were mapped to human orthologs using babelgene (v22.9)^108^. Where multiple mouse orthologs mapped to the same human gene, the maximum site count across orthologs was retained. Human gene-level association statistics were calculated using MAGMA (v1.10)^109^ from GWAS summary statistics for coronary artery disease^110^, atrial fibrillation^111^ and QT interval^112^, hypertension^113^, C-reactive protein^114^, triglyceride/HDL ratio^115^, and rheumatoid arthritis^116^. SNPs were assigned to genes using the hg19 reference genome and a window extending 1 kb upstream of each transcription start site. Analyses were restricted to human orthologs of protein-coding genes expressed in the mouse heart, yielding approximately 15,000 genes per trait. MAGMA Z scores were interpreted as measures of gene-level association strength rather than direction of effect. Mean gene-level MAGMA Z scores were calculated for each group and trait. Enrichment was tested using MAGMA competitive gene-set analysis, conditioning on gene size, gene density and minor allele count.

### Ethics Statement

All animal experiments were approved by the Memorial Sloan Kettering Cancer Center Institutional Animal Care and Use Committee (IACUC; protocol 17-11-008) and the Garvan Institute/St. Vincent’s Hospital Animal Experimentation Ethics Committee (protocols 22_15 and 25_08). All procedures were performed in accordance with institutional guidelines and the National Health and Medical Research Council (NHMRC) of Australia guidelines for the care and use of laboratory animals. All efforts were made to minimize animal suffering.

## Supplementary Figures

**Supplementary Figure 1.**
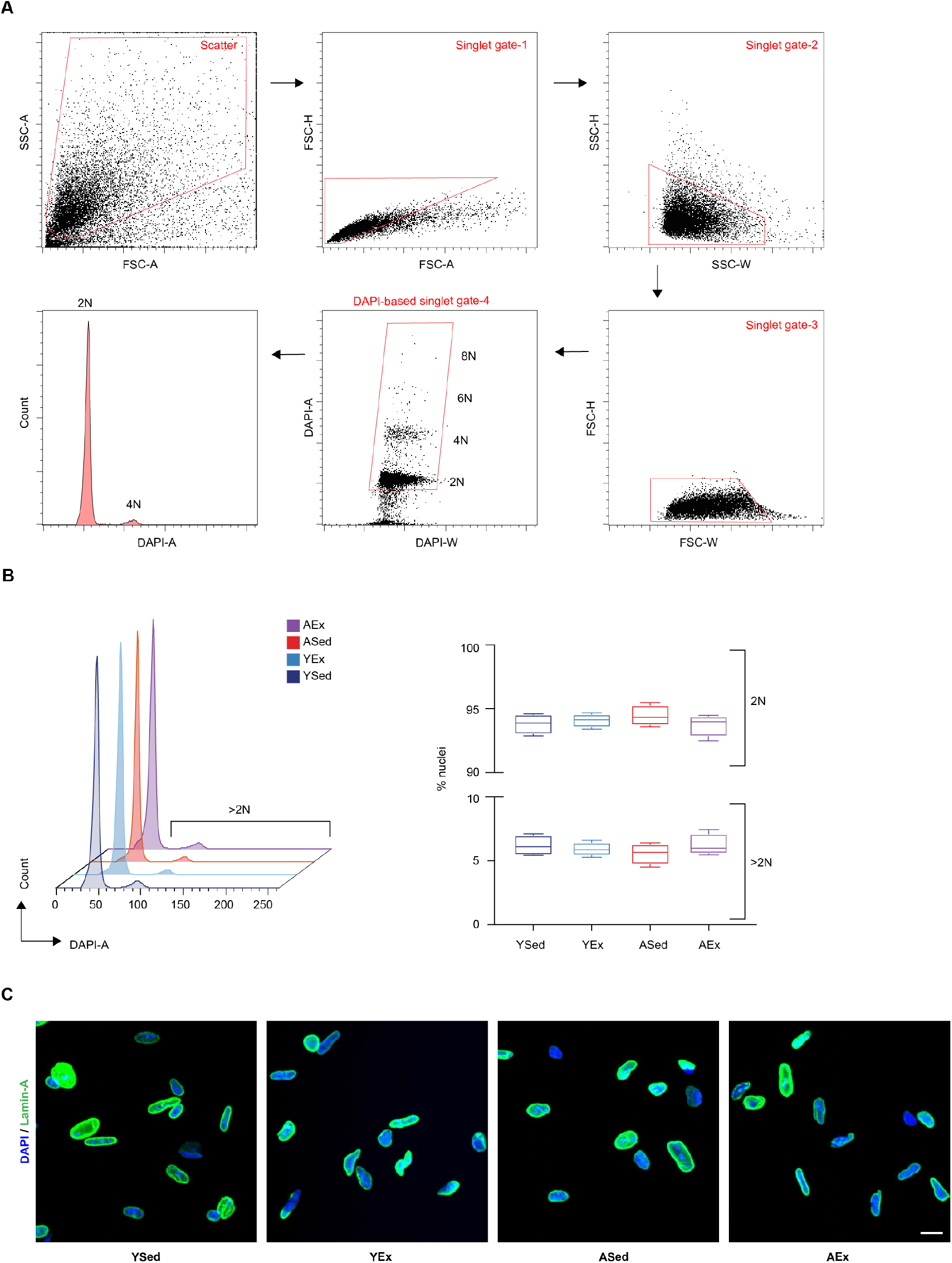
Nuclear sorting and ploidy quality control. **(A)** Sequential fluorescence-activated cell sorting (FACS) gating strategy used for single-nucleus isolation. Gates are shown in the order indicated by the arrows: an initial scatter gate to enrich for intact nuclei; three successive singlet discrimination gates using FSC and SSC area, height and width parameters; a DAPI-based singlet gate used for final sort selection; and a representative DAPI-area histogram showing the discrete 2N and 4N ploidy peaks used for final sort selection. **(B)** Left, overlaid DAPI-area histograms of nuclear DNA content across the four experimental groups (YSed, YEx, ASed and AEx), illustrating the predominant 2N peak with a minor >2N fraction. Right, quantification of 2N and >2N nuclear proportions across experimental groups, shown as boxplots displaying the median and interquartile range (IQR) (n = 4 animals per condition). **(C)** Representative immunofluorescence images of FACS-sorted nuclei stained with DAPI (blue) and anti-Lamin-A (green) across the four experimental groups (YSed, YEx, ASed and AEx), confirming preservation of nuclear integrity after sorting. Scale bar, 10 µm.

**Supplementary Figure 2.**
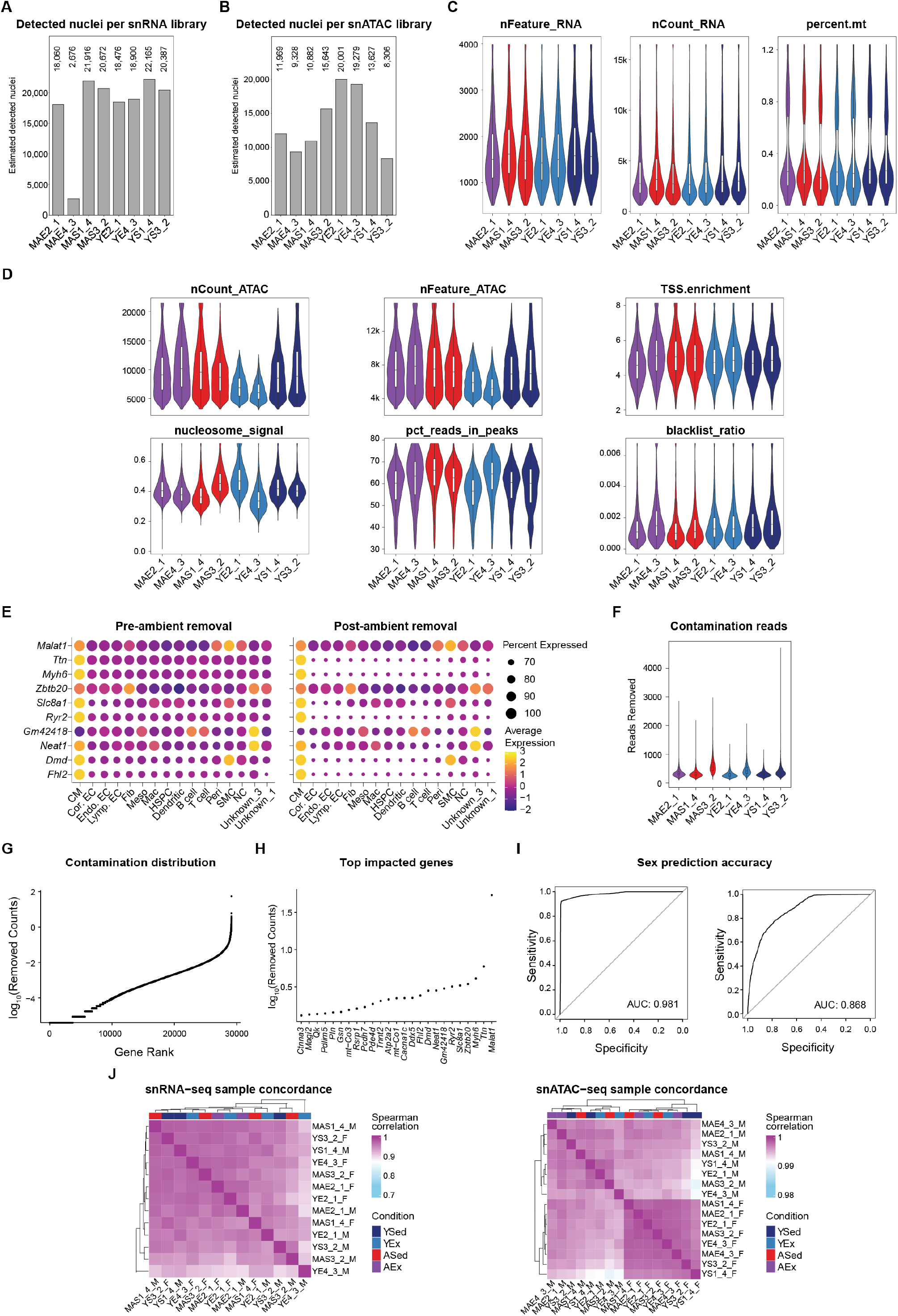
Single-nucleus sequencing quality control. **(A)** Estimated number of detected nuclei per snRNA Cell Ranger run. **(B)** Estimated number of detected nuclei per snATAC Cell Ranger run. **(C)** snRNA-seq quality-control metrics across sequencing runs, including the number of detected genes, total RNA counts and mitochondrial read percentages. **(D)** snATAC-seq quality-control distributions across sequencing runs, including total ATAC counts, detected ATAC features, TSS enrichment, nucleosome signal, percentage of reads in peaks and blacklist ratio. **(E)** Dot plots showing expression of the top ambient RNA-associated features before and after CellBender ambient RNA removal across annotated cell types. Point size indicates the percentage of nuclei expressing each feature, and color indicates average expression. **(F)** Number of reads removed per nucleus by CellBender for each snRNA-seq library. **(G)** Genes ranked by total removed counts after ambient RNA correction. **(H)** Top genes with the highest removed-count burden following ambient RNA correction. **(I)** Receiver-operating characteristic (ROC) curves for sex-assignment generalized linear models used during sample demultiplexing, shown for the RNA-based classifier (left) and the ATAC-based classifier (right). Area under the curve (AUC) values are shown for each plot. **(J)** Spearman correlation heatmaps showing pairwise sample similarity for snRNA-seq and snATAC-seq pseudobulk profiles, annotated by experimental condition.

**Supplementary Figure 3.**
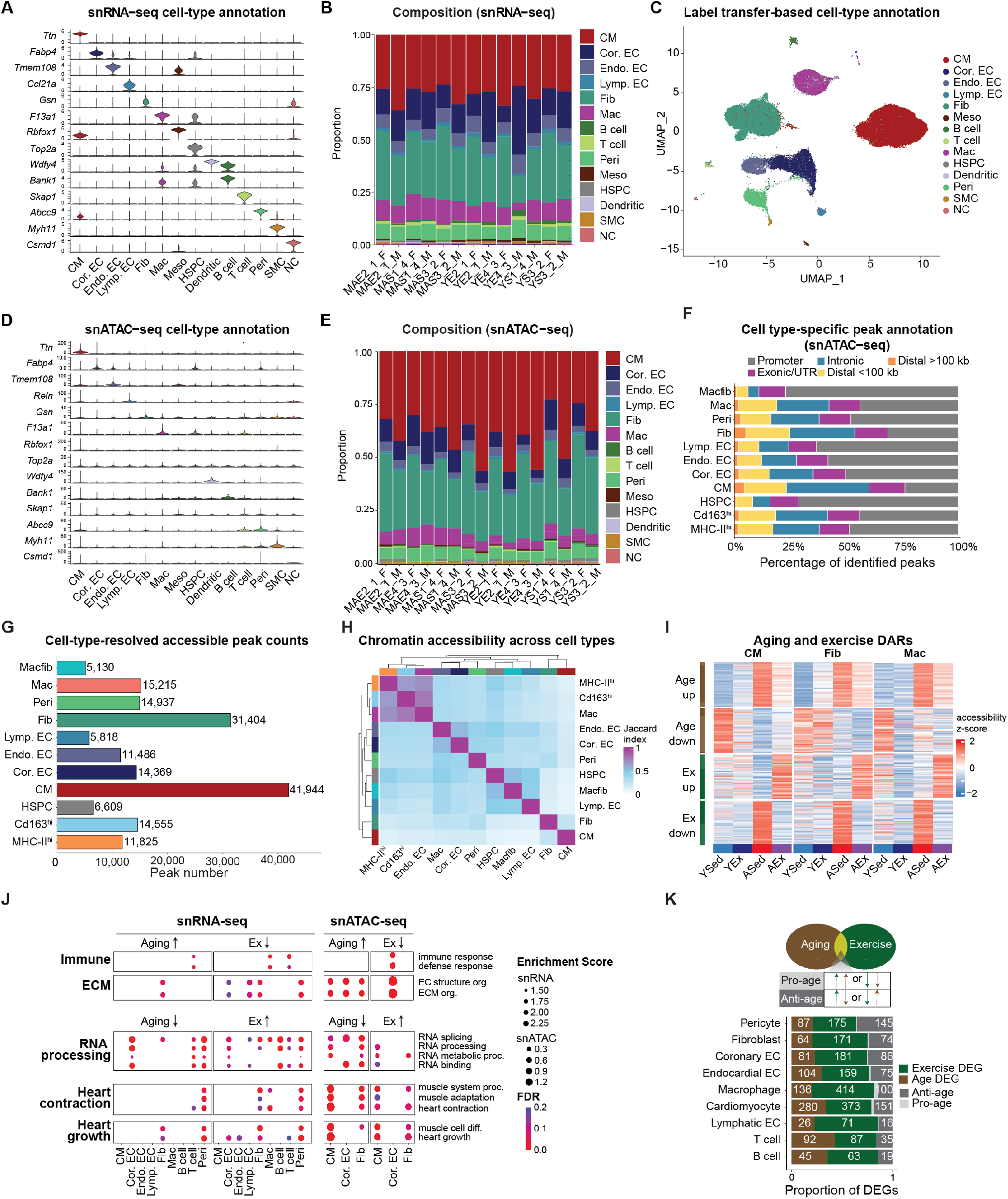
Cell-type annotation, peak summary and GO term enrichment. **(A)** Violin plots showing expression of canonical marker genes across snRNA-seq cell-type annotations. **(B)** Proportion of snRNA-seq nuclei assigned to each cell type across individual animals. **(C)** UMAP embedding of the snATAC-seq dataset colored by label-transfer-predicted cell type, using the annotated snRNA-seq dataset as the reference. **(D)** Violin plots showing accessibility of canonical marker loci across snATAC-seq cell types. **(E)** Proportion of snATAC-seq nuclei assigned to each cell type across sequencing libraries. **(F)** Genomic annotation of tested snATAC-seq peaks by cell type, shown as the proportion of peaks assigned to promoter, exonic/UTR, intronic, distal <100 kb and distal >100 kb categories. **(G)** Number of unique tested peaks per cell type after filtering. Bar labels indicate peak counts. **(H)** Pairwise Jaccard similarity of tested peak sets across cell types. **(I)** Normalized accessibility of the top 100 differentially accessible peaks, ranked by log₂FC and grouped by direction of change, for selected cell types and contrasts. Values are row-scaled z-scores. **(J)** Curated significant GO terms (FDR < 0.2) showing opposed enrichment with aging (ASed – YSed) and exercise (AEx – ASed) across cell types. RNA-based results used GSEA ranked differential-expression log₂FC values; ATAC-based enrichments were determined using GREAT based on differentially accessible peaks. **(K)** Classification of differentially expressed genes as ageing-only, exercise-only, exercise-opposing or exercise-reinforcing across cell types. Significant DEGs were defined as FDR < 0.05 and |log₂FC| > 0.5.

**Supplementary Figure 4.**
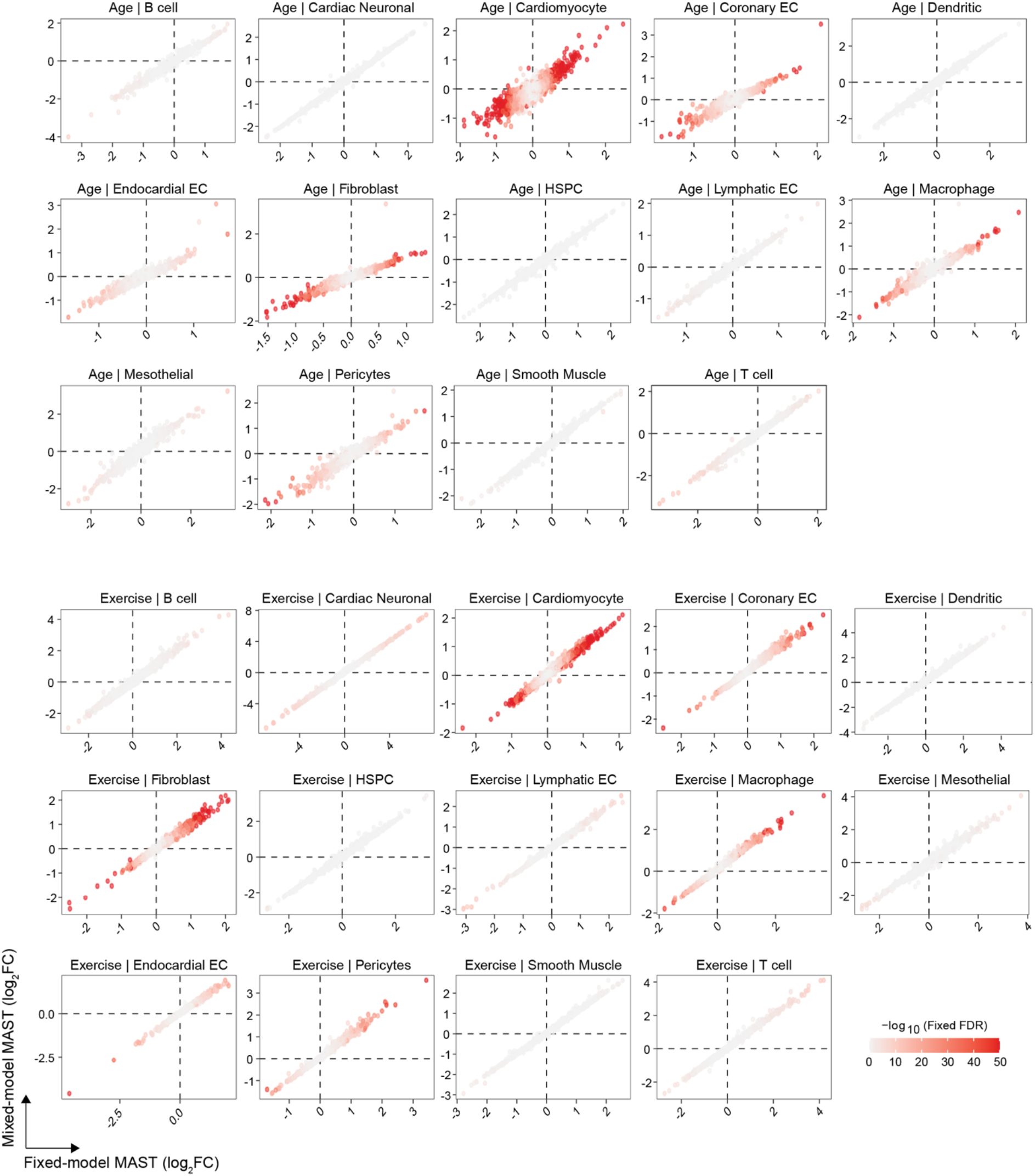
Concordance between fixed-effects and mixed-effects MAST differential-expression models. Scatter plots comparing gene-wise log_2_FC estimates derived from fixed-effects MAST models and mixed-effects MAST models including animal-level random effects for aging and exercise contrasts across cell types. Points represent tested genes and are colored by −log_10_(FDR) from the fixed-effects model. Dashed lines indicate zero effect in each model.

**Supplementary Figure 5.**
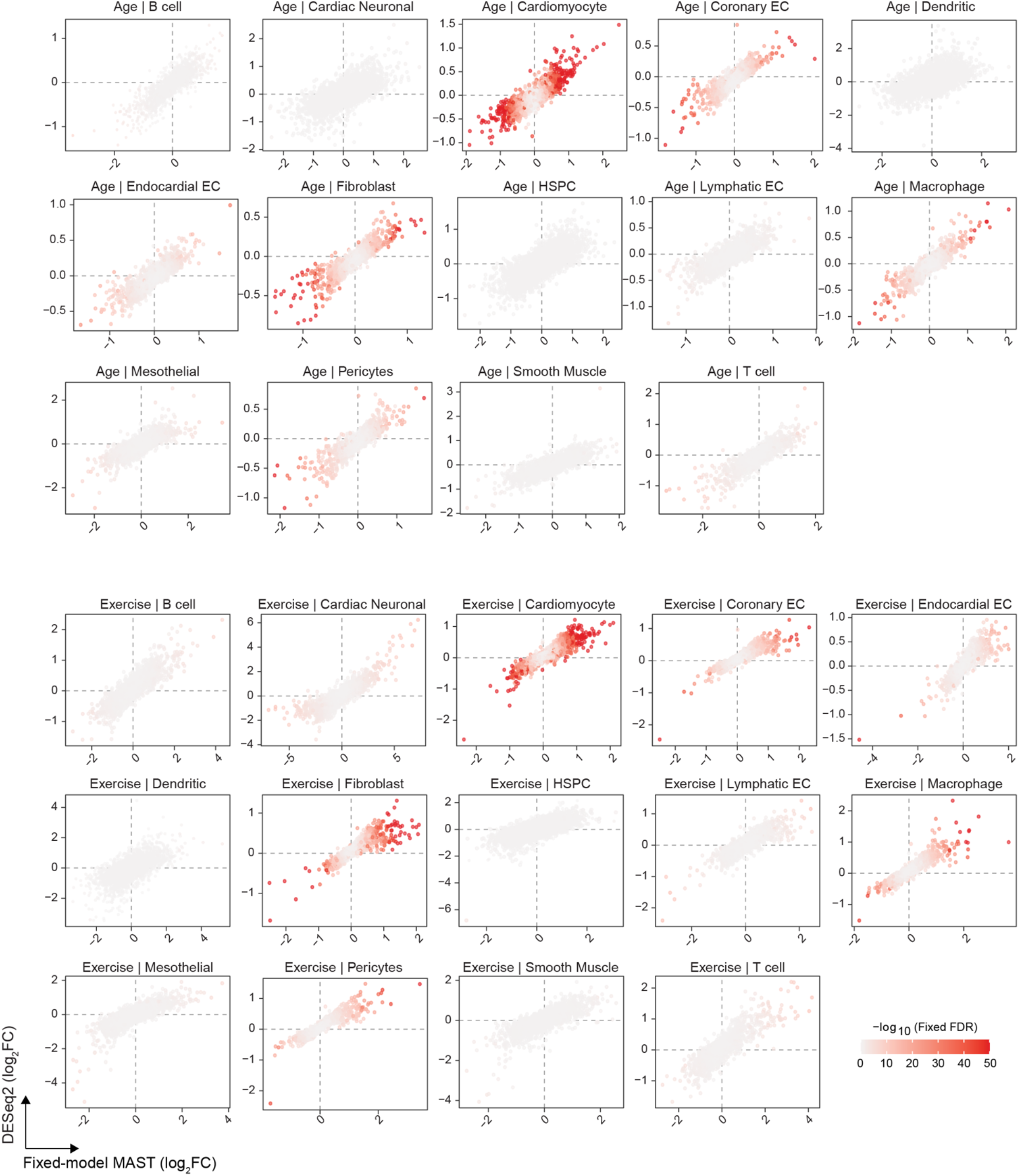
Concordance between fixed-effects MAST and pseudobulk DESeq2 differential-expression models. Scatter plots comparing gene-wise log_2_FC estimates from the fixed-effects MAST models and pseudobulk DESeq2 models for aging and exercise contrasts across cell types. Points represent tested genes and are colored by −log_10_(FDR) from the fixed-effects model. Dashed lines indicate zero effect in each model.

**Supplementary Figure 6.**
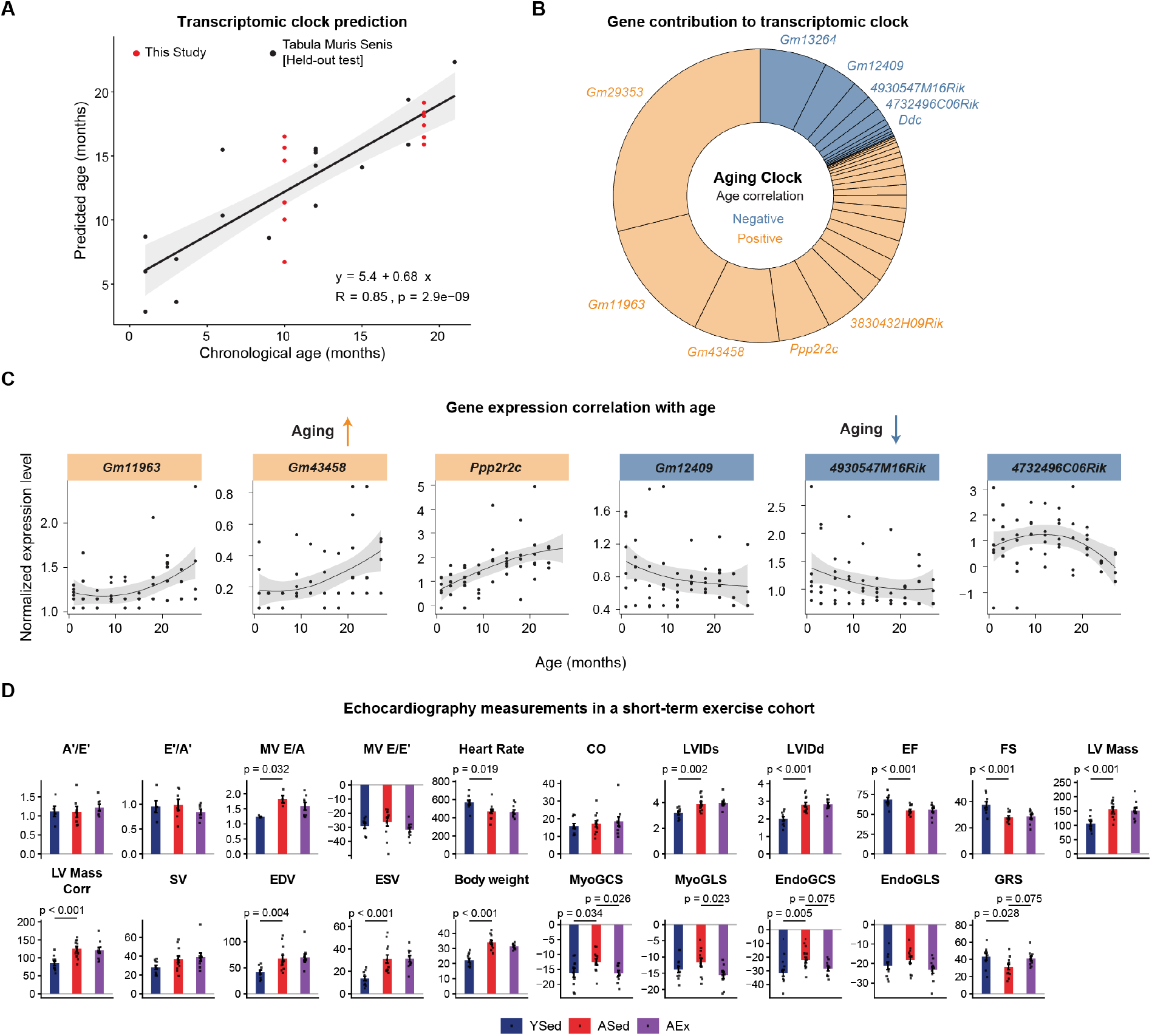
Transcriptomic ageing-clock model, feature weights and additional echocardiographic measurements in the exercise cohort. **(A)** Predicted biological age versus chronological age from the transcriptomic aging clock trained on Tabula Muris Senis cardiac RNA-seq data and applied to this study. Points represent individual animals from this study and held-out Tabula Muris Senis test samples. The fitted regression line, Pearson correlation coefficient and P value are shown. **(B)** Donut plot showing the gene weights contributing to the final ageing-clock model. Wedges are colored by positive or negative association with chronological age and selected high-weight model genes are labeled**. (C)** Bulk RNA-seq expression (normalized) of the top clock-contributing genes in each direction. Lines show smoothed regression; shaded regions show 95% CI. **(D)** Echocardiography-derived measures of cardiac function, mitral valve flow, and myocardial strain in YSed, ASed, and AEx mice from the short-term exercise cohort. For all non-flow measures and MV E/E′, n = 10 YSed, 12 ASed, and 10 AEx animals; for A′/E′ and E′/A′, n = 5, 7, and 7 animals; and for MV E/A, n = 4, 4, and 8 animals, respectively, due to the fusion of E and A waves in some samples. Bars show mean ± SEM; points represent individual animals. P values were calculated using sex-adjusted Tukey HSD comparisons.

**Supplementary Figure 7.**
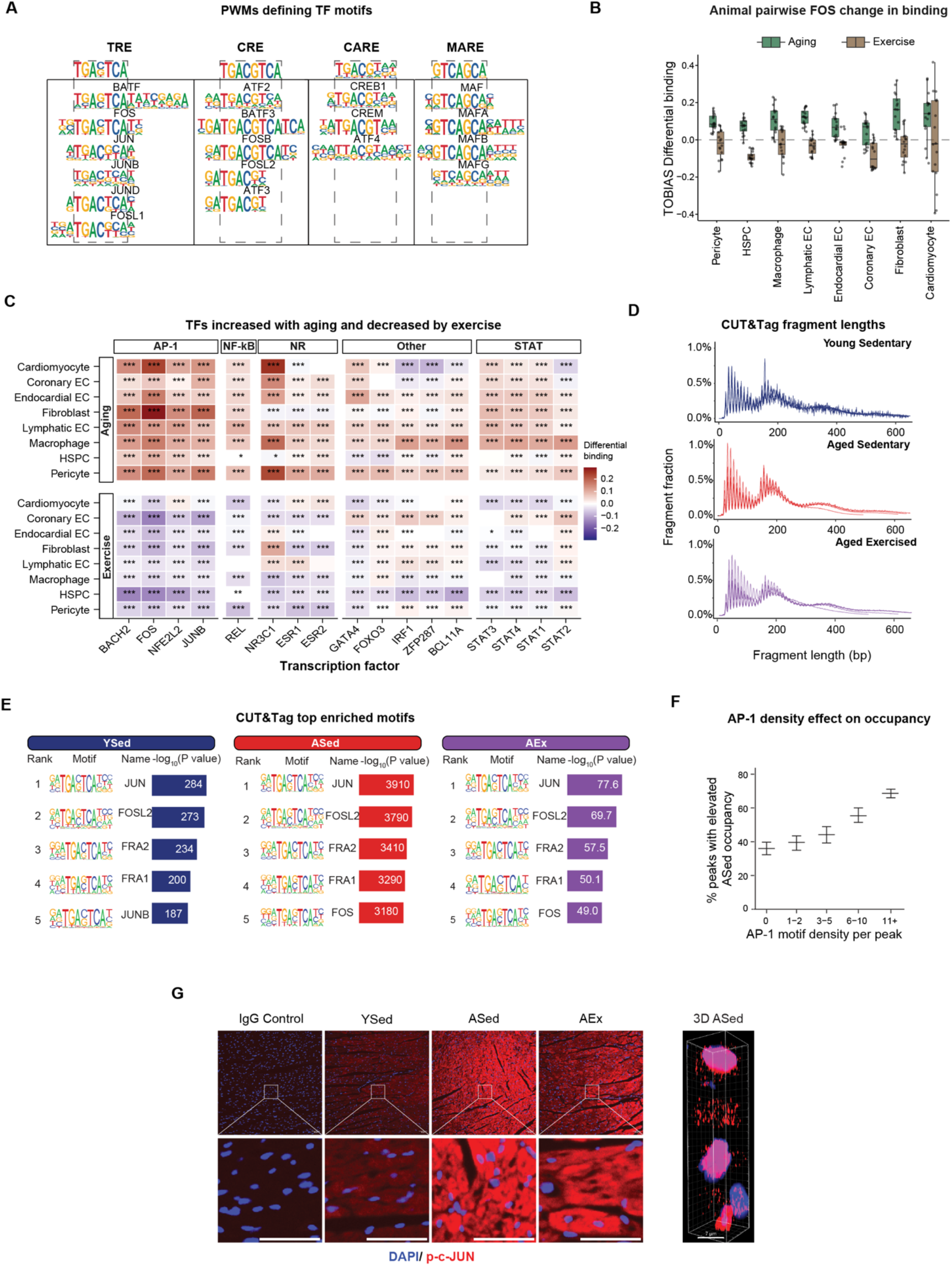
AP-1 motif architecture and opposing age-and exercise-associated TF changes. **(A)** Position weight matrices for AP-1-related motif families grouped into TRE, CRE, CARE and MARE sequence classes. Dashed boxes highlight the central core motif features used for motif-family assignment. **(B)** Boxplots showing pairwise sample-level changes in inferred c-FOS binding for the aging (ASed -YSed) and exercise (AEx - ASed) comparison, used to assess the consistency of binding changes across biological replicates. Each point represents a pairwise sample comparison. Boxplots show the median and IQR, with whiskers extending to 1.5 × IQR. **(C)** Heatmap showing TFs with increased inferred binding during aging and decreased inferred binding with exercise across cardiac cell types. TFs are grouped by AP-1, NF-κB, nuclear receptor, other and STAT classes. Color indicates TOBIAS differential binding. \**P* < 0.05, \*\**P* < 0.01 and \*\*\**P* < 0.001. **(D)** Fragment-length distributions for c-FOS CUT&Tag libraries from YSed, ASed and AEx samples. **(E)** Top enriched HOMER motifs across YSed, ASed and AEx conditions. Motif tables show rank, motif logo, assigned motif name and -log_10_(P value). **(F)** Percentage of peaks with higher c-FOS occupancy in aged sedentary relative to young sedentary hearts based on the number of AP-1 motifs per peak. Error bars show SEM; n = 673, 524, 422, 536 and 1,300 for peaks containing 0, 1–2, 3–5, 6–10 and 11+ AP-1 motifs, respectively. **(G)** Representative cardiac immunofluorescence images showing DAPI and p-c-JUN staining in YSed, ASed and AEx mice, with higher-magnification views of boxed regions. Scale bar, 50 μm (left). Three-dimensional reconstruction of z-stack confocal images showing p-c-JUN and DAPI staining in the ASed group (right).

**Supplementary Figure 8.**
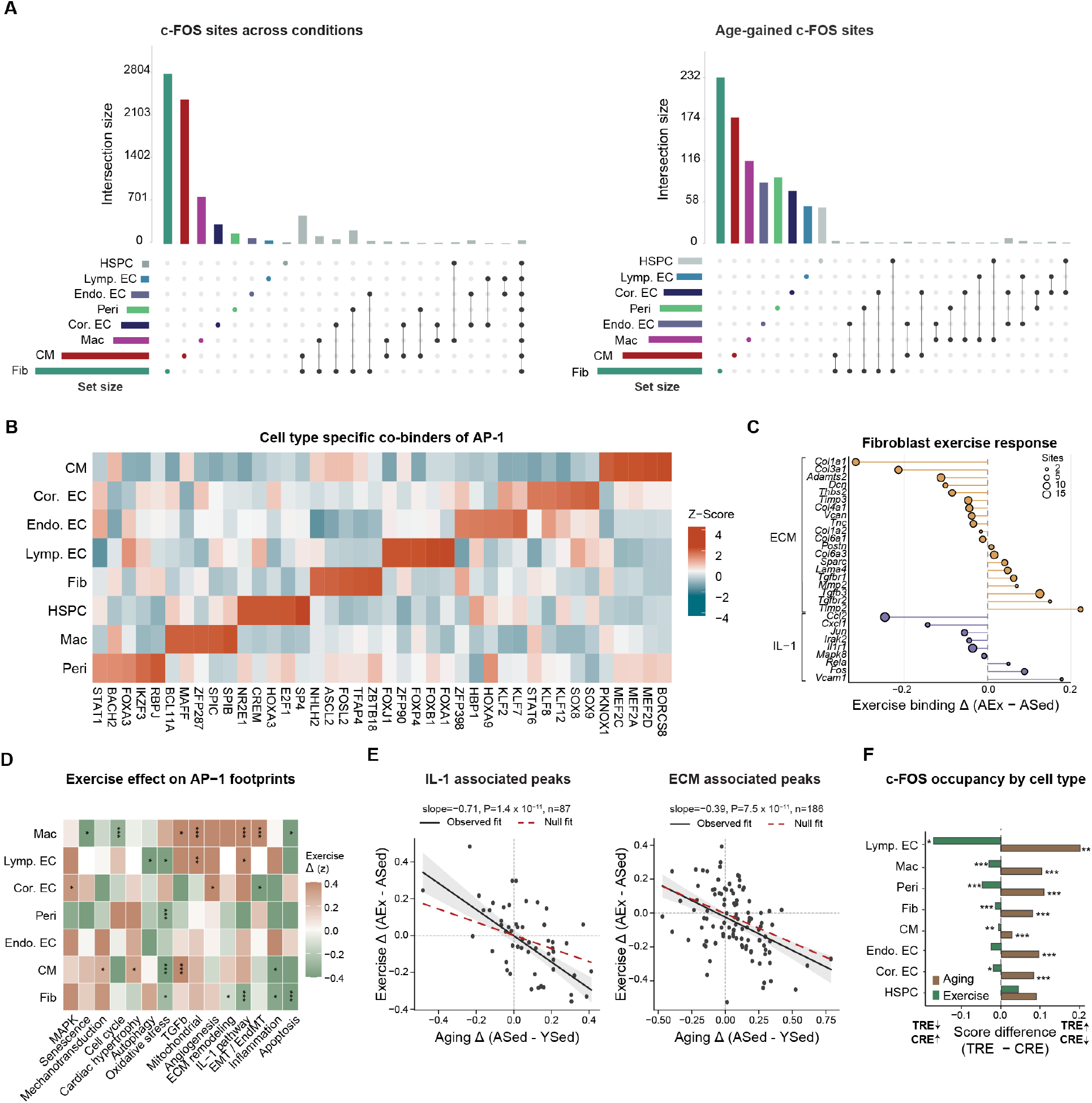
Cell-type overlap, co-binding context of c-FOS sites and exercise-associated AP-1 footprint changes across curated gene sets. **(A)** UpSet plots showing the overlap of c-FOS bound sites across major cardiac cell types for all detected c-FOS sites (left) and age-gained c-FOS sites (right). Bar heights indicate intersection sizes and side bars indicate cell-type set sizes. **(B)** Heatmap of the top five cell-type-specific motifs co-occurring near bound c-FOS sites. Motifs located within 50 bp of a bound AP-1 motif were quantified, normalized across cell types and transformed to z-score. Columns represent motifs with the strongest cell-type specificity, and rows represent cardiac cell types. **(C)** AP-1 footprinted sites located within 200 kb of ECM- and IL-1-module genes in fibroblasts. X-axis denotes the mean exercise-associated TOBIAS binding change (AEx – ASed). **(D)** Heatmap showing exercise-associated changes in AP-1 footprint activity across curated cardiac gene modules and cell types. Values indicate the mean exercise effect (AEx - ASed) and are z-scored within each module. **(E)** Relationship between ageing- and exercise-associated TOBIAS effects at AP-1 sites linked to IL-1 module genes (left) and ECM module genes (right). Regression slopes, 95% CI, and P values are shown. Dashed red line denotes the mean expected age–exercise slope from 1,000 random AP-1-site sets matched to the pathway set size. **(F)** Difference in mean TOBIAS binding-score changes between TRE- and CRE-containing motifs during aging and exercise across cell types.

**Supplementary Figure 9.**
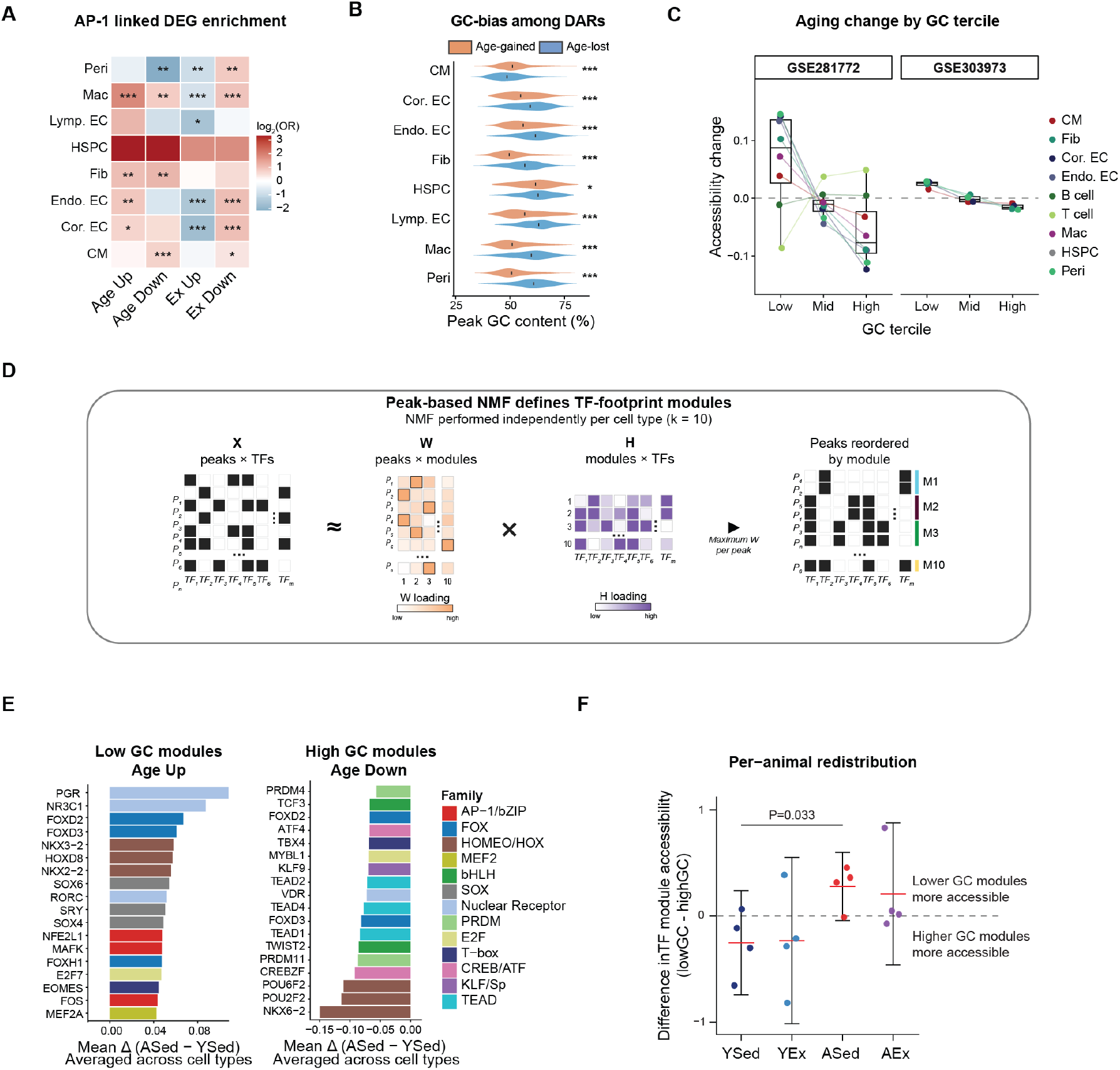
GC associated changes in accessibility and TF modules. **(A)** Enrichment of DEGs linked to AP-1 binding sites based on co-accessibility across cell types, relative to peaks without an AP-1 binding site for each cell type. Wald z-test; *P < 0.05; **P < 0.01; ***P < 0.001. **(B)** Distribution of peak GC-content for nominally age-responsive regions (P < 0.1), classified as age-gained or age-lost according to the direction of the log_2_FC. Black vertical bar indicates the median GC. (Wilcoxon rank-sum tests *FDR < 0.05, **FDR < 0.01 and ***FDR < 0.001). **(C)** Boxplots show cell-type-centred mean old-minus-young accessibility changes across GC terciles for an independent aging cohort Yucel et al.^25^ (left, 28 versus 4 months; n = 2 mice per age) assessed using matched cell-type-specific peak sets (left) and the global peak set (right). Boxplots show the median and IQR, with whiskers extending to 1.5 × IQR, points and lines denote individual cell types. Wilcoxon signed-rank test. **(D)** Schematic of the peak-based NMF performed independently for each cell type using inferred TF binding profiles. **(E)** Mean age-associated TF binding change (ASed - YSed) for individual TFs, averaged across all cell types and all chromatin modules in which the TF is scored in low- and high-GC modules. Each panel shows the top TFs ranked by absolute mean change with a maximum of three representatives per TF family. **(F)** Per-animal redistribution of TF module accessibility, calculated as the mean standardized accessibility of Low-GC modules minus the mean standardized activity of High-GC modules. Groups were compared using a Welch’s t-test.

**Supplementary Figure 10.**
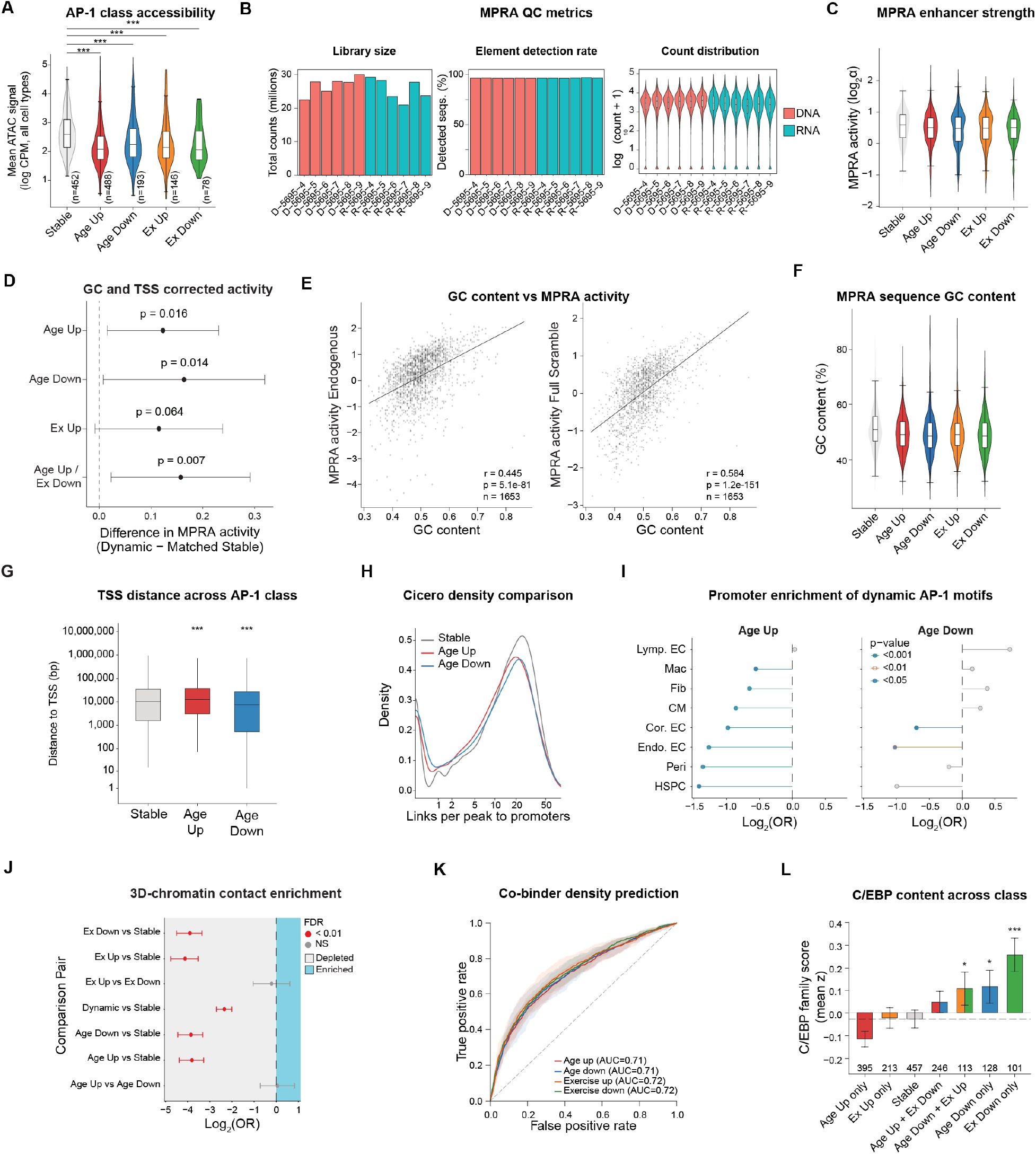
Chromatin context and MPRA quality control for dynamic c-FOS sites. **(A)** Mean snATAC-seq accessibility of MPRA-tested sequences stratified by binding category. **(B)** MPRA library quality-control metrics, including DNA and RNA library size, element detection rate and raw count distributions across replicate libraries. **(C)** Endogenous MPRA activity, measured as log₂(α) stratified by binding class. **(D)** GC- and TSS-distance-matched MPRA activity comparison between dynamic and stable sequences. Exact P values from Wilcoxon rank-sum tests. **(E)** Relationship between GC content and MPRA activity for fully scrambled sequences and endogenous sequences. Linear regression lines, correlation coefficients, P values and number of tested sequences are shown. **(F)** GC content of MPRA-tested sequences by binding category. **(G)** Distance from stable, age-gained and age-lost c-FOS sites to the nearest TSS. Wilcoxon rank-sum test, ***P < 0.001. **(H)** Density distribution of the number of Cicero links connecting stable, age-up and age-down c-FOS peaks to promoters. **(I)** Enrichment or depletion of promoter-overlapping c-FOS sites among age-up and age-down classes relative to stable c-FOS sites across cell types. Points show log_2_OR from Fisher’s exact tests and are colored by P value category. **(J)** Enrichment of dynamic c-FOS-bound regions overlapping cardiomyocyte Hi-C anchors^88^, relative to other classes of c-FOS sites. Points show log_2_OR from Fisher’s exact tests. **(K)** Receiver operating characteristic curves for an XGBoost classifier trained on motif density within 50 bp of c-FOS footprints to distinguish dynamic from stable c-FOS binding sites across four contrasts (age gain, age loss, exercise loss and exercise gain). Lines show the mean true positive rate across cell types; shading indicates ± 1 SD. **(L)** C/EBP-family composite score (mean z score) across mutually exclusive age- and exercise-binding classes. Error bars, chromosome-bootstrap 95% CI. Wilcoxon rank-sum test versus stable; *P < 0.05, **P<0.01, and ***P<0.001.

**Supplementary Figure 11.**
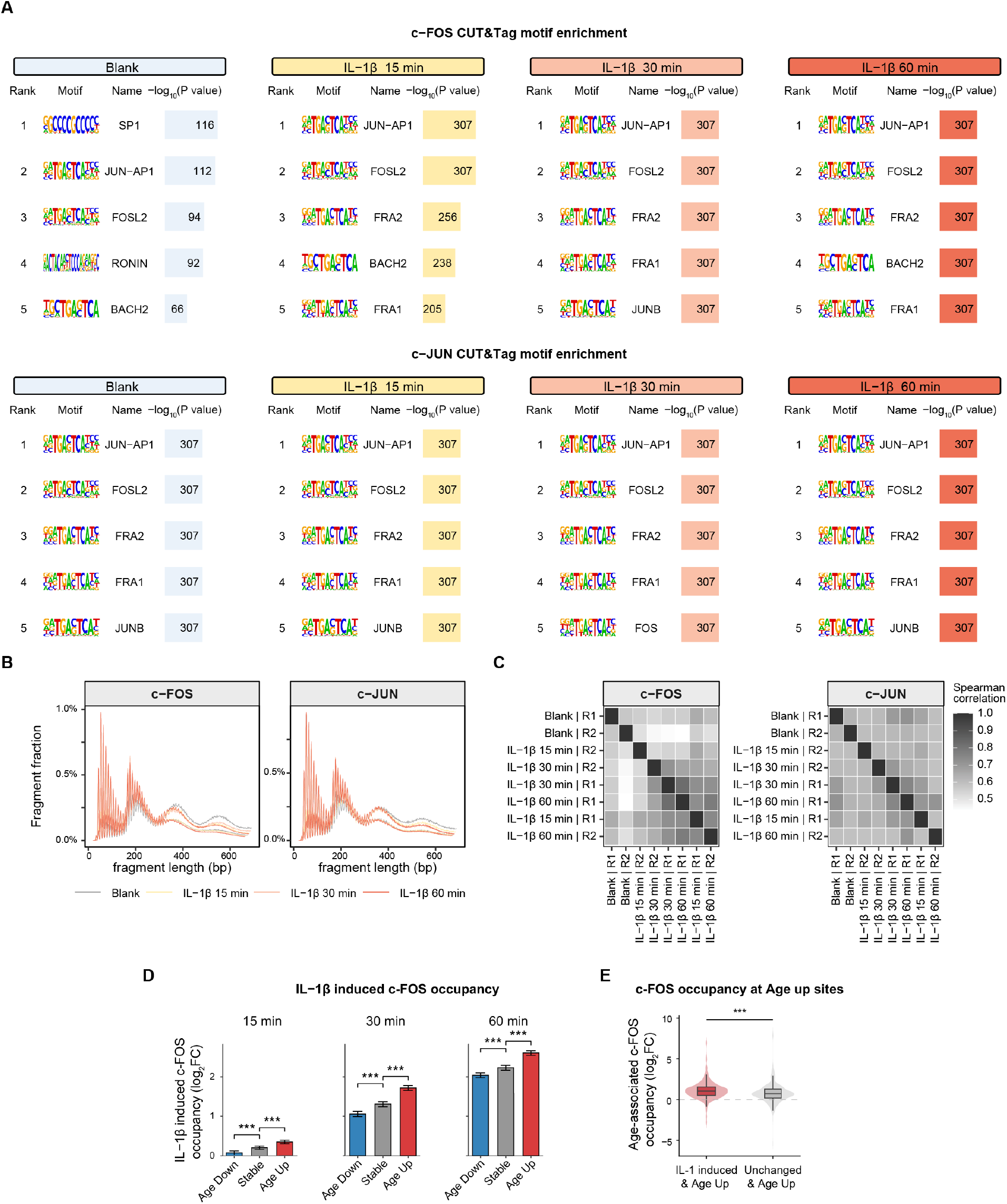
IL-1β CUT&Tag quality control in primary cardiac fibroblasts. **(A)** Top enriched HOMER motifs in c-FOS (top) and c-JUN (bottom) CUT&Tag peaks from blank controls and following 15-, 30-, and 60-minutes of IL-1β stimulation. Motif tables show motif rank, logo, assigned motif name, and −log_10_(P value). P values reaching computational limits are capped at 10^-307^. **(B)** Fragment-length distributions for c-FOS and c-JUN CUT&Tag libraries across blank controls, and IL-1β-stimulated samples collected after 15, 30, and 60 min. **(C)** Spearman correlation heatmaps showing concordance among c-FOS and c-JUN CUT&Tag replicate libraries. **(D)** IL-1β-induced c-FOS CUT&Tag changes at age-up, stable and age-down c-FOS sites (n = 1,089, 1,122, 1,138, respectively). Values show mean log_2_FC; error bars show 95% bootstrap confidence intervals from 2,000 iterations. Pairwise one-sided Wilcoxon rank-sum tests for increased induction \**P* < 0.05; \*\**P* < 0.01; \*\*\**P* < 0.001. **(E)** Age-associated change in c-FOS CUT&Tag occupancy at fibroblast age-up AP-1 sites grouped by c-FOS response to IL-1β stimulation (n = 372, 325; responsive and non-responsive sites, respectively). Boxplots: median, IQR, 1.5 × IQR whiskers. Wilcoxon rank-sum test.

**Supplementary Figure 12.**
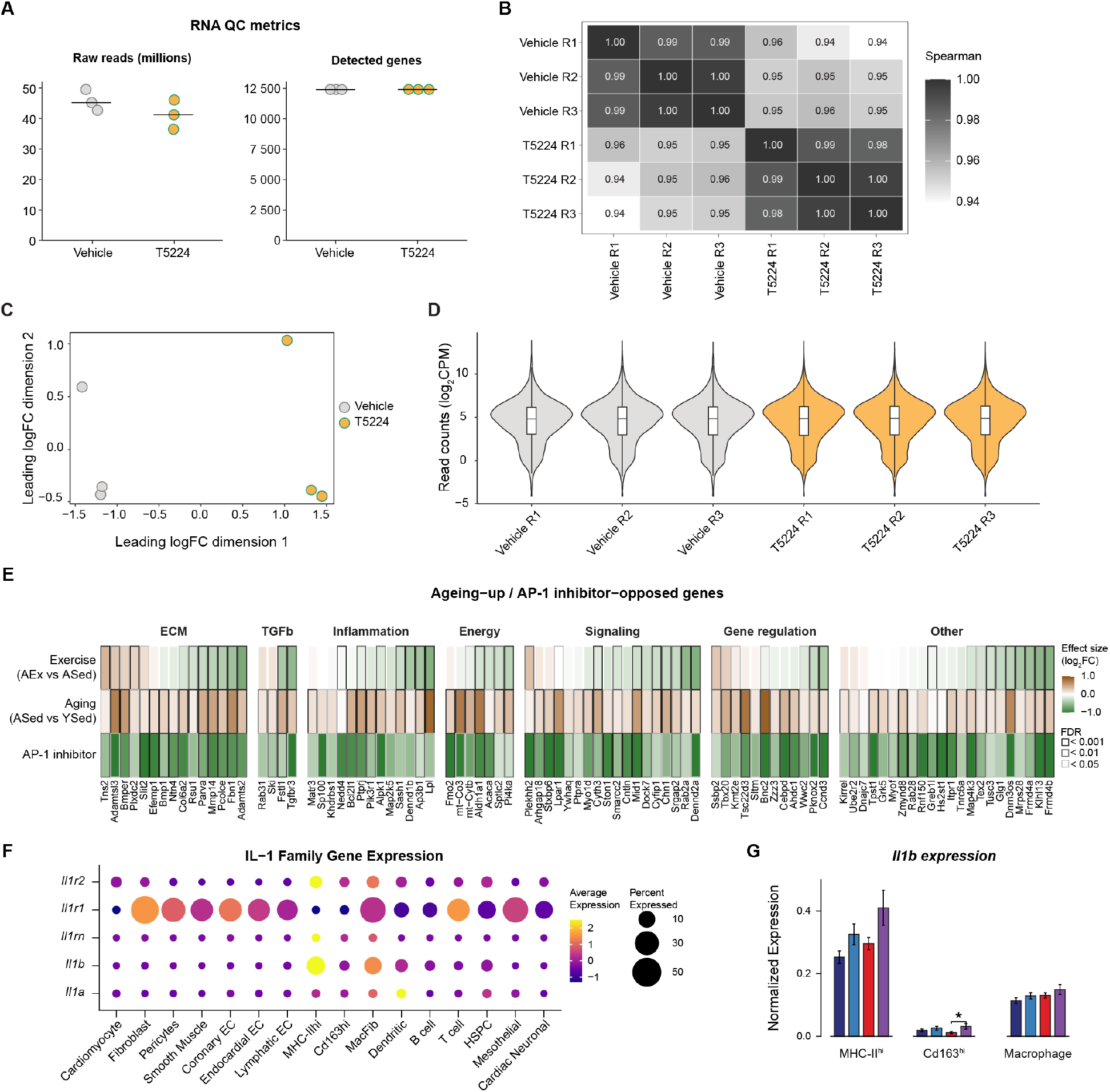
Bulk RNA-seq quality control for T-5224-treated cardiac fibroblasts and ECM gene-expression changes. **(A)** Bulk RNA-seq quality-control metrics for vehicle- and T-5224-treated primary cardiac fibroblast libraries, including the number of raw reads (left) and detected genes (right). Points represent individual replicate libraries; horizontal bars indicate the median. **(B)** Spearman correlation heatmap showing pairwise similarity among vehicle and T-5224-treated RNA-seq replicates. **(C)** Multidimensional scaling plot based on leading log-fold-change (logFC) dimensions, showing separation of vehicle and T-5224-treated samples. **(D)** Distribution of log_2_CPM values across vehicle and T-5224-treated replicate libraries. **(E)** Heatmap of log_2_FC values for genes with reduced expression after AP-1 inhibitor treatment in cardiac fibroblasts and increased with aging in vivo. Genes have been grouped into functional modules. Rows show exercise (AEx − ASed), aging (ASed − YSed) and AP-1 inhibitor versus vehicle effects. **(F)** Expression of *Il1a*, *Il1b*, *Il1rn*, *Il1r1*, and *Il1r2* across cell types. **(G)** Normalized expression of *Il1b* across cell types and conditions. Error bars show mean ± SEM across nuclei.

**Supplementary Figure 13.**
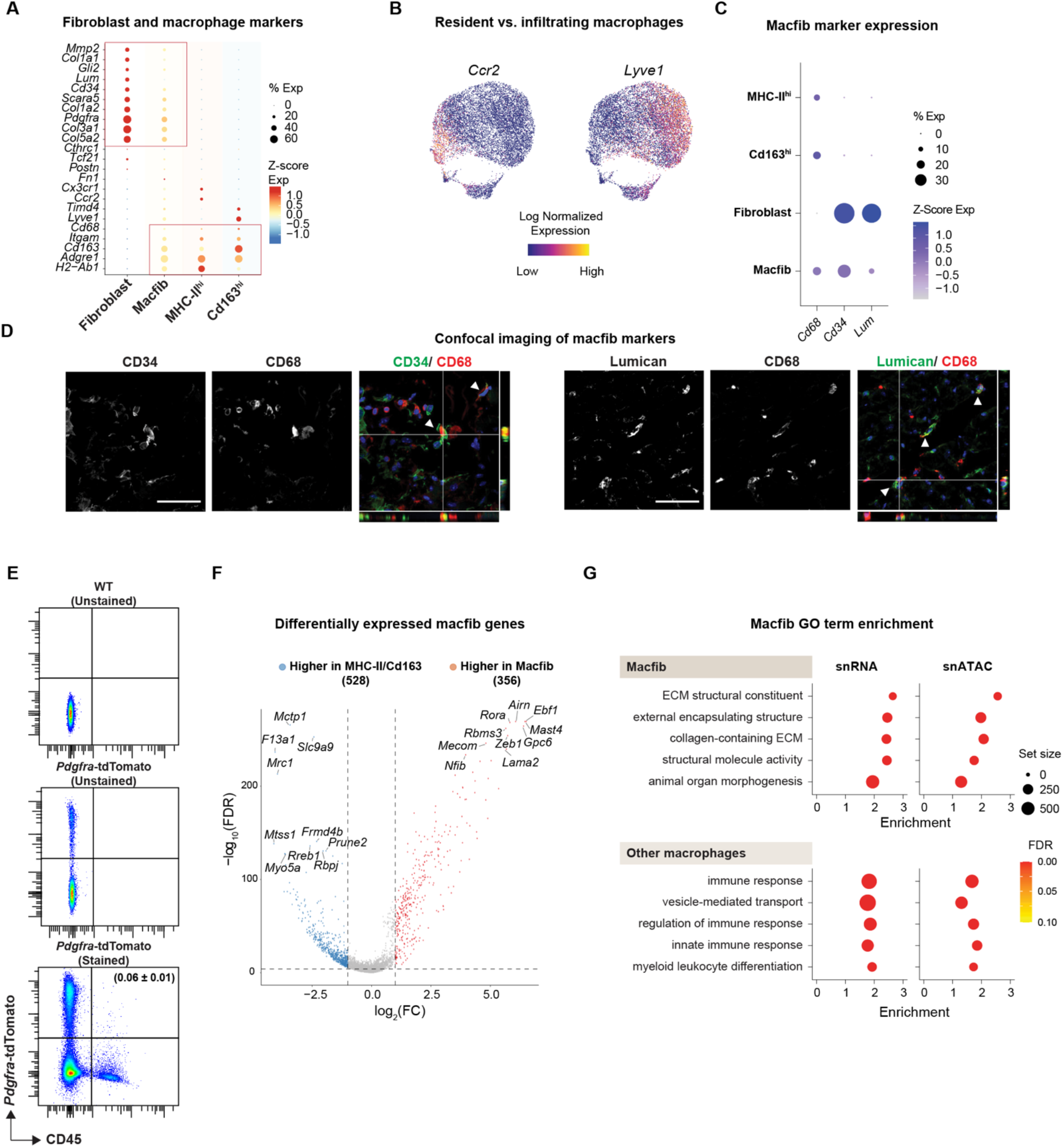
Macrophage subpopulation characterization and validation. **(A)** Expression of fibroblast and macrophage marker genes across macrophage subpopulations and fibroblasts. **(B)** Feature plots showing normalized expression of the resident macrophage marker *Lyve1* and monocyte-derived macrophage marker *Ccr2*. **(C)** Dot plot showing relative expression of *Cd68*, *Cd34* and *Lum* across MHC-IIhi macrophages, Cd163^hi^ macrophages, fibroblasts and macfib cells. Point size indicates the percentage of cells expressing each gene, and color indicates z-score-normalized expression. **(D)** Representative confocal images and orthogonal XZ and YZ projections showing CD34/CD68 and Lumican/CD68 staining in cardiac tissue. Arrowheads indicate nuclei or cells with close spatial apposition of macrophage and matrix-associated signals. Scale bar, 100 μm. **(E)** Representative flow cytometry plots showing live, single-cell gated Pdgfra lineage-traced (tdTomato⁺) cells analysed for tdTomato and CD45 expression. CD45⁺ tdTomato⁺ cells were identified as macfibs. Number indicates the frequency of macfibs within the live, single-cell gated total interstitial cell population. Data are presented as mean ± s.d. from three biological replicates (n = 3). **(F)** Volcano plot showing differential gene expression between macfib cells and MHC-II^hi^/Cd163^hi^ macrophages. Genes with higher expression in macfib cells or MHC-II^hi^/Cd163^hi^ macrophages are highlighted, and top marker genes are labeled. **(G)** Top enriched and depleted GO terms in macfib cells relative to macrophages. For snRNA-seq: preranked GSEA^117^ was performed on differential-expression log_2_FC, shown as NES. For snATAC-seq, GREAT^72,73^ was performed on differentially accessible peaks (P < 0.005 and |log_2_FC| > 1), shown as fold enrichment. Dot size indicates gene-set size.

**Supplementary Figure 14.**
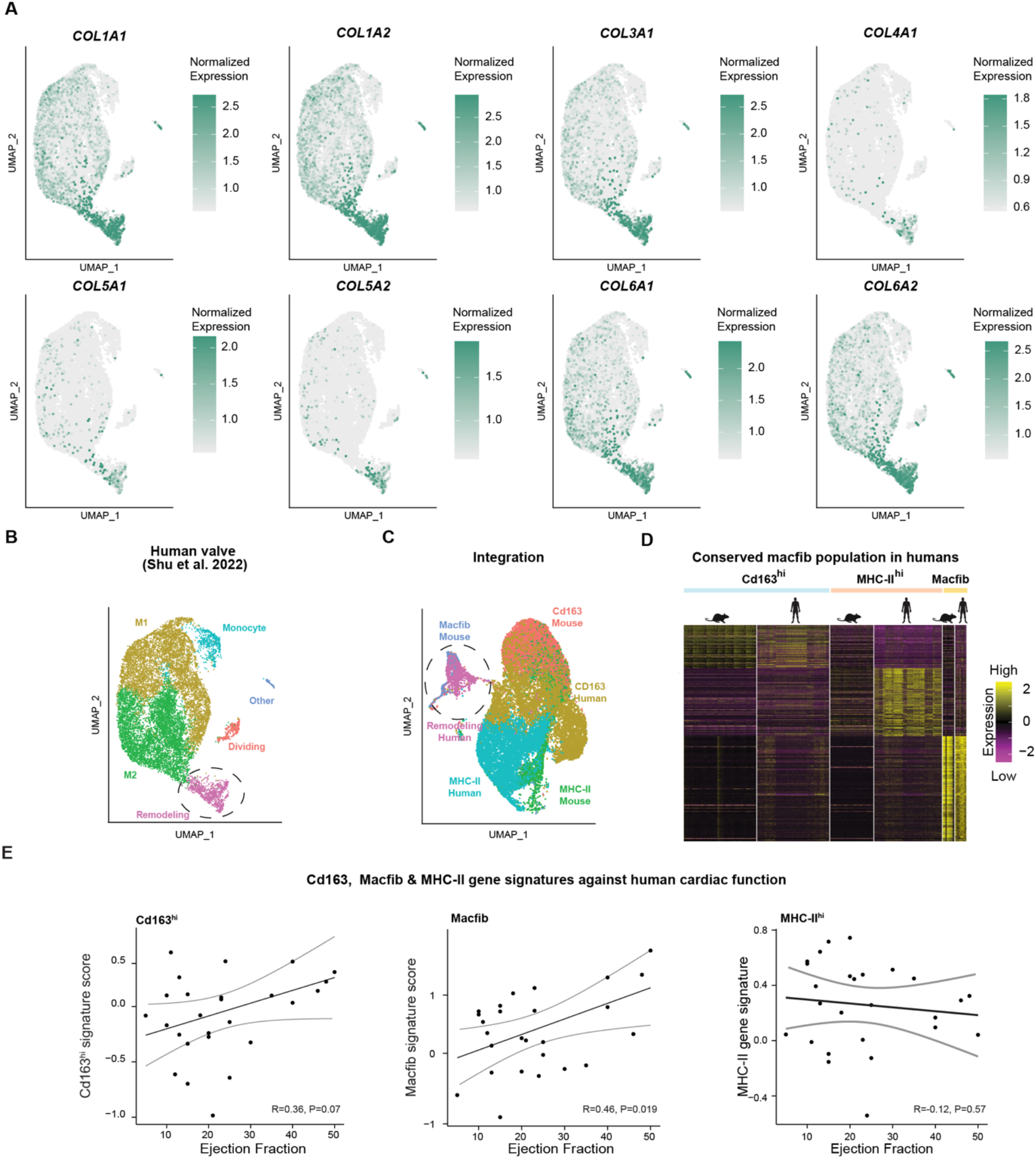
Annotation of a macfib-like population in human cardiac macrophages. **(A)** UMAP feature plots showing normalized expression of collagen genes in the human cardiac macrophage dataset, including *COL1A1*, *COL1A2*, *COL3A1*, *COL4A1*, *COL5A1*, *COL5A2*, *COL6A1* and *COL6A2*. **(B)** UMAP embedding of human cardiac macrophage (n samples = 7, n nuclei = 14,297) states annotated using original labels as dividing, M1, M2, monocyte, remodelling and other macrophages. Dashed outline identifies the macfib population. **(C)** Integrated mouse–human macrophage embedding after orthologue mapping, colored by mouse Cd163^hi^, MHC-II^hi^ and macfib and human CD163^hi^, MHC-II^hi^ and remodelling macrophage states. Dashed outline identifies the macfib population. **(D)** Cross-species comparison of mouse macfib and human cardiac remodelling macrophages^95^. Heatmap showing conserved marker genes after ortholog mapping and integration. Markers were defined using a Wilcoxon rank-sum test (P < 0.001 and log_2_FC > 0.5). **(E)** Association between Cd163^hi^, MacFib and MHC-II^hi^ human macrophage-state signatures and ejection fraction in control and LVAD patient samples^96^. Signature scores were calculated from myeloid pseudobulk profiles. Lines indicate linear regression with 95% CI; Pearson’s r and P value indicated.

**Supplementary Figure 15.**
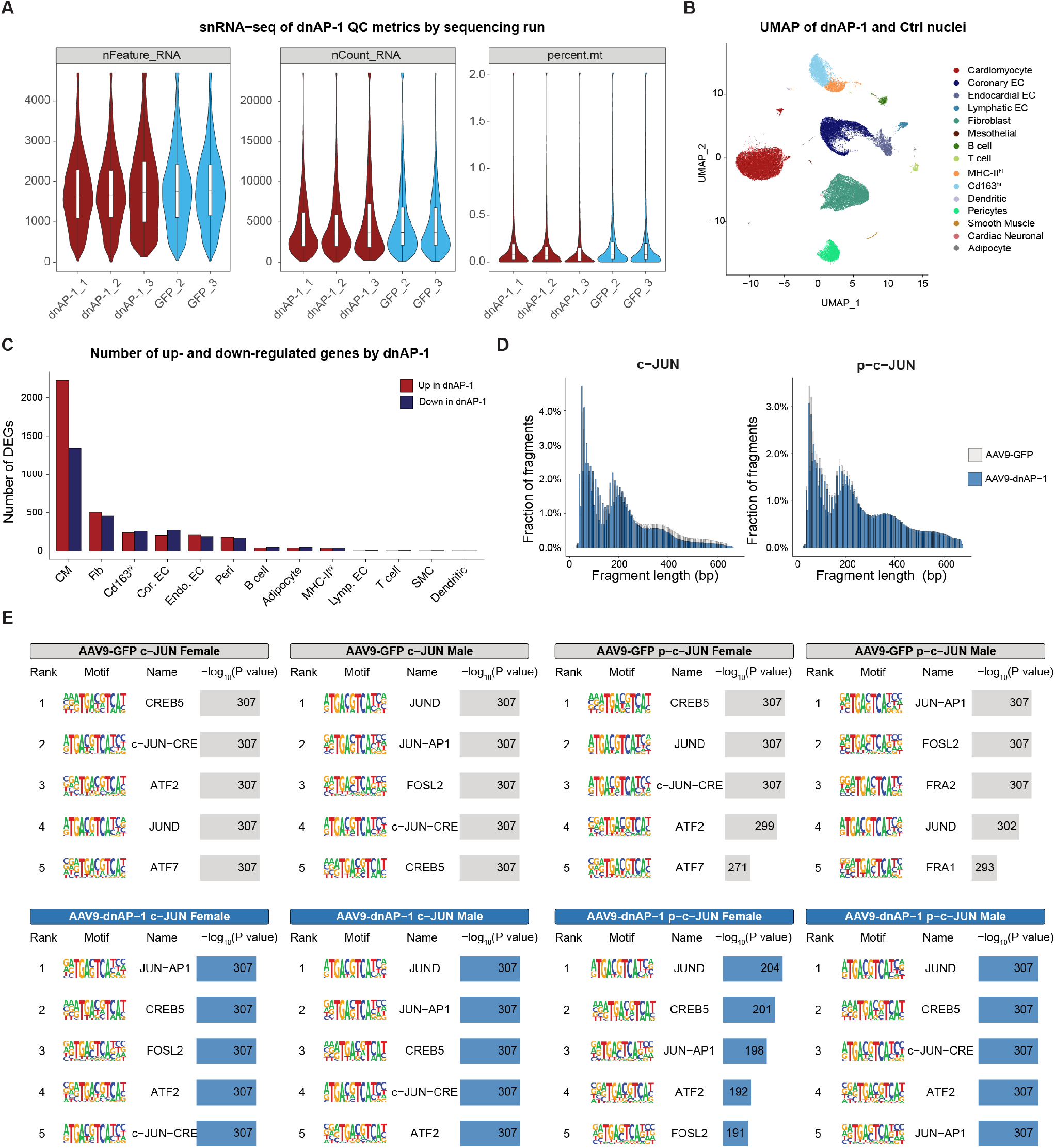
Transcriptomic and chromatin responses to in vivo AP-1 inhibition. **(A)** snRNA-seq quality-control metrics across samples, including the number of detected genes, total RNA counts and mitochondrial read percentage. **(B)** UMAP embedding of the snRNA-seq data colored by cell type. **(C)** Number of significant differentially upregulated or downregulated genes identified in each cell type following dnAP-1-treatment relative to control hearts (FDR < 0.05 and |log_2_FC| > 0.585). **(D)** Fragment-length distributions for c-JUN and p-c-JUN CUT&Tag libraries generated from aged mice receiving AAV9-GFP control or AAV9-dnAP-1. **(E)** Top enriched HOMER motifs identified in c-JUN and p-c-JUN CUT&Tag peaks, stratified by AAV9 treatment group and sex. Motif tables show motif rank, logo, assigned motif name, and −log₁₀(P value). P values reaching computational limits are capped at 10^-307^.

**Supplementary Figure 16.**
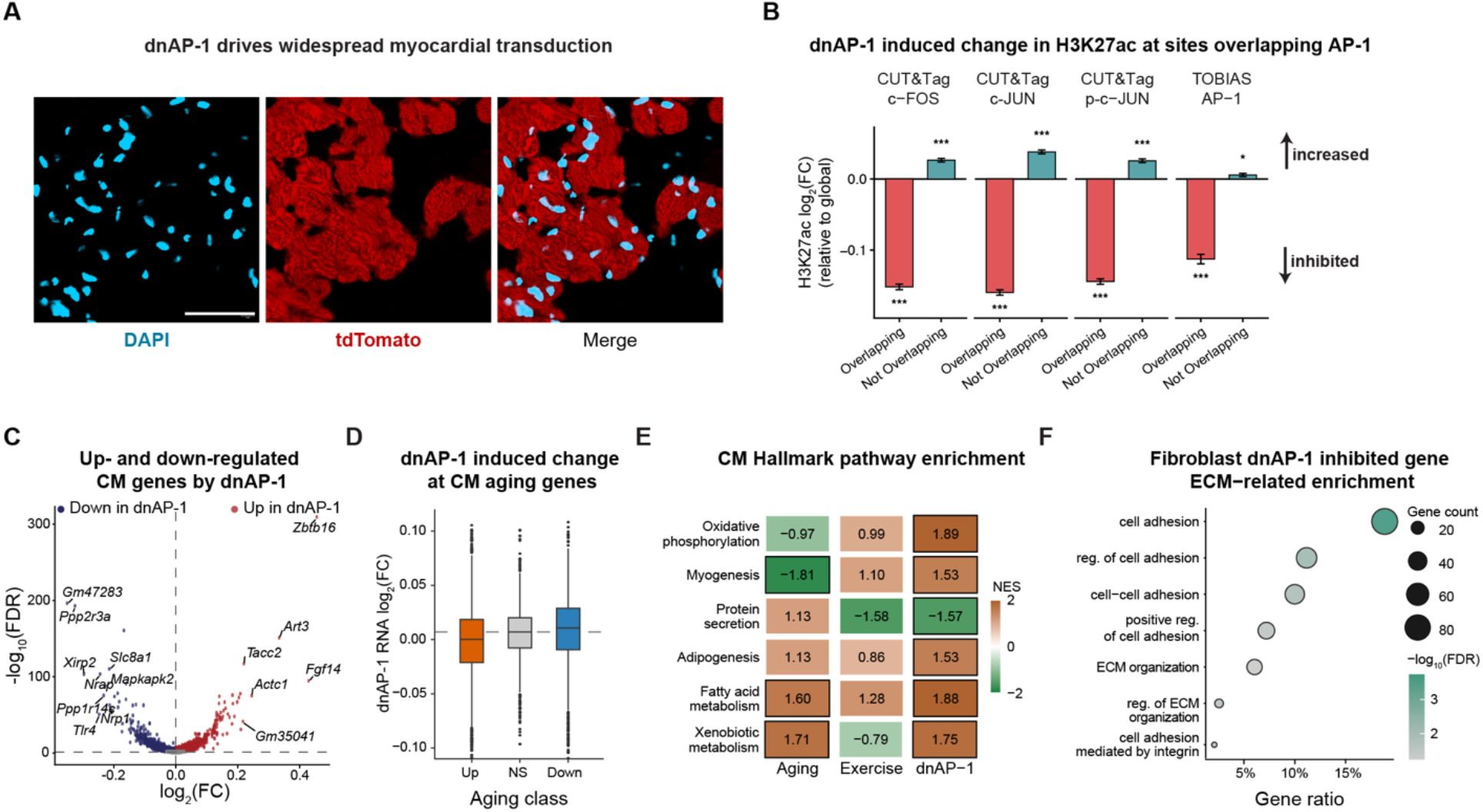
In vivo dnAP-1 target engagement and transcriptional responses. (A) Representative confocal images of heart sections from aged mice five weeks after administration of AAV9–cTnT– dnAP-1–tdTomato (n = 4 mice). Scale bars, 50 µm. **(B)** Mean H3K27ac log_2_FC after dnAP-1 expression, relative to the genome-wide mean, at peaks overlapping or not overlapping c-FOS, c-JUN, p-c-JUN CUT&Tag peaks or TOBIAS-inferred cardiomyocyte AP-1-binding sites. Bars show mean ± SEM. Two-sample Welch’s t-test; ***P < 0.001. **(C)** Volcano plot of differentially expressed genes in cardiomyocytes following dnAP-1 treatment. Significant upregulated genes are shown in red and downregulated genes in blue. **(D)** snRNA-seq log_2_FC after dnAP-1 treatment for cardiomyocyte genes classified as age-up, age-non significant (NS), or age-down. Boxes show the median and IQR; whiskers show 1.5× IQR. Dashed line indicates the median dnAP-1 response of age-NS genes. **(E)** Hallmark GSEA of cardiomyocyte gene expression following ageing, exercise and dnAP-1 treatment. Significant terms (FDR < 0.1) from the dnAP-1 experiment were selected. Terms with FDR < 0.1 are outlined in black. **(F)** Dot plot showing significant ECM- and adhesion-related GO terms enriched (FDR < 0.1) among dnAP-1-responsive fibroblast genes relative to all tested genes.

**Supplementary Figure 17.**
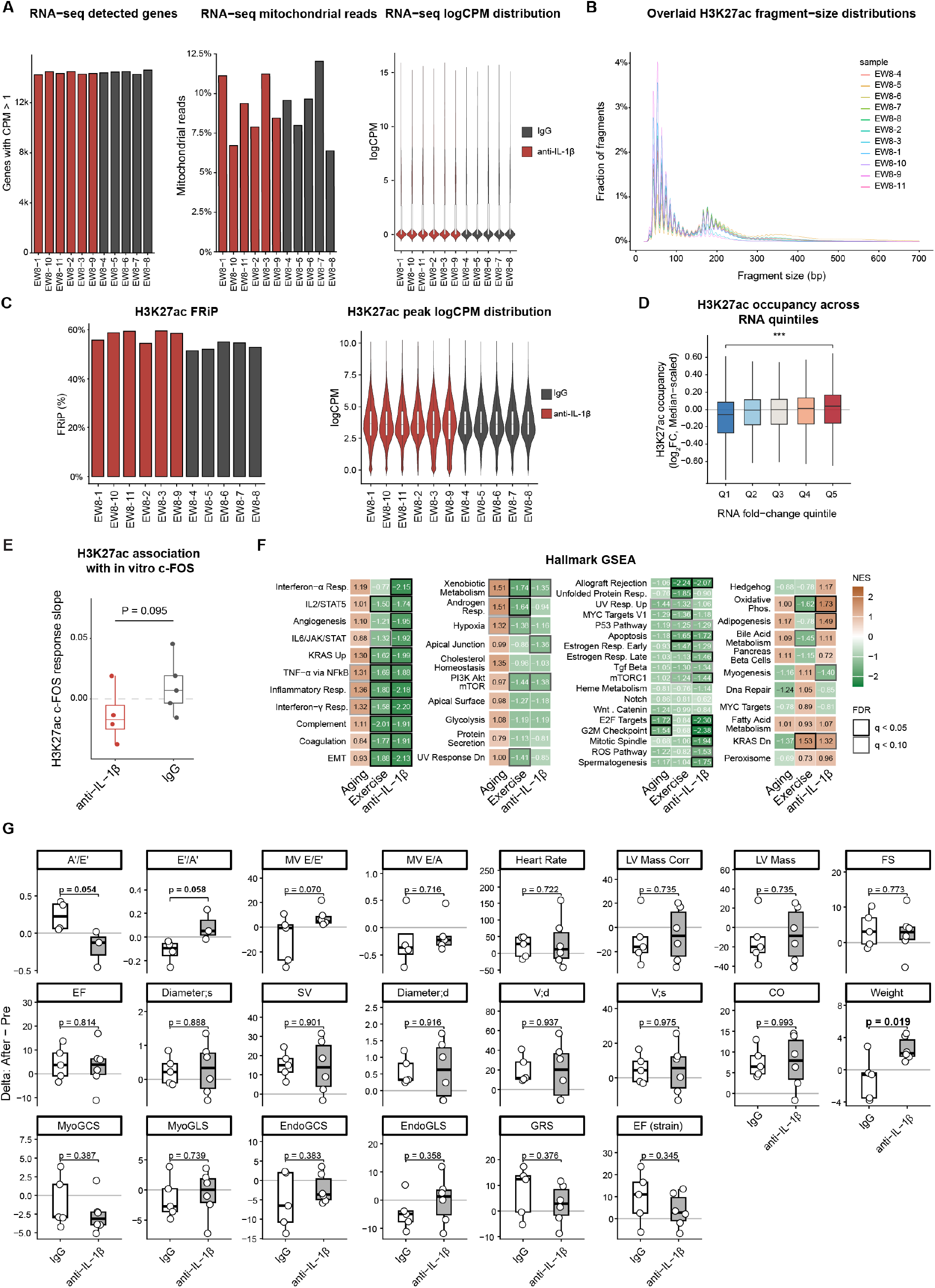
Bulk RNA-seq and H3K27ac CUT&Tag QC metrics and physiological measurements after in vivo IL-1β blockade. **(A)** RNA-seq quality-control metrics, showing the number of genes detected at CPM > 1 (left), the percentage of mitochondrial reads (middle), and gene-level logCPM values (right). **(B)** Overlaid H3K27ac CUT&Tag fragment-size distributions for individual samples. **(C)** H3K27ac CUT&Tag quality-control metrics, showing the fraction of mapped reads in peaks (FRiP) (left) and peak-level logCPM values (right). **(D)** Distribution of median-scaled H3K27ac log_2_FC across RNA-seq fold-change quintiles; boxplots show the median and IQR, with whiskers extending to 1.5 × the IQR. (Wilcoxon rank-sum test; FDR = 1.08 × 10^-33^). **(E)** Per-animal associations between the 60-min in vitro IL-1β-induced c-FOS response and in vivo H3K27ac signal at TOBIAS defined age-up AP-1-binding sites. Response slope values indicate the change in H3K27ac log_2_CPM per 1-standard-deviation increase in in vitro c-FOS responsiveness. Each point represents one animal; boxplots show the median, IQR, and whiskers extending to 1.5 × IQR. Exact one-sided label-permutation P values are shown. **(F)** Hallmark GSEA across the anti-IL-1β, aging and exercise contrasts. **(G)** Changes in echocardiographic and related physiological measurements, calculated as end-of-study minus baseline values, comparing IgG and anti-IL-1β groups. Points represent individual animals, boxplots show median, IQR and 1.5 × IQR whiskers; Welch’s t-test nominal P values are shown. A′/E′, late-to-early diastolic tissue Doppler velocity ratio; E′/A′, early-to-late diastolic tissue Doppler velocity ratio; MV E/E′, mitral valve early filling velocity to early diastolic tissue velocity ratio; MV E/A, mitral valve early-to-late filling velocity ratio; LV Mass Corr, corrected left ventricular mass; LV Mass, left ventricular mass; FS, fractional shortening; EF, ejection fraction; SV, stroke volume; V;d, end-diastolic volume; V;s, end-systolic volume; CO, cardiac output; MyoGCS, myocardial global circumferential strain; MyoGLS, myocardial global longitudinal strain; EndoGCS, endocardial global circumferential strain; EndoGLS, endocardial global longitudinal strain; GRS, global radial strain.

## Supplementary Methods and Results

### Per-nucleus sex assignment

Male and female samples were multiplexed within single-nucleus sequencing runs. To deconvolve the samples, we constructed GLM classifiers (stats::glm, 80:20 training:test split; classification threshold, 0.5, binomial family)^1^ using a combination of Xist and previously defined sex-associated genes from Skaletsky et al. (2003) and Carrel and Willard (2005)^2,3^. GLM classifiers were trained on an independent lung scRNA-seq dataset generated from the same animals in which sex was known (**Figure S2I**). For snATAC-seq, gene activity scores were calculated from chromatin accessibility using Signac::GeneActivity (biotype = NULL)^4^. Because *Xist* signal was sparse in snATAC-seq, the model was restricted to the sex-associated gene sets.

### Cell type annotation and label transfer

snRNA-seq count matrices were merged in Seurat (v4.4.0)^5^ without batch correction. Raw counts were log-normalized (scale.factor = 10,000) and the 2,000 most variable genes were used for PCA, UMAP embedding and clustering (PCs = 1:15, resolution = 0.5). Marker genes were identified using Seurat::FindAllMarkers (Wilcoxon rank-sum test, logfc.threshold = 0.5, min.pct = 0.01) and clusters with shared marker profiles were merged and assigned to cell types using canonical markers (**Table S30**)^6–8^. Endothelial cell (EC) subsets were defined using *Kank1*, *Smoc1* and *Cdh11* expression for endocardial ECs; *Timp4*, *Sparcl1* and *Ly6c1* for coronary ECs; and *Ccl21a* and *Mmrn1* for lymphatic ECs (**Figure S3A-B**)^9–11^.

snATAC-seq libraries were merged without batch correction and normalized using term frequency-inverse document frequency (TF-IDF)^4^. Feature selection was performed using Signac::FindTopFeatures followed by latent semantic indexing (LSI). UMAP embedding and clustering were performed using LSI dimensions 2–45; dimension 1 was excluded because it correlated with sequencing depth. Cell types were annotated via label transfer from the snRNA-seq dataset using Seurat::FindIntegrationAnchors and Seurat::TransferData (reduction = “lsi”)^5^ and confirmed using marker gene accessibility scores (**Figure S3C–E)**.

Both modalities recovered similar cardiac lineage composition. To assess whether cellular composition contributed to downstream differences, we compared relative cell-type abundances across prespecified within-age and within-exercise contrasts. No comparison was significant after FDR correction (FDR ≥ 0.356 for RNA-seq and FDR ≥ 0.144 for ATAC-seq; **Figure S3B,E; Table S3–4**). We next assessed whether the number of nuclei captured for each cell type affected snATAC-seq peak calling. As expected, cell types represented by more nuclei yielded larger cell-type-resolved peak sets, with lower abundance cell types being enriched for promoter associated peaks, with cell types of similar peak number clustering according to Jaccard similarity (**Figure S3F–H**).

### Assessment of single-nucleus batch effects

To assess batch variation across 10x sequencing libraries, we quantified run-associated variance. snRNA-seq cell-type-resolved pseudobulk expression profiles showed similar pairwise Spearman correlations within (mean r = 0.666) and across (mean r = 0.662) sequencing runs. PERMANOVA (vegan v2.6-4)^12^ tests on Spearman-distance matrices, adjusted for sex and experimental condition, found no evidence of run-associated variance (R^2^ = 0.233, P = 0.463). snATAC-seq pseudobulk profiles showed comparable agreement both within (r = 0.952) and across (r = 0.957) sequencing runs and lacked significant run-associated variance after adjustment for sex and condition (R^2^ = 0.182, P = 0.052). Together, these diagnostics support merged downstream analysis without global batch correction.

### Fixed-effects, mixed-effects, and pseudobulk differential expression analyses

To assess the robustness of the primary fixed-effects analysis, we compared its effect estimates and statistical significance with those obtained using mixed-effects and pseudobulk models that treated the animal as the biological replicate^13,14^.

Mixed-effects models were fitted separately for each cell type using MAST (v1.24.1; method = “glmer”)^15,16^. Sex, cellular detection rate (CDR) and experimental condition were included as fixed effects, with animal identity modelled as a random intercept:

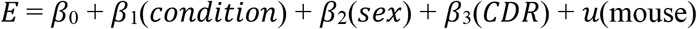

The corresponding fixed-effects model omitted the mouse-level random intercept:

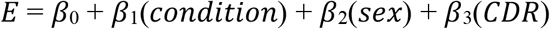

Empirical Bayes variance moderation, which was applied in the fixed-effects analysis, was disabled for the mixed-effects models because it is not supported by the MAST mixed-model implementation^17^. The models therefore differed in both their treatment of within-mouse dependence and their use of variance moderation.

Fixed-and mixed-effects models were compared using per-gene standard errors extracted from the continuous component variance-covariance matrix of each fitted model. Between-model differences in uncertainty (DEFF) were quantified using a model-based variance ratio^14^:

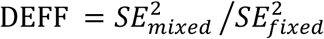

Values near 1 indicated comparable uncertainty; values greater than 1 indicated greater uncertainty under the mixed-effects model. Because the fixed-and mixed-effects models differed in both their treatment of within-animal dependence and their use of empirical Bayes variance moderation, this ratio was interpreted as an overall measure of between-model uncertainty rather than a direct estimate of variance inflation due to pseudoreplication. Fits that failed to converge or returned non-finite or extreme values (DEFF < 0.5 or DEFF > 500) were excluded.

Across 69,837 gene–contrast pairs, fixed-and mixed-effects models produced nearly identical log_2_FC estimates (median Pearson r = 0.995; median absolute difference = 0.017; directional concordance = 96.8%; **Figure S4, Table S34**). Explicitly modelling animal-level variation had little effect on the estimated direction or magnitude. The models differed primarily in statistical sensitivity: the fixed-effects model identified 4,555 significant gene–contrast pairs (FDR < 0.05 and |log_2_FC| > 0.5), compared with 780 under the mixed-effects model. This difference reflects adjustment for within-mouse dependence and the absence of empirical Bayes variance moderation in the mixed model. Consistent with the small number of biological replicates per condition (n = 2–4), 53.7% of mixed-model fits estimated the mouse-level variance component as zero.

Uncertainty differed most in cell types with many nuclei per animal, in which within-mouse dependence has the greatest potential to deflate fixed-effects standard errors. In cardiomyocytes (median, 2,576 nuclei per animal), the median DEFF was 11.6, corresponding to mixed-model standard errors 3.4-fold larger than fixed-model estimates. DEFF remained near 1 in moderately represented populations, including endocardial ECs (1.114) and lymphatic ECs (0.912), and fell below 1 in sparse populations, including cardiac neuronal cells (0.733) and B cells (0.746). Thus, modelling within-mouse dependence primarily increased uncertainty in abundant cell types but had little effect on fold-change estimates.

Pseudobulk differential expression analysis was performed using DESeq2 (v1.38.3)^18^. Counts were aggregated by mouse and cell type, and negative-binomial models included sex and experimental condition, with age and exercise effects assessed using Wald tests. Pseudobulk and fixed-effects estimates were broadly concordant (median Pearson r = 0.851; 82.4% directional agreement; **Figure S5**). DESeq2 identified fewer significant genes, as expected with only two to four biological replicates per condition. Nevertheless, gene–contrast pairs significant in the fixed-effects analysis were enriched approximately 15-fold for nominal DESeq2 significance relative to fixed-model null pairs (32.8% versus 2.2% with P < 0.05). Thus, fixed-effects signals were enriched for support in the biological-replicate analysis.

The three approaches showed strong concordance in effect directions and magnitudes but differed in statistical sensitivity. The fixed-effects model detected the most significant associations, reflecting its use of empirical Bayes variance moderation and nucleus-level analysis, whereas the mixed-effects and pseudobulk models accounted for animal-level biological replication and consequently provided more conservative significance estimates. We therefore retained fixed-effects MAST as the primary discovery framework, while using the mixed-effects and pseudobulk results to evaluate the consistency of the estimated effects.

### Ageing- and exercise-associated transcriptional opposition across cardiac cell types

To determine whether exercise modified ageing-associated transcriptional changes, MAST mixed-effect models were fitted separately for each cell type using fixed effects for age, exercise, sex, CDR, and an age-by-exercise interaction term, with mouse included as a random effect:

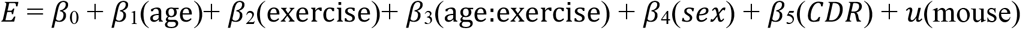

Among age-associated genes (FDR < 0.05, |log_2_FC| ≥ 0.5), interaction coefficients overwhelmingly opposed the ageing effect (167/176, one-sided exact binomial test, P = 4.00 × 10^−39^), with significant directional bias across cardiomyocytes, fibroblasts, coronary ECs, endocardial ECs, pericytes, T cells and macrophages (one-sided exact binomial tests, P < 0.05, **Table S35**). Relaxing the age-gene selection criterion to nominal P < 0.05 while retaining |log_2_FC| ≥ 0.5 did not materially alter the result: 3,974 of 4,345 interaction effects opposed the ageing direction (91.5%, P < 2.2 × 10^-308^), with significant directional bias in all 17 cell types.

### CUT&Tag protocol

For each CUT&Tag^25,26^ replicate, we isolated 50,000 nuclei and incubated in NE1 buffer (20 mM HEPES–KOH, pH 7.9; 10 mM KCl; 0.5 mM spermidine; 0.1% Triton X-100; 20% glycerol; Roche Complete Protease Inhibitor, EDTA-free) for 10 minutes at room temperature (RT). Samples were centrifuged at 500 × g for 5 minutes at RT, and the pellet was resuspended with activated concanavalin A–coated magnetic beads (Cell Signaling Technology, cat. no. 93569). Beads were captured using a magnetic stand and resuspended in antibody buffer [wash buffer (1 mL 1 M HEPES, pH 7.5; 1.5 mL 5 M NaCl; 12.5 μL 2 M spermidine; brought to a final volume of 50 mL with dH₂O; and 1 Roche Complete Protease Inhibitor EDTA-free tablet), supplemented with 2 mM EDTA, 0.1% BSA, and 0.01% digitonin] containing the primary antibody (1:50 dilution). After incubation at RT for 1 h, beads were magnetically separated, resuspended in antibody buffer (8 μL 0.5 M EDTA and 6.7 μL 30% BSA in 2 mL wash buffer) containing the secondary antibody (1:100 dilution), and incubated for 30 minutes at RT with rotation at 500 rpm. After antibody binding, beads were washed once with 300-wash buffer (20 mM HEPES, pH 7.5; 300 mM NaCl; 0.5 mM spermidine; Roche Complete Protease Inhibitor, EDTA-free) and resuspended in 300-wash buffer supplemented with 0.01% digitonin and pAG-Tn5 transposase (Cell Signaling Technology, cat. no. 79561; 1:25 dilution). Samples were incubated for 1 hour at RT with rotation at 500 rpm. Beads were then washed with 300-wash buffer while retained on a magnetic stand. For tagmentation, beads were resuspended in tagmentation buffer (300-wash buffer supplemented with 10 mM MgCl₂) and incubated for 1 hour at RT with rotation at 500 rpm. The reaction was terminated by adding EDTA, SDS, and proteinase K to final concentrations of 16 mM EDTA, 0.1% SDS, and 0.16 μg/μL proteinase K, followed by incubation at 70 °C for 10 minutes. Tagmented DNA was extracted with phenol–chloroform, precipitated with ethanol and resuspended in 21 μL TE buffer (1 mM Tris–HCl, pH 8.0; 0.1 mM EDTA).

### Validation of GC-associated ageing accessibility

Cardiac single-nucleus multiome data from Yucel et al. (GSE281772)^19^ were used to determine whether the association between peak GC content and age-associated accessibility changes was reproducible in an independent cohort. The dataset included 28-and 4-month-old animals (n = 2 per age group). Nuclei were retained following the authors’ original filtering criteria, requiring at least 800 RNA counts, 500 detected genes, and <10% mitochondrial RNA. RNA counts were library-size normalized, and cardiac cell types were assigned using canonical marker genes^6–8^. ATAC counts were then aggregated by animal and cell type to generate pseudobulk profiles.

For cell types containing more than 200 nuclei, peaks were filtered using edgeR^20^, pseudobulk profiles were normalized using the trimmed mean of M-values (TMM) method, and differential accessibility between old and young animals estimated using quasi-likelihood negative-binomial models. Peak GC fraction was calculated from the corresponding mm10 genomic sequence. Peaks were divided into cell-type-specific GC terciles, and the mean age-associated log₂FC was calculated for each tercile. To assess the robustness of this relationship, analysis was performed using both peaks overlapping cell-type accessible regions identified in our dataset and the global peak set provided by Yucel et al (**Figure S9C**).

### GC-stratified regulatory reorganization based on TF binding site modules

Ageing redistributed regulatory activity according to module sequence composition, with lower-GC modules gaining activity relative to higher-GC modules. High-GC modules (60–66% GC) lost inferred TF occupancy with age (26 of 27 modules; mean change = −0.036), whereas lower-GC modules (48– 53% GC) more often gained occupancy (16 of 27 modules; mean change = +0.020; Cohen’s d = 2.07 between GC-content terciles; **Figure 2O, Table S16-18**). The GC-dependent ageing effect was consistent across repeated measurements within animals and modules (P = 0.033; n = 8 animals; **Figure S9F**). This relationship was not explained by promoter proximity: GC content remained associated with age-related occupancy changes within both proximal and distal regulatory elements and after adjustment for distance to the transcription start site (P = 1.4 × 10^−4^). Exercise partially altered the module-level redistribution, with module-level ageing and exercise effects negatively correlated (Spearman’s ρ = −0.48, P = 7.5 × 10^−6^; ρ = −0.63 among high-GC modules, P = 6 × 10^−4^).

The magnitude and direction of age-associated change were more strongly related to module GC content than to the occupancy of any individual TF family. High-GC, age-depleted modules were enriched for KLF/SP, TEAD and ATF/CREB motifs, whereas lower-GC, age-gained modules were enriched for motifs of stress-responsive bZIP factors, including AP-1, and cardiac lineage regulators (**Figure S9E, Table S18**). These findings identify a sequence-composition-stratified redistribution that co-occurs with ageing.

### Cardiac fibroblast isolation

Fibroblasts were isolated from 5-month-old female C57BL/6J mice^27^. Hearts were rapidly excised and transferred to a 100-mm Petri dish containing sterile phosphate-buffered saline (PBS). Excess blood was removed by gentle compression with forceps. Residual aortic tissue was excised, and hearts were washed twice with 10 mL PBS. Cardiac tissue was minced into approximately 1-mm² fragments and transferred to a 70-mL specimen jar containing 6 mL collagenase solution (290 U/ml) (Collagenase Type II; Worthington, CAS 9001-12-1). Tissue fragments were gently triturated to disperse clumps and incubated at 37 °C in a shaking water bath (60 rpm) for 12 minutes, with intermittent gentle pipetting to facilitate digestion while minimizing bubble formation.

Following digestion, the supernatant was carefully collected and filtered through a 40-µm sterile cell strainer into a 50-mL conical tube. The digestion step was repeated twice more, for three rounds in total, and all supernatants were pooled. The cell suspension was centrifuged at 500 × g for 10 minutes at 4 °C, and the supernatant was discarded. The resulting pellet was gently resuspended in 2 mL red blood cell lysis buffer and incubated at room temperature for 1 minute, followed by centrifugation at 500 × g for 5 minutes. The pellet was then resuspended in 10 mL Dulbecco’s Modified Eagle Medium (DMEM, Gibco, cat. no. 11995073) supplemented with 10% fetal bovine serum (FBS, Gibco, cat. no. A5670701) and 1% penicillin–streptomycin (Gibco, cat. no. 15140122).

Cells were pre-plated onto 0.1% gelatin-coated 10-cm diameter dishes and incubated for 3 hours at 37 °C and 5% CO₂ to allow preferential fibroblast adherence. Non-adherent cells and debris were removed by aspirating the medium and washing adherent cells three to four times with pre-warmed culture medium. Fresh complete medium (10 mL) was added, and cells were maintained in a humidified incubator at 37 °C and 5% CO₂. Twenty hours after plating, cultures were gently washed twice with pre-warmed medium to remove dead cells and replenished with fresh complete medium. Cells were cultured to 80–90% confluence before downstream applications.

Primary cardiac fibroblasts were plated in 6-well plates in αMEM (α minimum essential medium; Gibco, cat. no. 12571063) supplemented with 20% FBS. At approximately 80% confluence, cells were serum-starved for 1 hour by replacing the culture medium with αMEM containing 1% FBS prior to treatment. For RNA-seq experiments, cells were allocated to one of three treatment groups: (i) vehicle control; (ii) IL-1β (10 ng/mL); and (iii) the AP-1 inhibitor T-5224 (80 µM). All groups were harvested at matched time points of 3 or 6 h, with three biological replicates per group per time point (n = 3; 18 samples in total). T-5224 was obtained from MedChemExpress (cat. no. HY-12270) and IL-1β from PeproTech (distributed by Thermo Fisher Scientific; cat. no. 200-01B-10UG). CUT&Tag was performed for c-FOS and c-JUN with two biological replicates per condition with an *E. coli* carry-over spike-in used for normalization. Total RNA was extracted using the Direct-zol™ RNA Miniprep Kit (Zymo Research, cat. no. R2052) according to the manufacturer’s instructions. RNA quality and integrity were assessed using High Sensitivity RNA ScreenTapes (Agilent Technologies; distributed by Integrated Sciences, cat. no. 5067 and 5579). Samples with DV200 ≥55% were used for downstream processing. For library preparation, 125 ng of total RNA per sample was used as input to the Illumina Stranded Total RNA Prep with Ribo-Zero Plus kit. This workflow includes depletion of cytoplasmic and mitochondrial ribosomal RNA and enables strand-specific library construction. Libraries were uniquely dual indexed, PCR amplified, normalized, and pooled prior to sequencing on an Illumina NovaSeq X Plus platform using paired-end 150 bp reads.

### Lenti-STARR-seq cloning and transfection

A total of 5,583 oligonucleotides were synthesized by Twist Bioscience, each flanked by 5′ and 3′ adaptor sequences. The pooled oligonucleotides were resuspended in TE buffer to a final concentration of 100 nM. Vector-specific overhangs were added to the oligonucleotide pool by PCR using NEBNext High Fidelity 2× PCR Master Mix (New England Biolabs, cat. no. M0541L) under the following cycling conditions: 20 cycles of 98 °C for 15 s, 65 °C for 20 s, and 72 °C for 10 s. PCR products were resolved by agarose gel electrophoresis, and the expected 216-bp band was excised and purified using the Wizard SV Gel and PCR Clean Up System (Promega, cat. no. A9281). Purified amplicons were cloned using NEBuilder HiFi DNA Assembly Master Mix (New England Biolabs cat. no. E2621S) into the EcoRI-digested pLS-mP vector^28^ (Addgene #81225), placing the candidate enhancer sequences in the 3′ untranslated region (UTR) of EGFP, such that enhancer-driven transcription of EGFP produced RNA molecules that served as their own barcodes^29^. The resulting lentiviral STARR-seq library was electroporated into ElectroMAX™ DH10B electrocompetent cells (Invitrogen, cat. no. 18290015) by electroporation in 0.1-cm cuvettes (Bio-Rad, cat. no. 1652089) at 2.0 kV on an Eppendorf. Each reaction combined 2.6 ng of assembled library DNA with 50 µL of cells, and a total of 2 electroporations were performed. Immediately after electroporation, 450 μL of prewarmed Stable Outgrowth Medium was added, and cells were recovered for 1 hour at 37 °C with agitation at 220 rpm. All electroporation reactions were pooled into 1 L of sterile LB medium containing ampicillin (100 μg/mL) and grown overnight at 37 °C at 220 rpm. Bacterial cultures were harvested by centrifugation, and plasmid DNA was purified using the PureLink™ HiPure Plasmid Maxiprep Kit (Invitrogen, cat. no. K210007).

HEK293T cells were cultured in 10-cm diameter dishes in Dulbecco’s Modified Eagle’s Medium supplemented with 10% heat-inactivated FBS and grown to approximately 70% confluence within 24 hours prior to transfection. Each dish received 10 μg STARR-seq library plasmid, 6.5 μg psPAX2 (Addgene #12260), and 3.5 μg pMD2.G (Addgene #12259). Plasmid DNA was mixed with 60 μg polyethylenimine (PEI; 1 μg/μL; Polysciences, cat. no. 23966) in a total volume of 1 mL Opti-MEM (Invitrogen, cat. no. 31985062), incubated for 15 minutes at room temperature to allow complex formation, and added dropwise to the cells with gentle swirling. Four hours after transfection, cells were washed once with DPBS and cultured in DMEM supplemented with 5% heat-inactivated FBS. Viral supernatant was collected 48 hours after transfection and stored at 4 °C. A second collection was performed at 72 hours after transfection, and supernatants were pooled. The pooled viral supernatant was filtered through a 0.45 µm PES filter and concentrated using PEG 8000. Briefly, viral supernatant was mixed with a 4× PEG8000/NaCl solution (final ∼8% PEG, 0.6 M NaCl), incubated overnight at 4 °C, and centrifuged to pellet viral particles, followed by resuspension in PBS in one-tenth of the original volume. Concentrated lentiviral particles were resuspended and snap-frozen in liquid nitrogen and stored at −80 °C until further use.

### Sequence modelling and in silico perturbations

We trained a linear probe on frozen representations from AlphaGenome, a ∼450-million-parameter sequence-to-function model comprising a convolutional encoder, a transformer and task-specific decoders^30^. The frozen ∼90-million-parameter convolutional encoder accepts up to ∼1 Mb of one-hot-encoded DNA sequence and returns representations of shape (ceil(L/128), 1536), where L is the length of the input DNA sequence. The frozen encoder provided fixed representations for the downstream probe^31^. The linear probe accepts representations of shape (2, 1536) and predicts scalar MPRA activity as log_2_(RNA/DNA). The resulting AlphaGenome-MPRA model takes one-hot-encoded 200-bp constructs as input, comprising a 170-bp sequence flanked by adapters and predicts a single activity score.

The mouse AP-1 MPRA assayed 170-bp endogenous genomic elements as reporter constructs, each paired with a measured activity label (log2_alpha_Endo). We randomly split 1,653 endogenous sequences into ten folds, using eight for training, one for validation, and one for testing. The probe was trained with a mean-squared-error loss using AdamW^32^ with default parameters and a weight decay of 1 × 10⁻² for up to 100 epochs, with early stopping after five epochs without improvement in validation loss. Sequences were augmented by reverse-complementation (p = 0.5) and random shifts of up to 15 bp (p = 0.5). Models were trained on a single NVIDIA H100 GPU.

Probe and optimization settings were selected by grid search, varying dropout^33^ (0.0, 0.1, 0.3), batch size (32, 64, 128), and learning rate (5e-4, 1e-3, 5e-3), yielding 27 configurations. The configuration with the lowest validation loss (dropout 0.3, batch size 64, learning rate 5e-3) was used in all subsequent analyses.

With hyperparameters fixed, one fold was held out as a common test set. The remaining nine folds were used to train a nine-model ensemble: each model used eight folds for training and one for validation and early stopping. Predictions were averaged across models. Performance on the held-out fold was assessed using Pearson and Spearman correlations and mean squared error (MSE).

To assess sensitivity to data partitioning, we repeated the procedure for 20 independent ten-fold splits, yielding 20 ensembles evaluated on distinct test sets. Chromosome-partitioned and random splits performed similarly, with the chromosome partitioned trained model used for downstream analysis.

### In silico saturation mutagenesis and AP-1 reliance

Single-nucleotide in silico saturation mutagenesis (ISM) was performed with the trained model on 310 native enhancers: 150 stable loci and 160 dynamic loci, with 40 dynamic loci from each age-up, age-down, exercise-up and exercise-down group. At every nucleotide position in each 170-bp insert, the native nucleotide was replaced by each of the other three bases. Positional importance was defined as the greatest absolute change in predicted activity between the native and the three alternative bases.

Canonical AP-1 sites were identified as TGA(C/G)TCA on either strand and an enhancer was classified as AP-1-reliant when at least one of its five highest-importance ISM positions overlapped an AP-1 footprint. Stable and dynamic enhancers were compared using two-sided Fisher’s exact tests, and the pooled site-count-adjusted association was assessed using a Cochran–Mantel–Haenszel test. Ninety-five per cent confidence intervals were estimated from 5,000 bootstrap resamples of enhancers.

Motif reliance across additional TF families was evaluated using full-length ISM profiles for all 1,653 native inserts. Eighteen candidate transcription-factor motifs were obtained from JASPAR 2024^34^ and scanned on both strands after converting position-weight matrices to log-odds scores using nucleotide frequencies pooled across the analysed inserts as background and a pseudocount of 0.5. Motif matches were called using an 80% relative-score threshold. For each enhancer and motif, reliance was quantified as the number of the 20 highest-importance ISM positions overlapping a motif footprint. To account for motif abundance, dynamic-minus-stable differences were standardized within strata containing 0, 1, 2, or at least 3 motif sites; strata lacking either enhancer class were omitted. Confidence intervals were estimated from 5,000 enhancer-level bootstrap resamples.

### Attribution-guided seqlet identification

To quantify the contribution of surrounding sequence, we applied the chromosome-held-out context probe to AP-1-motif-shuffled constructs. For each nucleotide position, the model calculated a gradient for each of the four possible bases, indicating how changes at that position affected the predicted score.

These four gradients were mean-centered^35^, and the value corresponding to the native nucleotide was retained as the observed-base contribution. Positional importance was defined as the absolute value of this observed-base contribution.

A 12-bp window was moved across each sequence and scored by the sum of positional importance. Windows overlapping the shuffled AP-1-motif were excluded. Within each enhancer, the three highest-scoring non-overlapping windows were designated high-attribution seqlets, and the three lowest-scoring non-overlapping eligible windows were designated low-attribution controls.

To select loci for follow-up seqlet analysis, loci were ranked using a reliability score constructed from the context probe. For locus i, reliability was ranked using:

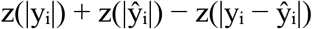

where yᵢ and ŷᵢ were the measured and predicted non-AP-1 context contrasts and the standard deviation was calculated across all 1,653 assayed loci. This score favors loci with large measured and predicted effects while penalizing disagreement between them. Sequences were required to show concordant measured and predicted effects across multiple replicates and were then ranked within enhancer class and effect direction. The 12 highest-ranking loci from each class were supplemented with prespecified model-failure and AP-1-responsive controls, producing a follow-up panel of 142 loci. Combinatorial analyses were restricted to 91 loci with complete sets of three high-attribution seqlets and three low-attribution controls.

### Direct seqlet perturbation

Each selected seqlet was disrupted independently by permuting the bases within its 12-bp window. Five unique, deterministic permutations were generated per seqlet. Mutants were scored with the chromosome-held-out context probe, and the perturbation effect was defined as the score of the unperturbed AP-1-motif-shuffled sequence minus the mutant score. Seqlet impact was summarized as the mean absolute score change across the five permutations.

For each locus, effects were averaged across the three high-attribution seqlets and separately across the three low-attribution controls and compared with a one-sided paired Wilcoxon signed-rank test. Association between continuous attribution magnitude and perturbation effect was assessed across evaluated seqlets using Spearman correlation. For comparisons between stable and dynamic enhancers, measurements were first reduced to one value per parent locus and then tested using a Kruskal–Wallis test.

### Combinatorial seqlet disruption and epistasis reliance

Among the 91 loci included in the combinatorial analysis, the three high-attribution seqlets were labelled A, B and C in descending order of measured perturbation effect; the three control seqlets were analyzed in parallel. For each locus, three matched local-permutation replicates were generated for each single, double and triple disruption (A, B, C, AB, AC, BC and ABC). All constructs were scored with the chromosome-held-out context probe.

### Pairwise epistasis for seqlets A and B was defined as

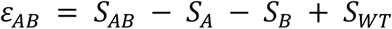

where S_WT_ was the score of the native AP-1-motif-shuffled sequence, S_A_ and S_B_ were the single-disruption scores, and S_AB_ was the double-disruption score. An additive interaction has ε = 0; negative values indicate a synergistic double-disruption loss, whereas positive values indicate buffering or redundancy. Interaction strength was summarized as |ε|. Values were averaged across permutations and seqlet pairs to obtain one high-attribution and one control value per locus, which were compared by a one-sided paired Wilcoxon signed-rank test.

Candidate interacting pairs were ranked by their mean |ε| across permutation replicates. A high-ranking pair was required to fall in the top decile of |ε|, exceed the corresponding within-locus control pair and retain the same interaction direction in at least two permutation replicates. Edge-to-edge seqlet distance and rescue behavior were retained as additional prioritization criteria.

Composition-preserving seqlet rescue

Rescue constructs were generated from the experimentally tested fully shuffled 170-bp sequence. Bases at one, two or all three selected seqlets were restored to the corresponding bases in the endogenous sequence. When restoration changed the nucleotide composition of the sequence, compensating base swaps were made outside the restored seqlets such that each construct retained the mononucleotide composition of the starting sequence.

For each rescue construct, we quantified how much of the activity difference between the fully shuffled and AP-1-motif-shuffled sequences was restored. This rescue fraction was defined as:

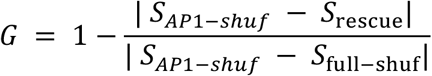

where S is the prediction score of each of the sequences. Values approaching 1 indicate restoration towards the AP-1-motif-shuffled score, whereas values at or below 0 indicate no rescue or movement away from the AP-1-motif-shuffled score. The rescue fraction was considered undefined when the predicted AP-1-motif-shuffled and fully shuffled baseline scores were exactly equal; one-, two- and three-seqlet rescue values were summarized for the high-attribution set. Triple-seqlet rescue was compared with the matched three-control rescue by a one-sided paired Wilcoxon signed-rank test. Enhancer-class comparisons used one triple-rescue value per locus and a Kruskal–Wallis test.

### Motif analysis

Native 170-bp inserts and local seqlet sequences were scanned on both strands using a 41-matrix JASPAR panel^34^. PWMs were converted to log_2_-odds scores relative to a uniform nucleotide background, with a pseudocount of 10⁻⁵. A motif hit was called when its score reached at least 80% of the matrix-specific range between the theoretical minimum and maximum. Native-insert occupancy was the number of qualifying hits. Dynamic and stable occupancy distributions were compared using two-sided Mann–Whitney U tests, followed by Benjamini–Hochberg correction across the 41 matrices.

For seqlet enrichment, raw PWM scores were evaluated in local 22-bp contexts centered on each seqlet. Scores were averaged separately across high-attribution and low-attribution seqlets within each locus and compared using one-sided paired Wilcoxon signed-rank tests. The prespecified family-level analysis comprised 15 motif families. Motif identities for individual interacting seqlets were assigned from the uncorrected PWM scan; seqlets failing the 80% threshold were labelled unresolved and accompanied by their highest-scoring candidate motif.

### Distributed sequence grammar, epistasis and rescue

Model-derived importance scores identified short 12-bp ‘seqlets’ that accounted for a substantial fraction of the predicted context effect. Across 142 high-confidence loci, perturbing high-attribution seqlets produced 2.58-fold larger effects than perturbing matched low-attribution regions (0.20 versus 0.08; paired P = 2.97 × 10^-21^), and attribution magnitude correlated with the subsequent perturbation effect (Spearman ρ = 0.571; P = 3.70 × 10^-74^). Pairwise disruption revealed greater absolute non-additivity among high-impact seqlets than among low-impact controls (0.06 versus 0.03; P = 4.26 × 10^-10^). Interaction direction was approximately balanced between generating a larger-than-additive loss of activity and a smaller-than-additive loss, indicating distributed and partially interchangeable sequence contributions rather than obligatory motif combinations.

Composition-preserving rescue provided a complementary test. Restoring one, two or three nominated seqlets within fully scrambled sequences recovered an average of 28.4%, 41.2% and 54.2% of the predicted context defect, respectively. Three-seqlet rescue was greater for high-impact than control seqlets (median 56.1% versus 31.9%; P = 2.25 × 10^-5^). Seqlet perturbation strength, epistasis and rescue did not differ significantly between stable and dynamic enhancers.

Beyond AP-1, motif analysis connected this distributed context code to candidate TFs. Count-matched ISM identified greater MEF2C reliance in dynamic enhancers (P = 5 × 10^−5^; **Figure 3P**). Consistent with the AT-rich nature of MEF2 motifs and the lower GC content of dynamic enhancers, complete native inserts from dynamic loci contained more MEF2C, TCF7L2 and GATA4 motif matches, whereas stable enhancers contained more NFATC3 and ETV4 matches (FDR ≤ 0.04). NKX2-5-family sequence was also enriched within high-impact sequence elements (FDR = 0.02). Motif annotation of the strongest interacting seqlets nominated MEF2C–TBX5, ETS–GATA-like, KLF4–NFATC3, AP-1/BATF–RUNX1 and TCF7L2–RUNX3 candidate pairs (**Figure 3L**). Broad AP-1-centred spacing screens did not reveal a reproducible grammar separating dynamic from stable enhancers across sequence backgrounds and model folds.

### Sparse detection of IL-1 ligands in snRNA-seq data

At the ligand level, *Il1a* and *Il1b* transcripts were detected in only a small fraction of nuclei (*Il1a*, 0– 2.9%; *Il1b*, 0–16.9%) across the examined cell populations, with the highest *Il1b* detection rate observed in MHC-II^hi^ macrophages (**Figure S12F**). Because expression was absent from most nuclei, condition-level estimates were driven by a small number of positive nuclei, limiting the precision of differential-expression tests. This limited detection is biologically plausible because *Il1b* exhibits low basal transcription and rapid stimulus-dependent induction in macrophages^22^, and technically plausible because snRNA-seq can underrepresent *Il1b* compared with single-cell RNA-seq^23^.

Consistent with this limitation, differential-expression testing identified no age-or exercise-associated ligand changes after multiple-testing correction (**Figure S12G**). Nevertheless, in a senescence-accelerated mouse model, RT-qPCR analysis of whole-heart tissue similarly reported increased *Il1b* expression with age and a reduction following resistance and aerobic training^24^.

### Macfib population distinct from technical doublets

The macfib population was not an artefact of residual droplet co-encapsulation following standard doublet removal for several reasons. Imaging identified macfib cells in heart tissue (**Figure S13D**), while flow cytometry confirmed CD45 expression in genetically labelled Pdgfra-lineage cells (**Figure S13E**). Furthermore, because each library pooled one male and one female animal, approximately half of all nuclear co-encapsulation events would be expected to be cross-sex. Macfib nuclei were assigned to a single sex as confidently as singlet macrophages. Moreover, macfib profiles fell outside the distribution of in silico macrophage–fibroblast mixtures, expressed myeloid markers at undiluted levels while carrying only part of the fibroblast program, and showed accessible chromatin absent from either parent population. Analogous macfibs/fibrocytes have been identified previously in the murine heart using lineage tagging and flow imaging independently of droplet-based methods^21^.

### AAV construction and treatment

To inhibit AP-1 DNA-binding activity in cardiomyocytes, we constructed an adeno-associated virus (AAV) expression plasmid: AAV9-cTnT-A-FOS-T2A-tdTomato. The dominant-negative AP-1 construct (A-FOS) was amplified by PCR using primers EW-FW-133 and EW-FW-134 (**Table S33**) with CMV500 A-FOS plasmid as the template (Addgene #33353). The tdTomato reporter cassette was amplified using primers EW-FW-135 and EW-FW-136 (**Table S33**) from the pHDR_DDX4-T2A-tdTomato-puroTK plasmid. The AAV9-cTnT backbone was amplified using primers EW-FW-131 and EW-FW-132 (**Table S33**) from the pENN-AAV-cTNT-PI-eGFP-WPRE-rBG plasmid (Addgene #105543). Purified PCR fragments were assembled using NEBuilder HiFi DNA Assembly Master Mix (NEB, cat. no. E2621L). The resulting plasmid was validated by whole plasmid sequencing. The plasmid was expanded and purified using the PureLink™ Fast Low Endotoxin Midi Plasmid Purification Kit (Thermo Fisher Scientific, cat. no. A35892). Endotoxin-free plasmid DNA was submitted to the Vectorology Facility, Children’s Medical Research Institute (CMRI), Sydney, NSW, Australia, for AAV9 packaging.

Mice aged 22–25 months were randomized to receive a single 100 μL (1 × 10¹¹ viral genomes per mouse) retro-orbital injection of AAV9-dnAP-1 (n = 7, comprising 3 males and 4 females) or AAV9-GFP control (n = 6, comprising 3 males and 3 females) under isoflurane anesthesia. All mice recovered rapidly following injection. Five weeks after injection, mice were euthanized and tissues were collected for downstream analyses.

### Physiological effects of IL-1 inhibitor in aged mice

Longitudinal echocardiography showed no treatment-associated change in systolic or chamber-geometry measures, including ejection fraction, fractional shortening, cardiac output, stroke volume, ventricular volumes, diameters and LV mass (**Figure S17G, Table S36**). Among diastolic indices, E′/A′ nominally diverged between groups (+0.087 anti-IL-1β versus −0.120 IgG; between-group difference = 0.207; Hedges’ g = 1.57; nominal P = 0.058; n = 3 and 4, respectively). Body weight also diverged (+2.57 g anti-IL-1β versus −1.08 g IgG; between-group difference = 3.65 g; Hedges’ g = 1.58; nominal P = 0.019; n = 6 and 5, respectively). Because weight loss in aged mice is itself a marker of systemic decline, and because E′/A′ and E/E′ are both load-dependent, this difference precludes attributing the diastolic finding to a direct cardiac action of IL-1β blockade. However, the diastolic finding accords with studies in aged animals and HFpEF models in which IL-1 signaling preferentially contributes to impaired relaxation and elevated filling pressure rather than reduced ejection fraction^36–38^.

## Notes

### Competing Interest Statement

The authors have declared no competing interest.

